# Mitochondrial GTP Metabolism Regulates Reproductive Aging

**DOI:** 10.1101/2023.04.02.535296

**Authors:** Yi-Tang Lee, Marzia Savini, Tao Chen, Jin Yang, Qian Zhao, Lang Ding, Shihong Max Gao, Mumine Senturk, Jessica Sowa, Jue D. Wang, Meng C. Wang

## Abstract

Healthy mitochondria are critical for reproduction. During aging, both reproductive fitness and mitochondrial homeostasis decline. Mitochondrial metabolism and dynamics are key factors in supporting mitochondrial homeostasis. However, how they are coupled to control reproductive health remains unclear. We report that mitochondrial GTP metabolism acts through mitochondrial dynamics factors to regulate reproductive aging. We discovered that germline-only inactivation of GTP- but not ATP-specific succinyl-CoA synthetase (SCS), promotes reproductive longevity in *Caenorhabditis elegans.* We further revealed an age-associated increase in mitochondrial clustering surrounding oocyte nuclei, which is attenuated by the GTP-specific SCS inactivation. Germline-only induction of mitochondrial fission factors sufficiently promotes mitochondrial dispersion and reproductive longevity. Moreover, we discovered that bacterial inputs affect mitochondrial GTP and dynamics factors to modulate reproductive aging. These results demonstrate the significance of mitochondrial GTP metabolism in regulating oocyte mitochondrial homeostasis and reproductive longevity and reveal mitochondrial fission induction as an effective strategy to improve reproductive health.

## INTRODUCTION

As one of the earliest signs of age-associated decline, reproductive senescence has a strong impact on society due to the trend of increased average maternal age at first birth^1^. Aged women exhibit decreased fertility and increased rates of birth defects and miscarriages^2^. It is estimated that fertility decline occurs on an average of 10 years prior to menopause, and an age-associated decrease in oocyte quality is the major cause for this decline^3^. Diverse factors can influence oocyte quality, and one of the main contributors is mitochondrial activity^4^. Oocytes have the largest number of mitochondria among all the cells in an organism^5^. Changes in mitochondrial ATP production, membrane potential, and DNA copy numbers have been reported to influence oocyte development, maturation and fertility^4, 6–8^. Meanwhile, mitochondria exhibit highly dynamic morphology and constantly undergo organellar fission and fusion, leading to changes in their shape, size, and distribution^9^. Specific types of protein machinery are required to maintain mitochondrial fission-fusion dynamics, including the mitochondrial fission GTPase DRP1, mitochondrial outer membrane fusion GTPases MFN1 and MFN2, and mitochondrial inner membrane fusion GTPase OPA1^9^. These regulators of mitochondrial dynamics also modulate mitochondrial distribution within the cell, especially in the oocyte. In mice with Drp1 knockout, the oocyte mitochondrial network is aggregated toward the perinuclear region^10^. Similarly, in mouse oocytes overexpressing Mfn1 or Mfn2, the mitochondrial network also exhibits perinuclear accumulation without increasing tubular elongation^11^. Mitochondrial dynamics factors have been linked with oocyte development and maturation^10, 12, 13^. Drp1 knockout in oocytes results in abnormal follicular maturation, defective meiotic resumption, and fertility decline in mice^10^. Oocyte-specific knockout of mouse Mfn1 causes defective folliculogenesis and apoptotic cell loss, leading to complete infertility^12, 13^. These findings indicate the importance of mitochondrial dynamics factors in regulating oocyte quality during development.

On the other hand, in *C. elegans*, mitochondrial dynamics factors have been linked with the regulation of somatic aging. Selectively overexpressing the *C. elegans* DRP1 homolog *drp-1* in the intestine prolongs lifespan^14^, and whole-body knockout of *drp-1* together with *fzo-1*, the *C. elegans* MFN homolog, leads to lifespan extension in *C. elegans*^15^. In addition, the lifespan-extending effect associated with DRP1 overexpression has been also reported in *Drosophila*^16^. Besides being a well-established model organism for studying somatic aging, *C. elegans* share similarities with humans regarding reproductive aging. Like in humans, the reproductive time window in *C. elegans* takes approximately one-third of its total lifespan, and with the increase of age, both oocyte quality and fertility decline^17, 18^. Not only genetic factors but also environmental cues including bacterial species contained in the diet are known to regulate reproductive aging in *C. elegans*^19^. Upon their exposure to different bacteria, worms exhibit a distinct reproductive lifespan (RLS), which can be further modified by genetic manipulations^19^. Moreover, through a full-genome RNA interference (RNAi) screen, we have identified several mitochondrial genes as regulators of reproductive aging in *C. elegans*^20^, which includes two subunits of Succinyl-CoA Synthetase (SCS).

SCS is a key mitochondrial enzyme in the TCA cycle converting succinyl-CoA to succinate with a production of GTP or ATP^21^. A functional SCS enzyme comprises one alpha subunit and one beta subunit. Two interchangeable beta subunits of SCS determine the GTP/ATP specificity by forming a complex with the constant alpha subunit^22, 23^. Results from immunoblotting analyses in mammals reveal that SCS beta subunits exhibit heterogeneous expression patterns across different tissues^24^. In humans, kidney and liver have a relatively high level of the GTP-specific beta subunit in comparison to the high level of the ATP-specific beta subunit in heart, testis, and brain^24^. In mitochondria, ATP is primarily synthesized through oxidative phosphorylation, while GTP is predominantly generated by the GTP-specific isoform of SCS in the TCA cycle. Thus, it is predicted that the GTP-specific beta subunit of SCS acts as a metabolic sensor of the TCA-cycle flux and couples it with glucose homeostasis^25^. Consistently, the GTP-specific beta subunit of SCS in pancreatic beta cells is essential for glucose-sensing and insulin secretion^25, 26^.

In this study, we discovered that the GTP-specific SCS in the germline regulates reproductive aging through tuning mitochondrial positioning in the oocyte, and that increasing mitochondrial fission selectively in the germline prevents age-associated perinuclear accumulation of mitochondria in the oocyte and promotes reproductive longevity in *C. elegans*. We found that knockdown of *sucg-1,* encoding the GTP-specific beta subunit of SCS, extends RLS, improves late fertility, and attenuates an age-associated increase in oocyte mitochondrial clustering around the nucleus. Germline-specific depletion of the DRP-1 protein suppressed the reproductive-longevity-promoting effect caused by the *sucg-1* knockdown. Conversely, germline-specific overexpression of *drp-1* or germline-specific knockdown of *eat-3,* the *C. elegans* OPA1 homolog, was sufficient to promote reproductive longevity and attenuate the age-associated perinuclear accumulation of oocyte mitochondria. Furthermore, we found that the regulation of reproductive aging by the GTP-specific SCS and mitochondrial dynamics factors responds to the level of vitamin B12 in bacteria. Taken together, our findings reveal a previously unknown function of mitochondrial GTP metabolism in the germline and its significance in the regulation of mitochondrial homeostasis and oocyte quality during aging. This work also suggests that fine-tuning mitochondrial distribution selectively in the reproductive system through either genetic manipulation or dietary intervention is an effective strategy to promote reproductive longevity.

## RESULTS

### GTP-specific SCS regulates reproductive aging

In the genome-wide RNAi screen searching for regulators of reproductive longevity, *sucl-2* and *sucg-1*, which encode the alpha and beta subunit of SCS respectively, were identified^20^. SCS catalyzes an essential step in the TCA cycle converting succinyl-CoA to succinate (**Figure 1A**). To further understand the role SCS plays in regulating reproductive longevity, we first performed longitudinal studies and found that inactivating either *sucl-2* or *sucg-1* by RNAi results in not only RLS extension, but also improved fertility in aged hermaphrodites (late fertility) (**Figures 1B-D, Supplementary Table 1**). As the age of control hermaphrodites increased from 1-day-old to 7-day-old and 9-day-old, the percentage of individuals capable of reproducing was decreased from 100% to less than 50% and 30% respectively when they were mated with 2-day-old young males (**Figure 1D**). With *sucg-1* or *sucl-2* RNAi knockdown, the percentage of the aged hermaphrodites capable of reproducing was increased to more than 70% or 90% at day 7, and more than 50% or 70% at day 9, respectively (**Figure 1D**).

**Figure 1.**
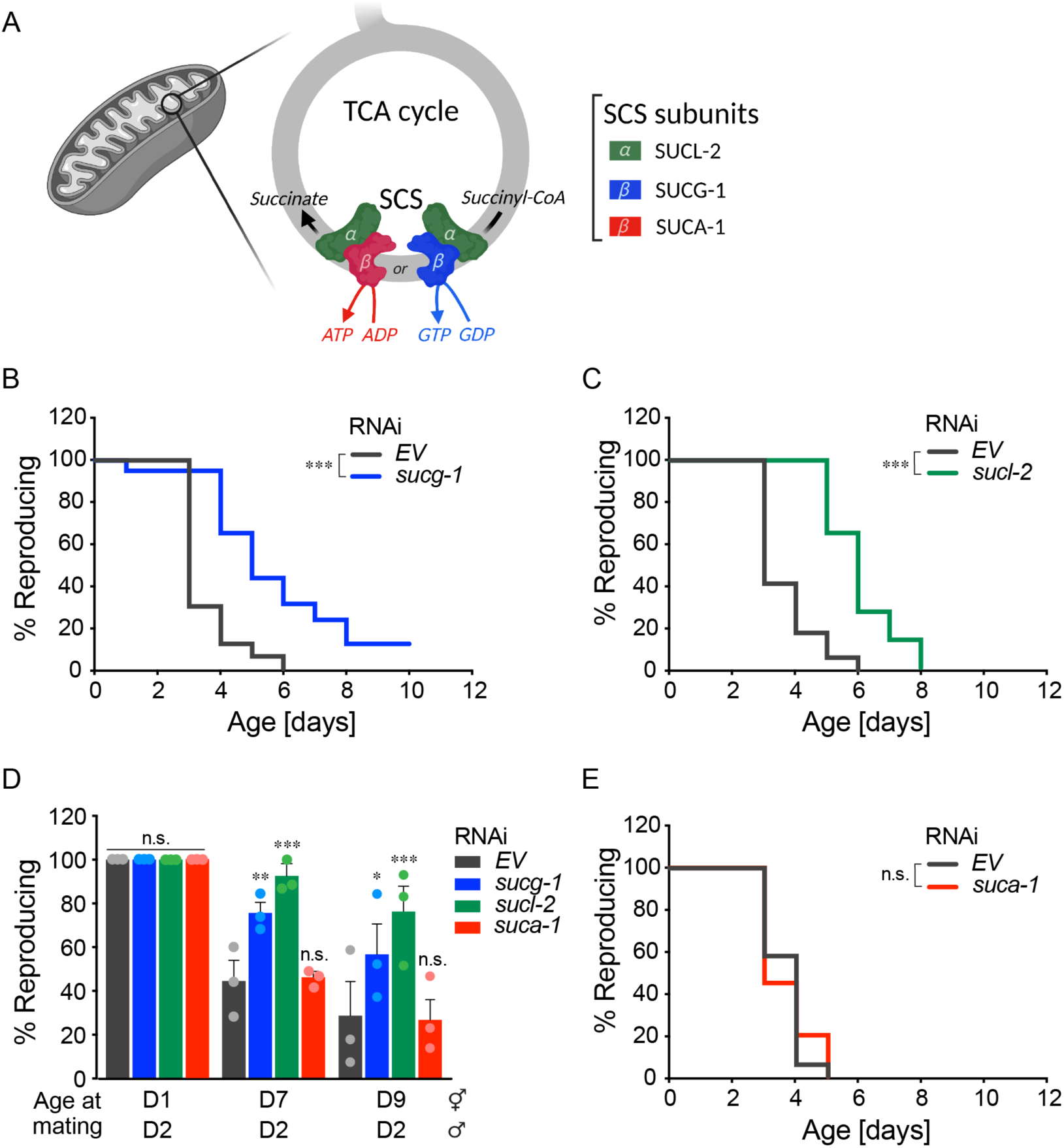
GTP-specific Succinyl-CoA Synthetase (SCS) regulates reproductive aging. (A) A diagram of SCS enzymatic function and its ATP or GTP specificity. (B) Wild-type (WT) worms subjected to *sucg-1* RNA interference (RNAi) have a significantly longer reproductive lifespan (RLS) than those subjected to the empty vector (EV) control. (C) WT worms subjected to *sucl-2* RNAi have a longer RLS than those subjected to the EV control. (D) Day 7 and 9 WT hermaphrodites subjected to *sucg-1* or *sucl-2* but not *suca-1* RNAi show higher rates of reproduction than those subjected to the EV control, when mated with day-2-old males. (E) WT worms subjected to *suca-1* RNAi show no significant differences in RLS compared to those subjected to the EV control. (B, C, E) n.s. *p > 0.05,* *** *p < 0.001* by log-rank test; n = 3 biological independent replicates, ∼20 worms per replicate, see Supplementary Table 1 for full RLS Data. (D) Error bars represent mean ± s.e.m., n = 3 biologically independent samples, n.s. *p > 0.05*, * *p < 0.05,* ** *p < 0.01,* *** *p < 0.001* by Fisher’s exact test adjusted with the Holm–Bonferroni method for multiple comparisons, ∼20 worms per replicate.

It is known that SCS forms either an ATP- or GTP-specific heterodimer enzyme complex, which produces ATP or GTP alongside the conversion of succinyl-CoA to succinate, respectively (**Figure 1A**). This specificity for ATP or GTP production relies on distinct beta subunits, but not on the constant alpha subunit^22^. In *C. elegans*, *sucg-1* encodes the GTP-specific beta subunit and *suca-1* encodes the ATP-specific beta subunit. We found that unlike *sucg-1,* the RNAi knockdown of *suca-1* shows no RLS extension or improvement of late fertility (**Figures 1D and 1E, Supplementary Table 1**). These results suggest that the SCS complex formed by the alpha subunit encoded by *sucl-2* and the beta subunit encoded by *sucg-1* is specifically involved in regulating RLS and late fertility. Given that SUCG-1 is responsible for converting GDP to GTP in mitochondria, these results indicate a possible role of mitochondrial GTP metabolism in modulating reproductive aging.

### GTP-specific SCS functions in the germline to regulate mitochondria and reproductive aging

To understand how the GTP-specific SCS regulates reproductive aging, we first examined the expression pattern of *sucg-1* using a CRISPR knock-in line in which the endogenous SUCG-1 was tagged with eGFP at the C terminus. Using this line, we detected SUCG-1::eGFP expression predominantly in the germline, with weaker signals in the pharynx, intestine, hypodermis, and muscle (**Figure 2A**). Using CRISPR knock-in, the endogenous SUCA-1 was also tagged with eGFP at the C terminus, which revealed the predominant expression of *suca-1* in the pharynx, neuron, intestine, hypodermis, and muscle but a very weak signal in the germline (**Figure S1A**). Moreover, we found that the GFP intensity in the germline of the SUCG-1::eGFP worms is increased at day-5 adulthood when compared to day-1 adulthood (**Figures 2B and 2C**), suggesting an elevation of germline SUCG-1 levels with aging. These findings suggest that mitochondrial SUCG-1 may function in the germline to regulate reproductive aging cell-autonomously.

**Figure 2.**
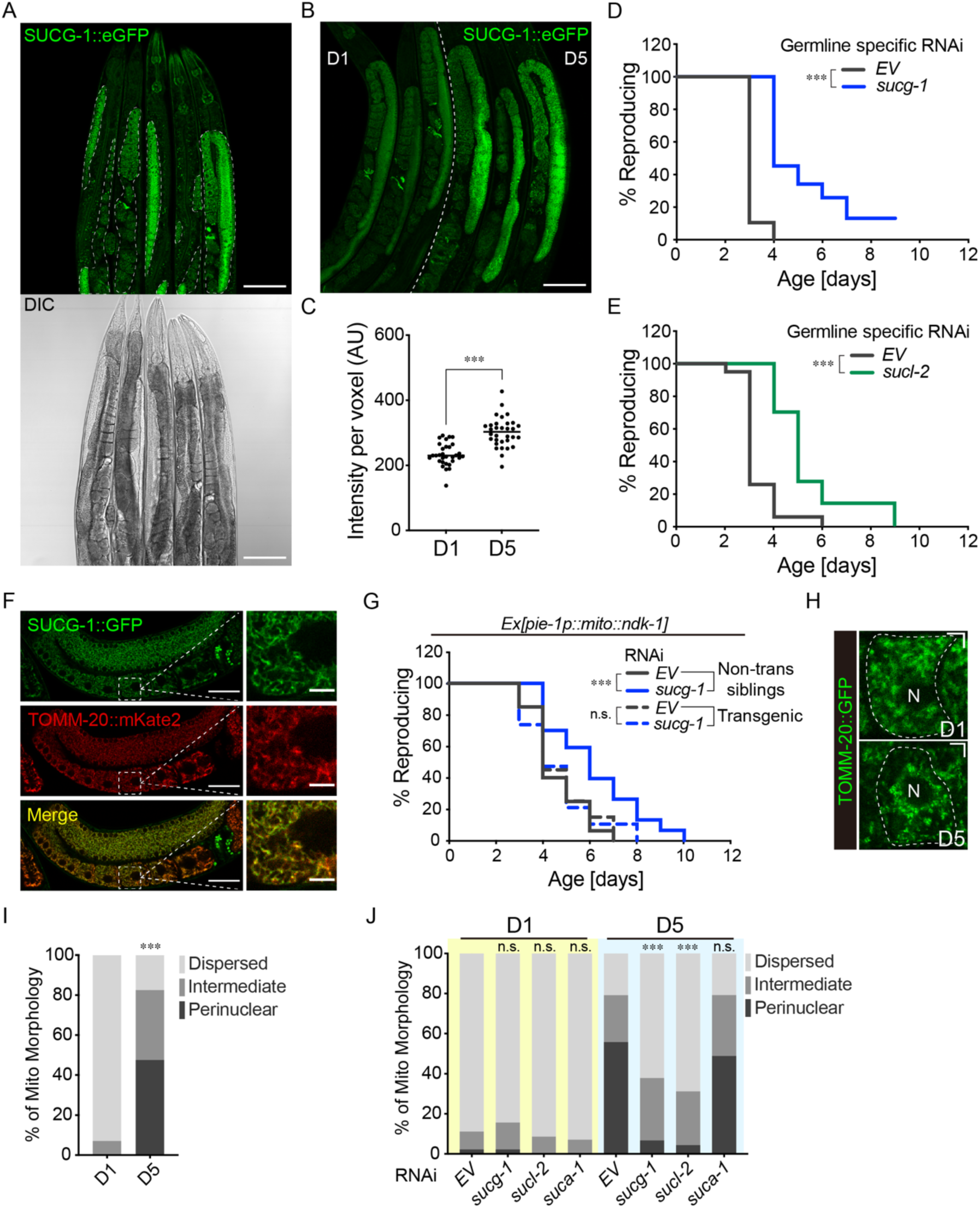
GTP-specific SCS functions in the germline to regulate oocyte mitochondria during reproductive aging. (A) Confocal imaging of the SUCG-1::eGFP knock-in line, in which the endogenous *sucg-1* is tagged with *egfp,* reveals its predominant expression in the germline but weak expression in the intestine, pharynx, muscle, hypodermis and neurons (Scale bar: 100μm; Dashed white line: germline). (B) The SUCG-1::eGFP level in the germline is increased at day 5 compared to day 1 (Scale bar: 100μm). (C) Quantification of SUCG-1::eGFP level in the germline at day 1 and day 5. (D) Germline-specific RNAi inactivation of *sucg-1* extends RLS. (E) Germline-specific RNAi inactivation of *sucl-2* extends RLS. (F) SUCG-1::eGFP colocalizes with mitochondrial TOMM-20::mKate2 in the germline (Scale bar: 30μm for the images with lower magnification; 5μm for the images with higher magnification). (G) Overexpression of mitochondria-targeted *ndk-1*(*mito::ndk-1*) in the germline suppressed the RLS extension caused by *sucg-1* RNAi knockdown. (H) Representative images show that oocyte mitochondria are largely dispersed at day 1 while increasing perinuclear distribution at day 5 (Scale bar: 5μm; Dashed white line: oocyte outline; N: nucleus). (I) Perinuclear clustering of oocyte mitochondria is increased from day 1 to day 5. (J) The increase in perinuclear distribution of oocyte mitochondria at day 5 is suppressed by *sucg-1* or *sucl-2*, but not *suca-1* RNAi knockdown. (C) *** *p < 0.001* by Student’s t-test; n = 31 (day 1), n = 32 (day 5). (D, E, G) *** *p < 0.001* by log-rank test; n = 4 (D) or 3 (E, G) biological independent replicates, ∼20 worms per replicate, see Supplementary Table 1 for full RLS Data. (I) n = 43 (day 1) and n = 40 for (day 5); *** *p < 0.001* by Chi-squared test. (J) n= 45 (EV, D1), n = 45 (*sucg-1*, D1), n = 45 (*sucl-2*, D1), n = 45 (*suca-1*, D1), n = 42 (EV, D5), n = 45 (*sucg-1*, D5), n = 44 (*sucl-2*, D5) and n = 48 (*suca-1*, D5); RNAi vs. EV, n.s. *p > 0.05,* *** *p < 0.001* by Chi-squared test adjusted with the Holm–Bonferroni method for multiple comparisons.

To further confirm this cell-autonomous regulation, we utilized a tissue-specific RNAi strain, in which the expression of the RNAi-induced silencing complex component RDE-1 is restored specifically in the germline of the *rde-1* null mutant^29^. We knocked down either *sucg-1* or *sucl-2* by RNAi selectively in the germline and found that germline-specific knockdown of *sucg-1* extends RLS compared to control worms treated with the empty vector (**Figure 2D, Supplementary Table 1**). Germline-specific knockdown of *sucl-2* led to similar RLS extending effects (**Figure 2E, Supplementary Table 1**). These results suggest that *sucg-1* and *sucl-2* act in the germline to regulate reproductive longevity. We also measured the progeny number in those worms with extended RLS and observed 7% or 11% reduction associated with the *sucg-1* or *sucl-2* germline-specific RNAi knockdown, respectively (**Figures S1B and S1C**). The decrease in the progeny number has been previously observed in other interventions leading to RLS extension, such as the loss-of-function mutant of *daf-2*, *eat-2,* and *sma-2*^18, 30–32^. In addition, the *daf-2* and the *eat-2* mutants not only prolong RLS but also extend lifespan^33, 34^. We found that the whole-body RNAi knockdown of either *sucg-1* or *sucl-2* leads to a mild lifespan extension (∼15%), but the *suca-1* knockdown does not affect lifespan (**Figure S1D, Supplementary Table 2**). Upon germline-specific RNAi knockdown, the result was similar except that *sucg-1* showed no lifespan extension in one out of three trials, and *suca-1* slightly shortened lifespan in one out of three trials (**Figure S1E, Supplementary Table 2**).

Both GTP- and ATP-specific SCS isoforms function in the TCA cycle to catalyze succinate production from succinyl-CoA, and their loss both lead to increased succinyl-CoA and decreased succinate levels. However, the RNAi knockdown of *suca-1* and *sucg-1* exert distinctive effects on RLS, suggesting that the change in either succinate or succinyl-CoA level is unlikely linked with the regulation of reproductive aging. In supporting of this idea, we found that dietary supplementation of sodium succinate or succinic acid does not alter RLS (**Figure S1F, Supplementary Table 1**). Additionally, germline-specific knockdown of *ogdh-1*, which encodes a subunit of α-ketoglutarate dehydrogenase (upstream of SCS), led to sterile phenotype in worms. Meanwhile, germline-specific knockdown of *mev-1* or *sdhb-1* encoding subunits of succinate dehydrogenase (downstream of SCS) resulted in a very short reproductive time window (**Figure S1G, Supplementary Table 1**). Together, these results demonstrate that GTP-specific SCS functions in the germline to regulate reproductive aging, and this regulatory effect is not associated with changes in succinate or succinyl-CoA levels.

We further confirmed mitochondrial localization of SUCG-1 by crossing the SUCG-1::eGFP line with the transgenic strain that expresses mKate2 tagged TOMM-20 on the outer mitochondrial membrane in the germline^35^. We observed co-localization between SUCG-1::eGFP and TOMM-20::mKate2 (**Figure 2F**). Thus, like its human homolog SUCLG1, SUCG-1 resides in mitochondria. Next, we tested whether SUCG-1 regulates reproductive longevity through affecting mitochondrial GTP (mtGTP) levels in the germline. To this end, we have made a transgenic strain expressing mitochondrial-matrix-targeting-sequence tagged *ndk-1* specifically in the germline. *ndk-1* encodes the nucleoside diphosphate kinase that catalyzes the synthesis of GTP from ATP, and thus its expression would increase GTP levels^36^. We found that *ndk-1* overexpression in germline mitochondria is sufficient to suppress the RLS extension caused by *sucg-1* knockdown (**Figure 2G, Supplementary Table 1**), suggesting that GTP-specific SCS regulates reproductive aging through modulating mtGTP levels in the germline.

Next, to test whether the loss of SCS affects germline mitochondrial homeostasis, we utilized the transgenic strain expressing GFP tagged TOMM-20 in the germline^35^ and imaged mitochondrial morphology at day 1 and day 5 of adulthood. We found that mitochondrial fragmentation and tubulation morphology exhibits high variations between individuals of the same genotype, which prevented us to draw an explicit conclusion. On the other hand, we observed that the mitochondrial network of oocytes increases perinuclear distribution in day 5 aged worms, while being largely dispersed in day 1 young worms (**Figure 2H**). We wrote an imaging analysis script to quantify mitochondrial distribution. This method first divided oocyte cells into five rings, with the first ring being the closest and the fifth ring being the most distant from the nucleus, and then calculated the percentage of mitochondrial GFP signal intensity in each ring among all five (**Figure S2A**). Next, based on the percentage of the signal intensity within ring 1, the images of oocyte mitochondria were categorized into three categories – dispersed, intermediate, and perinuclear. The quantification results using this method showed that mitochondrial GFP signals are evenly distributed throughout the five rings in the oocyte of day 1 worms (**Figure S2B**), while in the oocyte of day 5 worms, the percentage of the mitochondrial GFP signal derived from ring 1 is increased (**Figures S2B and S2C**). Further categorization analysis revealed that the perinuclear distribution of oocyte mitochondria is increased in day 5 worms (**Figure 2I**). To test whether this change in mitochondrial distribution is associated with a decrease in mitochondrial content, we dissected germline and measured mitochondrial DNA (mtDNA) levels using quantitative PCR (qPCR). The result showed that the mtDNA level is 60% higher in the germline of day 5 aged worms than that in day 1 young worms (**Figure S2D**), indicating that the age-associated perinuclear accumulation of oocyte mitochondria is unlikely due to a decline in mitochondrial numbers.

Interestingly, we found that RNAi knockdown of *sucg-1* or *sucl-2* suppresses the age-associated increase in mitochondrial clustering around the nucleus, while RNAi knockdown of *suca-1* shows no such effect (**Figures 2J and S3A**), which are consistent with their effects on RLS and late fertility (**Figures 1B-E**). Furthermore, the qPCR result showed that *sucg-1*, *sucl-2*, or *suca-1* germline-specific RNAi knockdown does not affect the germline mtDNA level at day 1 but leads to ∼30% increase at day 5 (**Figure S2E**). Thus, the loss of either SCS isoform increases mitochondrial content in the germline with aging, which is not specific to *sucg-1* knockdown and thus unlikely related with its effect on oocyte mitochondria positioning. Together, we found that the GTP-specific SCS works specifically in the germline to regulate oocyte mitochondrial distribution during reproductive aging.

### Mitochondrial fission drives reproductive longevity

It is known that mitochondrial dynamics and distribution are both controlled by the dynamin family of GTPases that mediate the balance between organellar fusion and fission^9^. To determine whether the key dynamin family of large GTPases regulate reproductive aging, we examined EAT-3, FZO-1 and DRP-1, which are *C. elegans* homologs of human OPA1, MFN1/2 and DNM1L respectively^37^. Both EAT-3 and FZO-1 control mitochondrial fusion, with EAT-3 driving inner mitochondrial membrane fusion while FZO-1 being responsible for the fusion of the outer mitochondrial membrane (**Figure 3A**)^38, 39^. We found that germline-specific RNAi knockdown of *eat-3* increases RLS and late fertility (**Figures 3B and 3C, Supplementary Table 1**). Meanwhile, knocking down *fzo-1* selectively in the germline did not affect late fertility (**Figure 3C**), and only showed slight RLS extension (11.5%) in one out of three trials but having no effect in the other two (**Figure 3D, Supplementary Table 1**). Thus, in the germline, EAT-3-mediated inner mitochondrial membrane fusion is involved in regulating reproductive aging. The *eat-3* mutant was originally discovered showing abnormal pharyngeal pumping and food intake, like the *eat-2* mutant^40^. The *eat-2* mutant is known to slow down reproductive aging as a result of caloric restriction^33, 40^. To test whether the effect of *eat-3* on reproductive aging is also due to a reduction in food intake, we have measured the pharyngeal pumping rate and the body size in worms with the germline-specific *eat-3* RNAi knockdown. We found that worms with germline-specific *eat-3* RNAi knockdown show a pharyngeal pumping rate and body size indistinguishable from the controls (**Figures S4A-C**), suggesting that the RLS extension does not result from caloric restriction.

**Figure 3.**
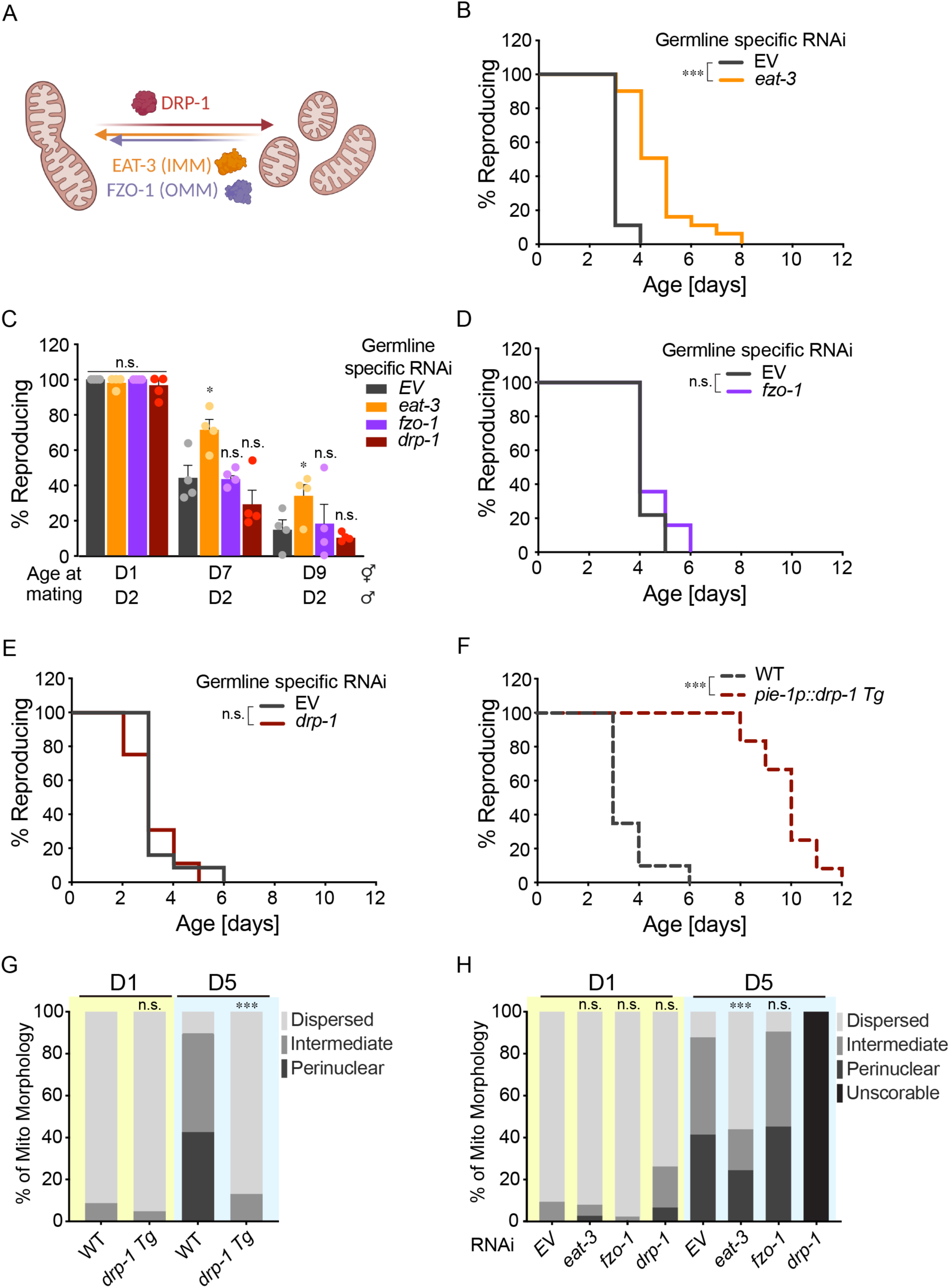
Mitochondrial dynamics factors regulate reproductive longevity. (A) A diagram showing regulation of mitochondrial dynamics by GTPase DRP-1, FZO-1 and EAT-3 (IMM: Inner mitochondrial membrane; OMM: Outer mitochondrial membrane). (B) Germline-specific RNAi inactivation of *eat-3* extends RLS. (C) Day 7 and 9 aged hermaphrodites subjected to germline-specific *eat-3* RNAi have a higher rate of reproduction than those subjected to the EV control when mated with day-2-old young males, while germline-specific RNAi inactivation of *fzo-1* or *drp-1* RNAi does not affect the rate of reproduction at all ages. (D) Germline-specific RNAi inactivation of *fzo-1* does not affect RLS. (E) Germline-specific RNAi inactivation of *drp-1* does not affect RLS. (F) Germline-specific overexpression of *drp-1* prolongs RLS. (G) The perinuclear clustering of oocyte mitochondria at day 5 is decreased in the transgenic strain with germline-specific *drp-1* overexpression. (H) The increase in the perinuclear distribution of oocyte mitochondria at day 5 is decreased upon *eat-3* but not *fzo-1* RNAi knockdown. The distribution of oocyte mitochondria is not scorable in day 5 aged worms subjected to *drp-1* RNAi knockdown due to distorted germline. (B, D, E, F) n.s. *p > 0.05,* *** *p < 0.001* by log-rank test; n = 3 biological independent replicates, ∼20 worms per replicate, see Supplementary Table 1 for full RLS Data. (C) Error bars represent mean ± s.e.m., n = 4 biologically independent samples, n.s. *p > 0.05*, * *p < 0.05* by Fisher’s exact test adjusted with the Holm–Bonferroni method for multiple comparisons, ∼15 worms per replicate. (G) n= 46 (WT, D1), n = 42 (*drp-1 OE*, D1), n = 40 (WT, D5), n = 46 (*drp-1 OE*, D5); WT vs. *drp-1* OE, n.s. *p > 0.05,* *** *p < 0.001* by Chi-squared test. (H) n= 43 (EV, D1), n = 38 (*eat-3*, D1), n = 40 (*fzo-1*, D1), n = 46 (*drp-1*, D1), n = 41 (EV, D5), n = 41 (*eat-3*, D5), n = 42 (*fzo-1*, D5); RNAi vs. EV, n.s. *p > 0.05,* *** *p < 0.001* by Chi-squared test adjusted with the Holm–Bonferroni method for multiple comparisons.

In contrast to EAT-3 and FZO-1, DRP-1 drives mitochondrial fission (**Figure 3A**)^41, 42^. When we knocked down *drp-1* by RNAi selectively in the germline, we found that RLS either remains unchanged (in two replicates) or is slightly decreased (in one replicate), and late fertility is not altered in these worms (**Figures 3C and 3E, Supplementary Table 1**). Conversely, when we overexpressed *drp-1* selectively in the germline, the transgenic worms showed an extremely long RLS compared to control worms (**Figure 3F, Supplementary Table 1**). Together, these results show that increasing mitochondrial fission factors and decreasing inner mitochondrial fusion factors in the germline are both sufficient to promote reproductive longevity.

Next, we examined whether these mitochondrial dynamics factors regulate oocyte mitochondrial distribution. We found that in the *drp-1* germline-specific overexpression transgenic strain, the age-associated perinuclear accumulation of oocyte mitochondria is greatly suppressed in day 5 worms (**Figures 3G and S3B**). RNAi knockdown of *eat-3* also decreased the perinuclear accumulation of oocyte mitochondria in day 5 aged worms (**Figures 3H and S3A**); however. RNAi knockdown of *fzo-1* did not affect oocyte mitochondrial distribution in either day 1 or day 5 worms (**Figures 3H and S3A**). Upon *drp-1* RNAi knockdown, we observed an increase in the perinuclear distribution of oocyte mitochondria in day 1 young worms, which however did not reach statistical significance (**Figures 3H and S3A**). In day 5 aged worms, *drp-1* RNAi knockdown caused disruption in oocyte organization, and mitochondrial morphology became largely unscorable (**Figures 3H and S3C**). In few oocytes that still have recognizable cell boundaries, we observed one-sided perinuclear aggregation of mitochondria (**Figure S3A**). These results suggest that mitochondrial dynamics factors modulate mitochondrial distribution in the oocyte, which correlates with their regulatory effects on reproductive aging.

### GTP-specific SCS regulates reproductive aging through tuning mitochondrial distribution

We then asked whether the change in mitochondrial distribution is responsible for the reproductive longevity-promoting effect conferred by the *sucg-1* knockdown. To answer this question, we have utilized an auxin-inducible degron (AID) system to deplete the DRP-1 protein specifically in the germline upon the auxin treatment (**Figure 4A**). We first generated a CRISPR knock-in line *(gfp::degron::drp-1)* in which the endogenous DRP-1 is tagged with GFP and degron at the N terminus^43^. This line was next crossed with the single-copy integrated transgenic strain where the auxin-inducible F-box protein TIR1 in the E3 ubiquitin ligase complex is selectively expressed in the germline (*sun-1p::TIR1::mRuby*)^44^. Using this AID system, the auxin administration led to TIR1-mediated degradation of the degron-tagged DRP-1 protein in the germline but not in other tissues (**Figure 4B**). We found that the auxin-induced DRP-1 depletion in the germline causes no significant change in RLS (**Figure 4C, Supplementary Table 1**), recapitulating the finding from germline-specific RNAi knockdown of *drp-1* (**Figure 3E**). More importantly, although the germline-specific DRP-1 depletion does not affect RLS on its own, it fully suppressed the RLS extension in the *sucg-1* RNAi knockdown worms (**Figure 4D, Supplementary Table 1**).

**Figure 4.**
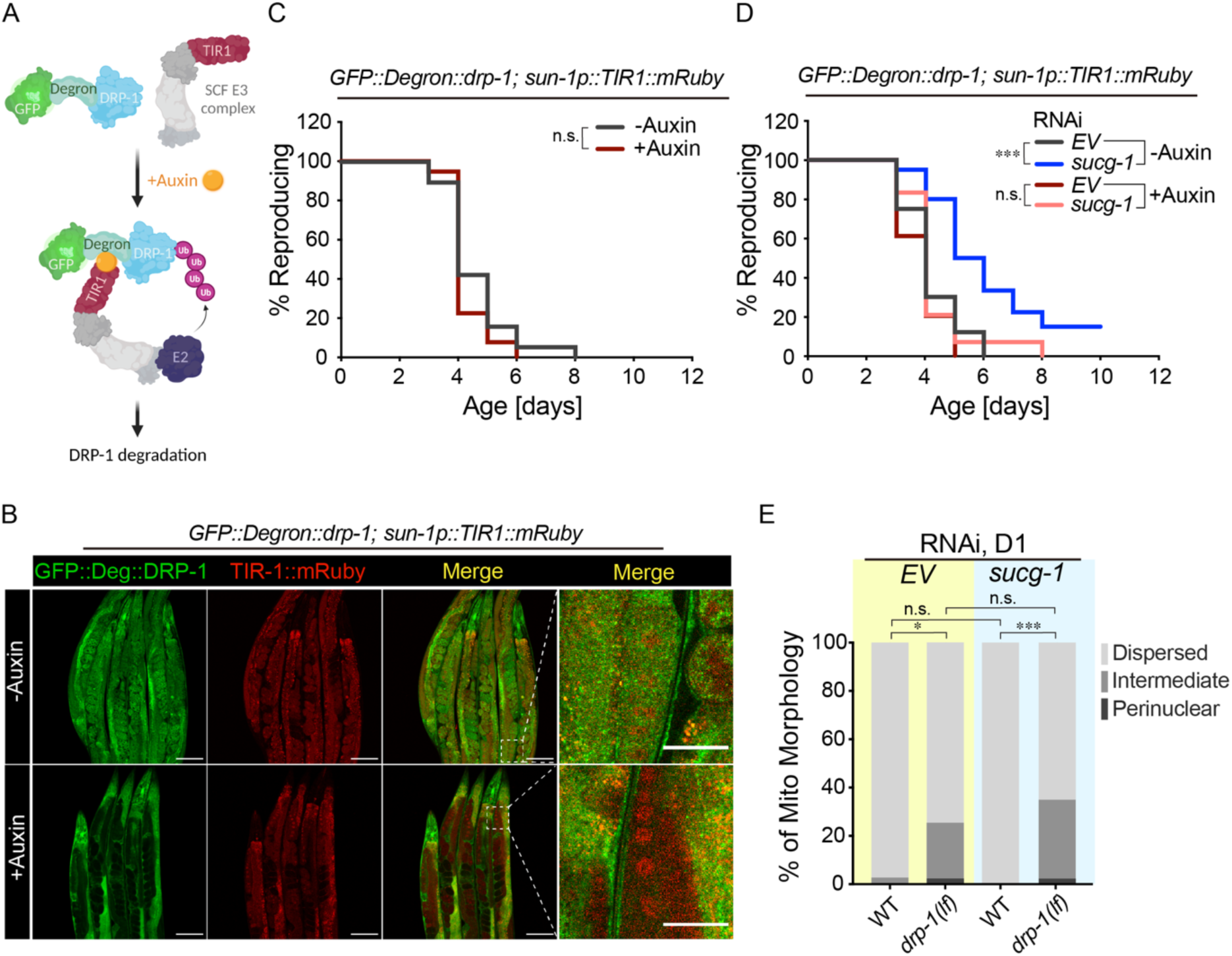
GTP-specific SCS regulates reproductive aging through mitochondrial dynamics factors. (A) A diagram demonstrating auxin-induced degradation of endogenous DRP-1 tagged with GFP and Degron. (B) Confocal imaging of GFP shows that the endogenous DRP-1 protein is specifically depleted in the germline upon the auxin treatment (Scale bar: 100μm for the images with lower magnification; 30μm for the images with higher magnification). (C) Auxin-induced germline-specific depletion of DRP-1 does not affect RLS. (D) Auxin-induced germline-specific depletion of DRP-1 abrogates the RLS extension caused by *sucg-1* RNAi. (E) The *drp-1* loss-of-function mutant increases the perinuclear clustering of oocyte mitochondria at day 1, which is not suppressed by *sucg-1* RNAi knockdown. (C, D) n.s. *p > 0.05,* *** *p < 0.001* by log-rank test; n = 3 biological independent replicates, ∼20 worms per replicate, see Supplementary Table 1 for full RLS Data. (E) n= 38 (WT, EV RNAi, D1), n = 41 (*drp-1(tm1108),* EV RNAi, D1), n = 41 (WT, *sucg-1* RNAi, D1), n = 46 (*drp-1(tm1108), sucg-1* RNAi, D1); RNAi vs EV, * *p < 0.05,* *** *p < 0.001 by* Chi-squared test adjusted with the Holm–Bonferroni method for multiple comparisons.

Furthermore, the *drp-1* loss-of-function mutant increased perinuclear clustering of oocyte mitochondria at day 1, and *sucg-1* RNAi knockdown failed to suppress this increase (**Figures 4E and S3D**), which suggests that DRP-1 is required for the loss of SUCG-1 to drive oocyte mitochondrial dispersion. Therefore, mitochondrial GTP metabolism can regulate reproductive longevity by affecting mitochondrial positioning in the germline through a DRP-1-mediated mechanism.

### GTP-specific SCS regulates reproductive aging in response to bacterial inputs

To further confirm the difference between *sucg-1* and *suca-1* in regulating reproductive aging, we generated CRISPR knockout lines for both (**Figure S5A**). *suca-1* knockout worms were phenotypically wild type, and similarly to the RNAi knockdown worms, did not show a change in RLS (**Figures S5B and S5C, Supplementary Table 1**). On the other hand, while *sucg-1* homozygous knockout worms appeared wild-type in the parental generation, their progeny exhibited delayed development as a result of maternal *sucg-1* deficiency. To avoid this maternal effect, we have generated a heterozygous parental line by crossing the *sucg-1* knockout line (KO) with the *sucg-1::egfp* knock-in line (GFP) (**Figure 5A**). This way, we can examine the reproductive phenotype of the progeny that carries the following genotypes: KO/KO, KO/GFP, and GFP/GFP on the *sucg-1* locus (**Figure 5A**). We found that the *sucg-1* homozygous KO/KO worms have extended RLS compared to either KO/GFP heterozygous or GFP/GFP homozygous worms (**Figures 5B and S5D, Supplementary Table 1**). These results confirm the specificity of GTP-specific SCS in regulating reproductive aging.

**Figure 5.**
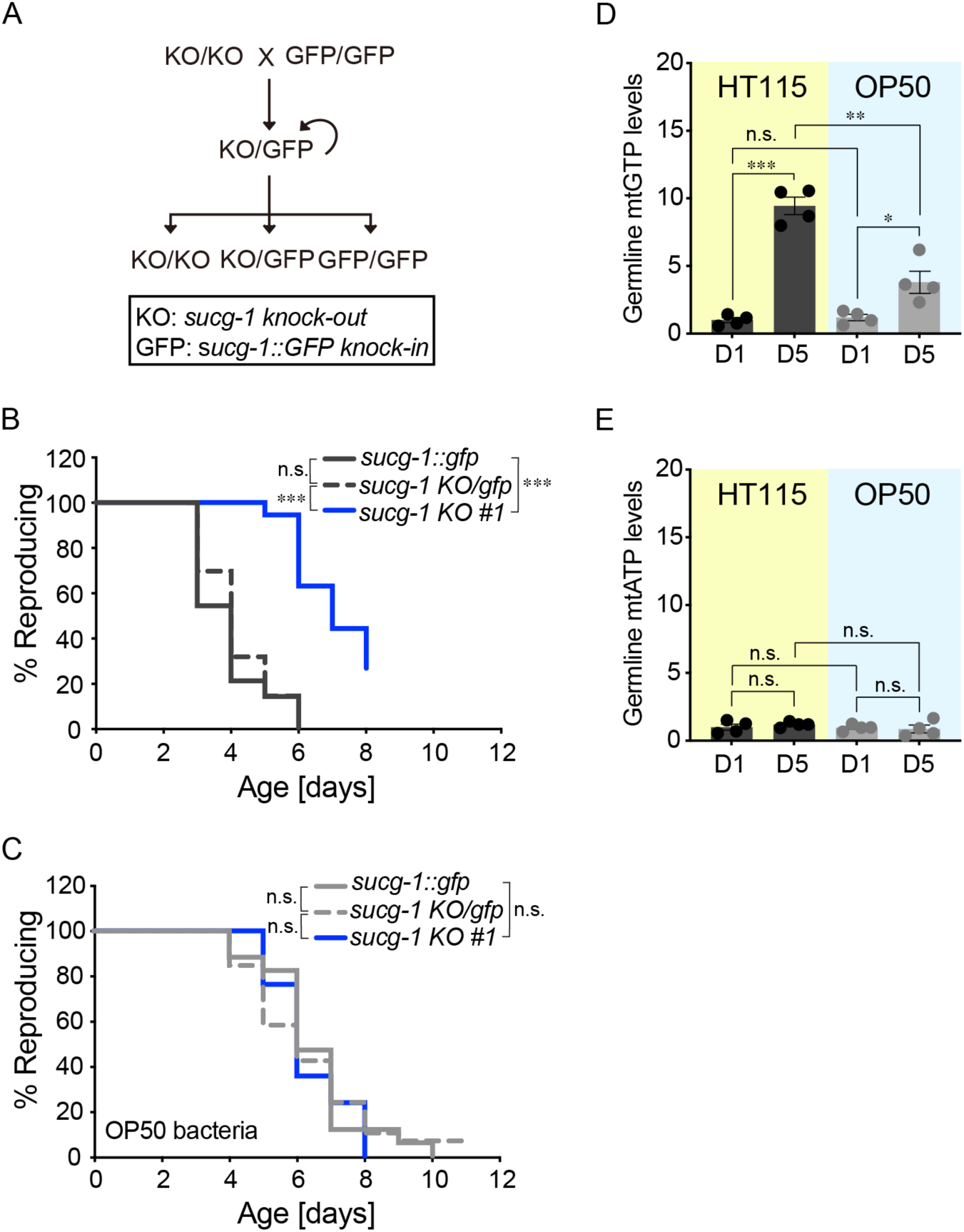
Bacterial inputs regulate germline mitochondrial GTP and reproductive aging. (A) A diagram showing the strategy to obtain *sucg-1* homozygous knockout (KO) mutants from heterozygous mutants with *sucg-1* KO at one locus and *sucg-1::egfp* (GFP) at the other locus. (B) *sucg-1* KO/KO mutants show a significant increase in RLS compared to *sucg-1* GFP/GFP and *sucg-1* KO/GFP worms. (C) With OP50 bacteria, *sucg-1* KO/KO mutants show no significant differences in RLS compared to *sucg-1* GFP/GFP or *sucg-1* KO/GFP worms. (D) Germline mitochondrial GTP (mtGTP) level is increased by 9.4-fold in day 5 aged worms compared to day 1 young worms on HT115 bacteria. With OP50 bacteria, the germline mtGTP level increase from day 1 to day 5 is 3-fold. The germline mtGTP level is higher in worms on HT115 than those on OP50 at day 5, but not at day 1. (E) Germline mitochondrial ATP (mtATP level) is not significantly different in worms of different ages and on different bacteria. (B, C) n.s. *p > 0.05,* *** *p < 0.001* by log-rank test; n = 3 biological independent replicates, ∼80 worms per replicate split into 3 genotypes, see Supplementary Table 1 (B) and Supplementary Table 3 (C) for full RLS Data. (D, E) Error bars represent mean ± s.e.m., n = 4 biologically independent samples, n.s. *p > 0.05*, * *p < 0.05,* ** *p < 0.01,* *** *p < 0.001* by Student’s t-test adjusted with the Holm–Bonferroni method for multiple comparisons.

When examining these knockout mutant worms, we also had an interesting observation that on OP50 *E. coli,* neither the *sucg-1* nor the *suca-1* homozygous knockout caused an RLS extension when compared to KO/GFP heterozygous or GFP/GFP homozygous worms (in the case of *sucg-1* KO) or wild-type worms (in the case of *suca-1* KO) respectively (**Figures 5C and S5E-G, Supplementary Table 3**). Previous findings in our lab revealed that *C. elegans* have distinct reproductive strategies when exposed to different bacteria. The wild-type worms that host OP50 *E. coli* have a longer RLS and slower oocyte quality decline with age than those on HB101 *E. coli*^19^. The wild-type worms on HT115 *E. coli* had similar RLS and late fertility to those on HB101 but were distinct from those on OP50^19^ (**Figures S6A and S6B, Supplementary Table 4**). For the germline-specific RNAi knockdown of *sucg-1, sucl-2* or *suca-1*, the experiments were conducted in the background of HT115 *E. coli* (**Figures 2D, 2E, and S6C, Supplementary Table 1**). When we examined their effects in the background of OP50 *E. coli,* we found that none of them enhances the RLS extension caused by OP50 (**Figures S6D-F, Supplementary Table 3**). These results suggest that different *E. coli* may affect mitochondrial GTP to exert different effects on worm reproductive aging.

To examine whether bacterial inputs affect mtGTP levels in the germline, we utilized the transgenic strain where germline mitochondria were tagged with GFP and triple HA and purified mitochondria using anti-HA immunoprecipitation. We then measured germline mtGTP using liquid chromatograph coupled with mass spectrometry. We found that in the germline of day 5 aged worms on HT115 *E. coli*, the mtGTP level is increased by nearly 10-fold compared to day 1 young worms, but the mtGTP induction level is only around 3-fold in the germline of worms on OP50 *E. coli* (**Figure 5D**). Moreover, day 5 aged worms on HT115 *E. coli* had a higher germline mtGTP level compared to worms on OP50 *E. coli*, while no difference in the germline mtGTP level was observed in day 1 young worms (**Figure 5D**). On the other hand, the germline mitochondrial ATP (mtATP) level were the same between worms at different ages or on different bacteria (**Figure 5E**). These results suggest that reproductive longevity conferred by OP50 *E. coli* may be linked to an attenuation in the age-related increase in GTP production.

### Bacteria modulate mitochondrial distribution during reproductive aging

Next, we examined whether OP50 *E. coli* causes changes in oocyte mitochondrial morphology using the transgenic strain expressing TOMM-20::GFP in the germline. When compared to worms on HT115 *E. coli,* worms on OP50 *E. coli* attenuated the age-associated increase in the mitochondrial clustering around the oocyte nucleus (**Figures 6A and S3E**). Moreover, germline-specific depletion of the DRP-1 protein or the germline-specific RNAi knockdown of *drp-1* fully suppressed the RLS extension in the worms on OP50 *E. coli* (**Figures 6B and S6G, Supplementary Table 3**). In addition, with the AID system, we could apply the auxin treatment only during adulthood. This way, the loss of DRP-1 occurred after the germline completes development and switches from spermatogenesis to oogenesis. We found that this adult-only depletion of DRP-1 in the germline suppresses the RLS extension in the worms on OP50 *E. coli* (**Figure 6C, Supplementary Table 3**), supporting the significance of oocyte mitochondrial distribution in regulating reproductive aging. Furthermore, the germline-specific overexpression of *drp-1* increases RLS in worms on either HT115 (**Figure 3F, Supplementary Table 1**) or OP50 *E. coli* (**Figure 6D, Supplementary Table 3**); while the germline-specific RNAi knockdown of *eat-3* could only extend RLS in worms on HT115 *E. coli* (**Figure 3B, Supplementary Table 1**) but failed to further enhance the RLS extension in worms on OP50 *E. coli* (**Figure 6E, Supplementary Table 3**). Like in worms on HT115 *E. coli*, germline-specific RNAi knockdown of *fzo-1* does not alter RLS in worms on OP50 *E. coli* (**Figure S6H, Supplementary Table 3**).

**Figure 6.**
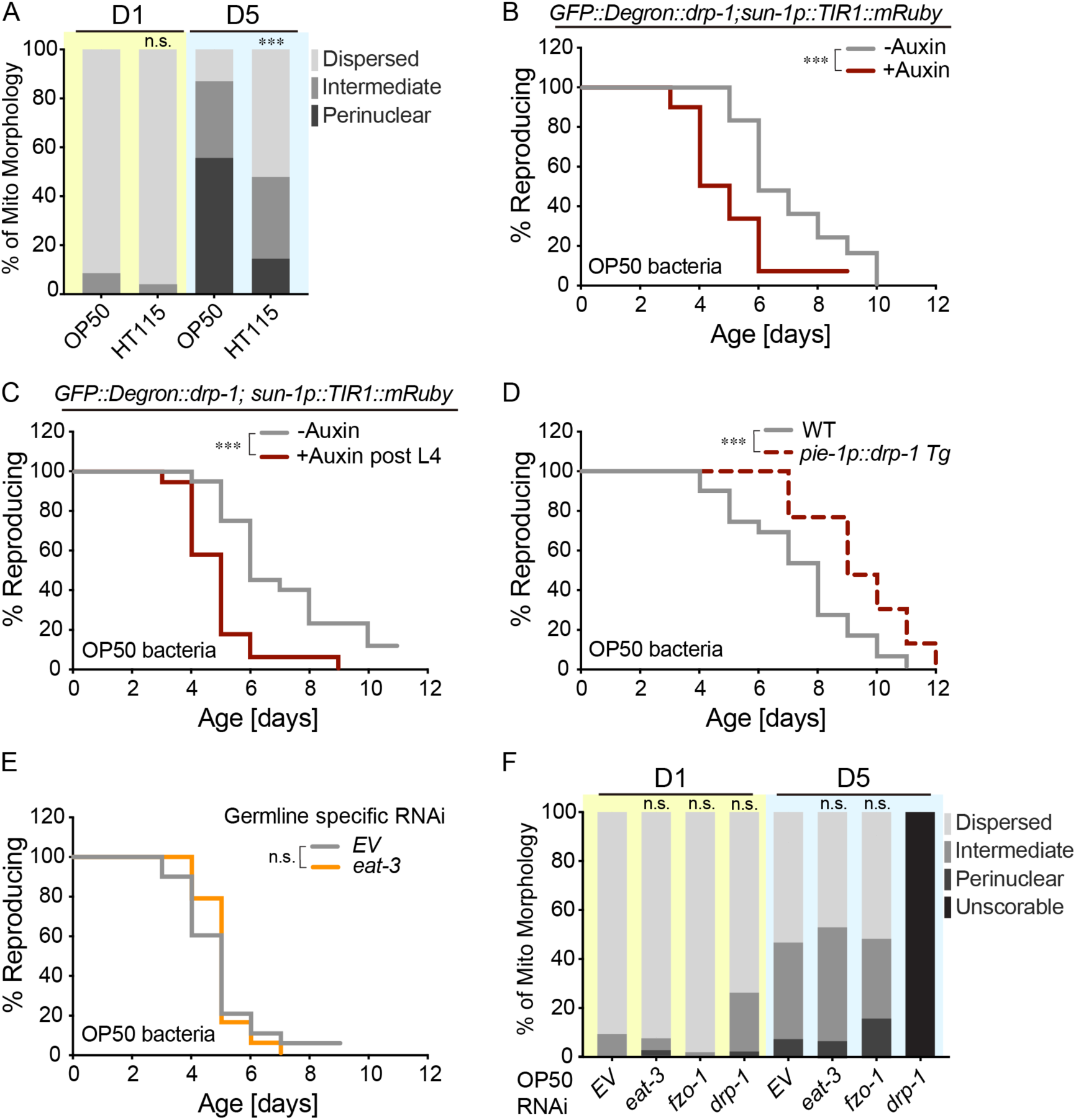
Mitochondrial dynamics factors mediate bacterial effects on reproductive longevity. (A) The perinuclear clustering of oocyte mitochondria is decreased in day 5 worms on OP50 compared to those on HT115 bacteria. (B) Auxin-induced germline-specific depletion of DRP-1 reduces RLS in worms on OP50 bacteria. (C) Adult-only depletion of DRP-1 reduces RLS in worms on OP50 bacteria. (D) Germline-specific overexpression of *drp-1* prolongs RLS in worms on OP50 bacteria. (E) Germline-specific RNAi inactivation of *eat-3* fails to extend RLS in worms on OP50 bacteria. (F) With OP50 bacteria, the distribution of oocyte mitochondria is not significantly different between control worms and those with *eat-3* or *fzo-1* RNAi knockdown at day 5. With *drp-1* RNAi knockdown, oocyte mitochondrial distribution becomes unscorable due to distorted germline. (A) n = 48 (HT115, D1), n = 53 (OP50, D1), n = 47 (HT115, D5), and n = 48 (OP50, D5); HT115 vs. OP50, n.s. *p > 0.05*, *\*\*\* p < 0.001* by Chi-squared test. (B, C, D, E) n.s. *p > 0.05*, *\*\*\* p < 0.001* by log-rank test; n = 3 (B, C, D) or 4 (E) biological independent replicates, ∼20 worms per replicate, see Supplementary Table 3 for full RLS Data. (F) n= 44 (EV, D1), n = 43 (*eat-3*, D1), n = 41 (*fzo-1*, D1), n = 42 (*drp-1*, D1), n = 43 (EV, D5), n = 43 (*eat-3*, D5), n = 43 (*fzo-1*, D5); OP50 condition; RNAi vs. EV, n.s. *p > 0.05* by Chi-squared test adjusted with the Holm– Bonferroni method for multiple comparisons.

Furthermore, we found that *drp-1* RNAi knockdown largely disturbs oocyte organization and mitochondrial distribution in day 5 aged worms on OP50 *E. coli* (**Figures 6F and S3G**), and in the small percentage of oocytes with recognizable cell boundaries, one-sided perinuclear aggregation of mitochondria was observed (**Figure S3F**). On the other hand, RNAi knockdown of either *eat-3* or *fzo-1* had no effects on oocyte mitochondrial distribution in worms on OP50 *E. Coli* (**Figures 6F and S3F**). RNAi knockdown of *sucg-1*, *sucl-2,* or *suca-1* could not alter oocyte mitochondrial distribution in worms on OP50 *E. coli* either (**Figures S6I and S3F**). Together, these results suggest that like GTP-specific SCS, OP50 bacterial inputs modulate mitochondrial distribution and reproductive longevity via mitochondrial dynamics factors.

### Vitamin B12 deficiency in OP50 *E. coli* contributes to reproductive longevity

The previous study in our lab revealed that the trace amount of HB101 *E. coli* mixing in OP50 *E. coli* is sufficient to shorten RLS of *C. elegans*, suggesting the involvement of bioactive metabolites in regulating reproductive aging. Interestingly, it is known that OP50 *E. coli* is low in vitamin B12 (VB12), and the VB12 level affects mitochondrial dynamics in the muscle of *C. elegans*^45–47^. To test whether VB12 deficiency in OP50 *E. coli* could contribute to reproductive longevity, we supplied two different forms of VB12, methylcobalamin (meCbl) and adenosylcobalamin (adoCbl) to worms on OP50 and HT115 *E. coli*. We discovered that supplementation of either meCbl or adoCbl reduces the RLS extension in worms on OP50 *E. coli* but does not affect RLS in worms on HT115 *E. coli* (**Figures 7A, 7B, S7A, and S7B, Supplementary Table 1 and 3**). In addition, meCbl supplementation increased the perinuclear accumulation of oocyte mitochondria in day 5 worms on OP50 *E. coli* (**Figures 7C and S3H**), to the level similar in worms on VB12 sufficient HT115 *E. coli*. These results suggest that bacteria-derived VB12 plays a crucial role in regulating oocyte mitochondrial distribution and reproductive aging.

**Figure 7.**
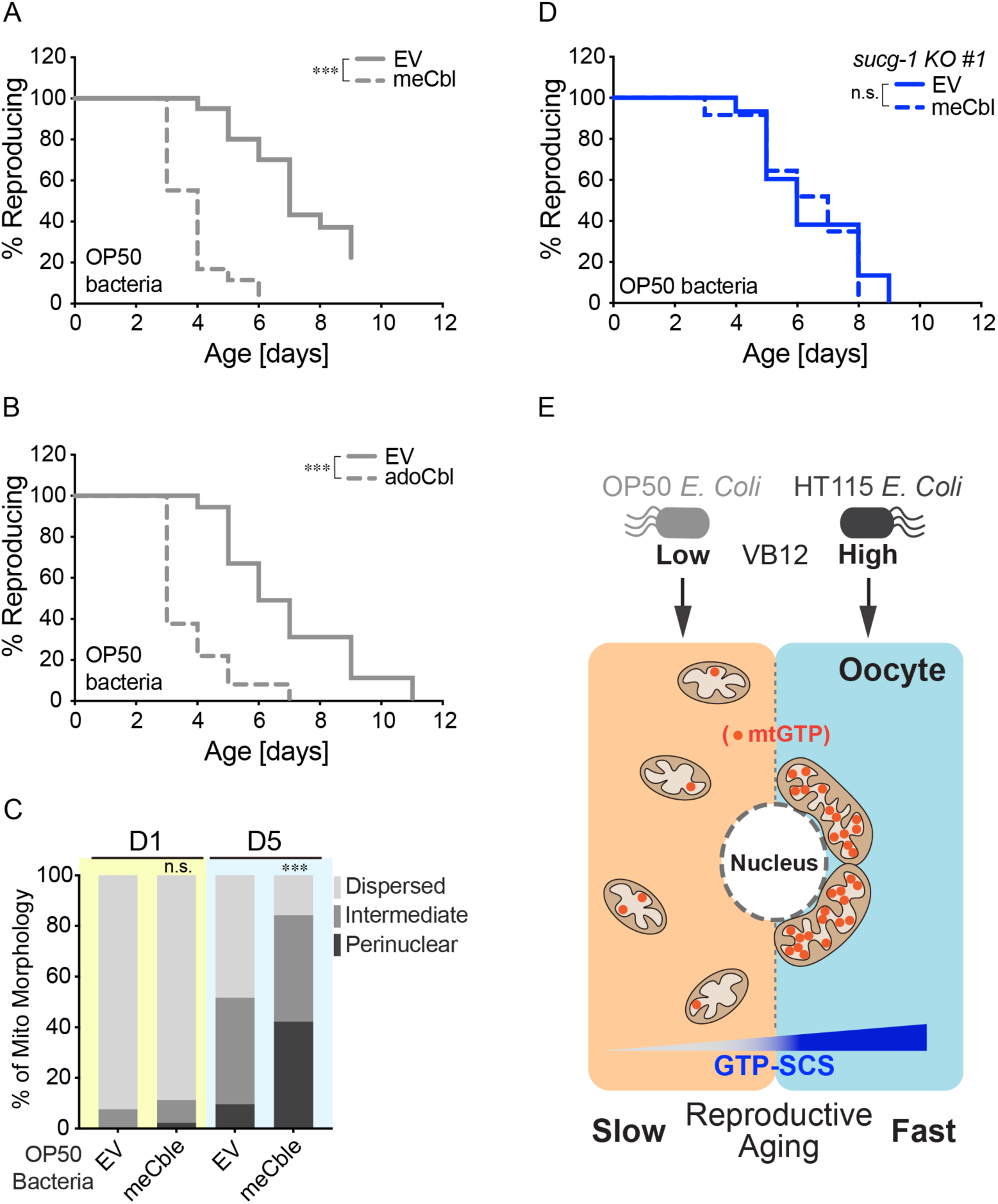
Bacterial VB12 regulates oocyte mitochondria and reproductive aging. (A, B) Supplementation of meCbl or adoCbl shortens RLS of WT worms on OP50 bacteria. (C) Supplementation of meCbl increases the perinuclear clustering of oocyte mitochondria in WT worms on OP50 bacteria at day 5. (D) Supplementation of meCbl does not shorten RLS of the *sucg-1* mutant worms on OP50 bacteria. (E) Summary model representing mitochondrial GTP metabolism and mitochondrial dynamics couple in the oocyte to regulate reproductive longevity, which is modulated by metabolic inputs from bacteria. (A, B) *\*\*\* p < 0.001* by log-rank test; n = 3 biological independent replicates, ∼20 worms per replicate, see Supplementary Table 3 for full RLS Data. (C) n= 40 (EV, D1), n = 45 (128nM meCbl, D1), n = 42 (EV, D5), n = 38 (128nM meCbl, D5); OP50 condition; 128nM meCbl vs EV, n.s. *p > 0.05, *** p < 0.001* by Chi-squared test. (D) n.s. *p > 0.05* by log-rank test; n = 3 biological independent replicates, ∼80 worms per replicate split into 3 genotypes, see Supplementary Table 3 for full RLS Data.

We further examined whether VB12 signals through GTP-specific SCS to modulate reproductive aging. We found that although the *sucg-1* heterozygous mutant (KO/GFP) and *gfp* homozygous (GFP/GFP) worms experience a decrease in RLS when supplied with meCbl, this decrease was not observed in the *sucg-1* homozygous (KO/KO) mutant worms (**Figures 7D, S7C, and S7D, Supplementary Table 3**). This result suggests that SUCG-1 is required for VB12 to regulate reproductive aging. Two enzymes utilize VB12 as cofactor for their function, namely MMCM-1, which is a mitochondrial enzyme that converts methymalonyl-CoA to succinyl-CoA, and METR-1, the methionine synthase that converts homocysteine to methionine. We discovered that *mmcm-1* RNAi knockdown does not affect RLS in worms on either OP50 or HT115 *E. coli* (**Figures S7E and S7F, Supplementary Table 1 and 3**). These results further support that succinyl-CoA is not involved in the regulation of reproductive aging. On the other hand, *metr-1* RNAi knockdown increased RLS in worms on HT115 but not OP50 *E. coli* (**Figures S7E and S7F, Supplementary Table 1 and 3**), suggesting that the VB12-methionine synthase branch, which controls purine synthesis^48, 49^, mediates the bacterial effect on reproductive aging.

## DISCUSSION

In summary, our work discovered that mitochondrial GTP-specific SCS plays a crucial role in regulating oocyte mitochondrial distribution and reproductive health during aging, and further revealed that bacterial inputs act through this mechanism to modulate reproductive longevity (**Figure 7E**). We found that mitochondria exhibit dispersed structure in young oocytes, but undergo perinuclear clustering in aged oocytes. Interestingly, the similar age-associated change in oocyte mitochondrial distribution was also observed in mice, which has been linked to decreased Drp1 activity^10^. In our studies, we found that germline-specific overexpression of *drp-1* is sufficient to prolong RLS, through suppressing perinuclear clustering of oocyte mitochondria. These findings demonstrate an evolutionally conversed role of DRP1-directed mitochondrial fission in regulating reproductive health.

The knockdown of *eat-3* or *fzo-1* should tilt the balance toward mitochondrial fission as well. However, their effects on reproductive longevity are distinct. While the *eat-3* RNAi knockdown was sufficient to prolong reproductive lifespan and improve late fertility, the *fzo-1* RNAi knockdown failed to do so. In mice, oocyte-specific knockout of Mfn1 but not Mfn2 results in increased mitochondrial clustering, as well as defective folliculogenesis, impaired oocyte quality and sterility^12^. Thus, different mitochondrial fusion factors may play distinctive roles in regulating oocyte quality and reproductive aging. Moreover, unlike the germline-specific overexpression of *drp-1* that extended RLS in both HT115 and OP50 bacterial background, the germline-specific knockdown of *eat-3* could not further enhance the RLS extension caused by OP50 bacteria. In addition to its requirement for the RLS extension conferred by OP50 bacteria, DRP-1 is reported in a recent study to be necessary for the RLS extension in the mutant of *daf-2*, the *C. elegans* homolog of insulin and IGF-1 receptor^50^. Our studies also showed that the RLS extension caused by the loss of the GTP-specific SCS is dependent on DRP-1. These findings together suggest that multiple regulatory mechanisms may converge on the mitochondrial fission factor DRP-1 to regulate the reproductive aging process. Therefore, increasing DRP-1 levels selectively in the reproductive system is an effective way to promote reproductive longevity, which drives mitochondrial dynamics toward fission without disrupting the fusion process.

It’s important to note that although most studies related to mitochondrial dynamics factors focus on mitochondrial morphology (tubular vs fragmented), their regulation of mitochondrial distribution has been observed in both oocytes and somatic cells. In aged mice and mice with Drp1 KO, the oocyte mitochondrial network is aggregated toward the perinuclear region and only a small part of the mitochondrial network exhibits tubular morphology^10^. Interestingly, calcium homeostasis that is crucial for oocyte quality is also disrupted in these mice, which attributes to increased ER-mitochondria aggregation^10^. Moreover, in mouse oocytes overexpressing Mfn1 or Mfn2, the mitochondrial network becomes aggregated toward the perinuclear region without increasing tubular elongation^11^. The Mfn1-induced perinuclear aggregation of mitochondria results in disrupted chromosome alignment and disorganized spindle formation in oocytes^11^. In somatic cells, perinuclear clustering of mitochondria is also observed when mitochondrial dynamics factors are modified. Notably, Drp1 knockdown abrogates mitochondria mobilization toward peripheral immune synapse following T-cell activation^51^. In pancreatic beta cells, Drp1 knockout results in mitochondria clustering on one side of the nucleus^52^. Similarly, OPA1 overexpression leads to perinuclear clustering of mitochondria in HeLa cells^53^, and overexpression of MFN2 causes perinuclear clustering of mitochondria in multiple cell types^54–57^.

Now, our data support that the key role of mitochondrial dynamic factors in regulating reproductive aging is predominantly attributed to their control of mitochondrial positioning in oocytes. Perinuclear clustered mitochondria have been associated with cellular stress, such as viral infection, heat shock, hypoxia, and apoptotic stress^58–64^. Transient perinuclear clustering may help elicit transcriptional responses^58^ and sequester damaged mitochondria^65^, to restore mitochondrial homeostasis. However, prolonged perinuclear clustering of oocyte mitochondria in aged worms and mice could block mitophagy-mediated clearance of damaged mitochondria, increase ER-mitochondria aggregation to impair calcium homeostasis, as well as disrupt mitochondrial segregation required for cell division upon fertilization. Our studies provide direct evidence that dietary and genetic interventions that drive mitochondrial dispersion from the perinuclear cluster sufficiently promote reproductive longevity in worms. It would be interesting to test whether similar mechanisms could help improve reproductive health during aging in mammals.

It is known that mitochondrial dynamics is influenced by cellular metabolism^66^. However, whether and how mitochondrial metabolism is directly linked with mitochondrial dynamics remains poorly understood. Our studies reveal that the GTP-specific SCS regulates mitochondrial dynamics in oocytes during the aging process and contributes to the regulation of reproductive longevity. SCS locates in the mitochondrial matrix, likely to be very close to the mitochondrial inner membrane. It is reported that various enzymes in the TCA cycle interact closely and form a metabolon to facilitate their reactions^67^. One of these enzymes, succinate dehydrogenase, is part of the respiratory complex II and anchored in the mitochondrial inner membrane^68^. Given that SCS provides succinate for succinate dehydrogenase as a substrate, the interaction between these two enzymes may recruit SCS close to the inner membrane, leading to a high local GTP level when the TCA cycle is active. Interestingly, it was reported that inner mitochondrial membrane fusion requires a higher concentration of GTP than outer mitochondrial fusion^69^. Furthermore, members of another family of GTP-producing enzymes, nucleoside diphosphate kinases, have been shown to directly interact with OPA1 in the mitochondrial inner membrane to regulate mitochondrial membrane dynamics in human cells^36, 70^. Our studies found that the germline-loss of EAT-3/OPA1, but not FZO-1/MFN1/2 recapitulates the effect of the germline-loss of SUCG-1 in promoting reproductive longevity. Considering the age-associated increase in the germline SUCG-1 level, it is possible that an increase in GTP production close to the inner membrane drives mitochondrial fusion via EAT-3 during the reproductive aging process, and this imbalance of mitochondrial dynamics consequently contributes to the decline of oocyte quality.

Upon the exposure to OP50 *E. coli*, worms exhibited extended RLS, which was suppressed by VB12 suppression but could not be further enhanced by the *sucg-1* knockdown. However, the *sucg-1* knockdown could sufficiently restore RLS extension in worms on OP50 *E. coli* supplemented with VB12. Based on these results, we speculate that bacterial VB12 accelerates reproductive aging through increasing germline mtGTP levels. Interestingly, RNAi knockdown of *metr-1*, which encodes the VB12-dependent methionine synthase (MTR), extends RLS in the background of HT115 but not OP50 *E. coli*. MTR catalyzes the production of methionine from homocysteine, in accordance with converting 5-methyl-tetrahydrofolate into tetrahydrofolate (THF). Two recent studies discovered that THF replenishing by MTR promotes tumor growth by supporting purine synthesis, and MTR loss results in decreased GTP and ATP levels^48, 49^. Thus, on HT115 *E. coli,* the higher level of VB12 could increase the MTR-mediated metabolic process, leading to more GTP synthesis in the cytosol and in turn a higher mtGTP level. Consistently, we found that the germline mtGTP level is elevated in day 5 aged worms, which is likely a result of the age-associated increase in GTP-specific SCS. Moreover, this increase in the germline mtGTP level is significantly greater in the background of HT115 *E. coli* than OP50 *E. coli*, which is likely due to the high VB12-MTR level associated with HT115. At present, we do not have direct evidence on how bacteria-derived VB12 modulates GTP-specific SCS in the reproductive system, aside from genetic analysis confirming the requirement of SUCG-1 for the effect of VB12. No mRNA or protein level difference was detected between OP50 and HT115 conditions. It is possible that VB12 influences the activity and/or substrate availability of GTP-specific SCS in oocyte mitochondria, which remains to be determined in future studies.

We discovered that low environmental VB12 levels are associated with reproductive longevity. There are significant variations in VB12 levels among different bacterial species. It is known that high levels of VB12 in the *Comamonas* DA1877 diet results in decreased fertility in *C. elegans*^45, 71^. On the other hand, *Qin et al.* reported that early-life VB12 deficiency is associated with adulthood sterility caused by germline ferroptosis in *C. elegans*^72^. Furthermore, in humans, a high maternal VB12 level at birth is associated with an increased risk of developing autism spectrum disorder in children^73^. However, VB12 deficiency can also lead to adverse maternal and child health problems^74, 75^, and an adequate amount of VB12 supplementation during pregnancy is recommended by the World Health Organization. Thus, there may be an antagonistic pleiotropy-like effect at the nutrient level, wherein VB12 is essential for appropriate development of germline and progeny, but later accelerates reproductive decline during aging. Our study suggests that environmental inputs from the microbiota should also be taken into account when considering this antagonistic pleiotropic effect.

**Figure S1.**
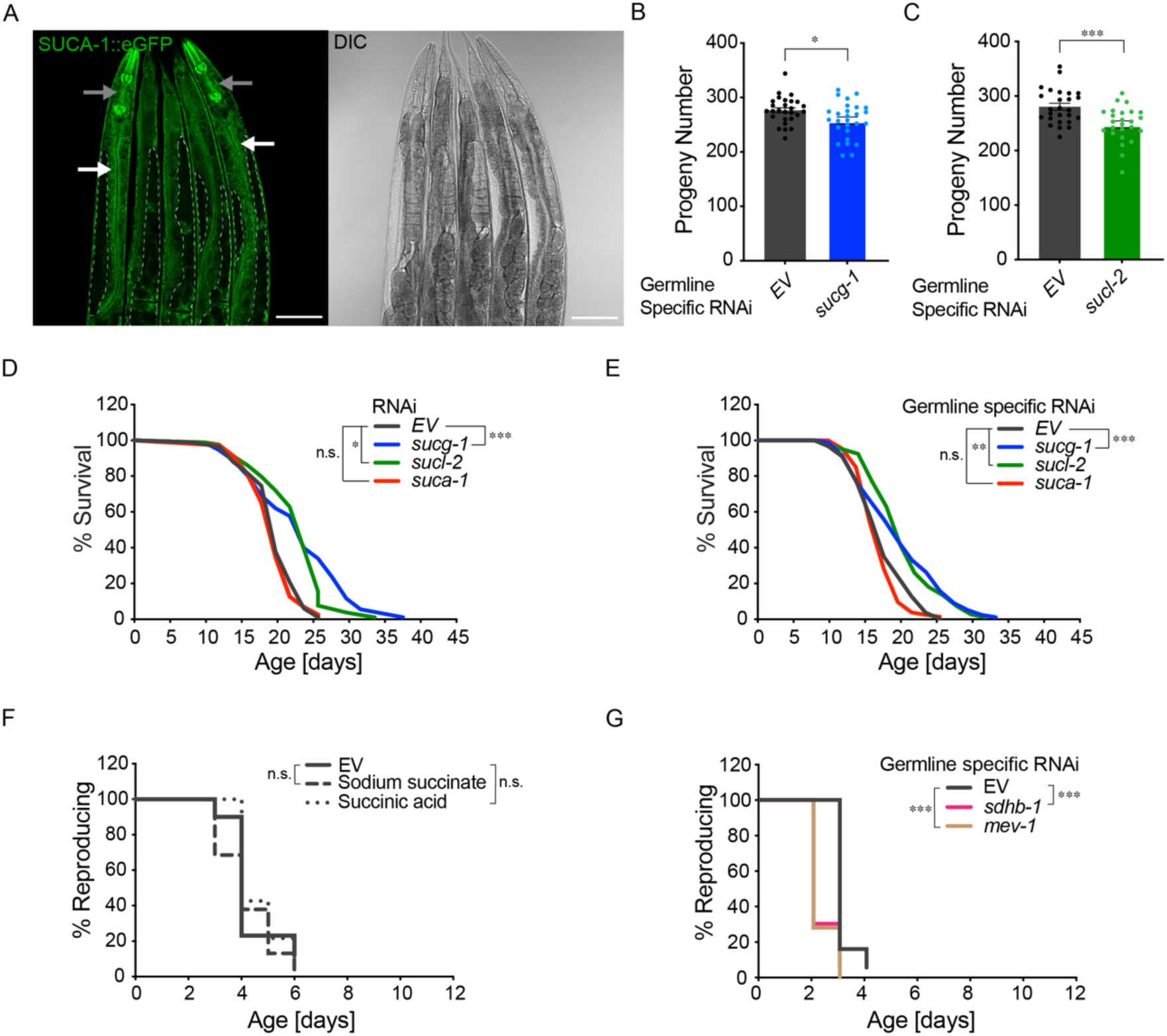
The effect of SCS on reproduction and lifespan, the effect of succinate or succinyl-CoA on RLS. (A) Confocal imaging of the SUCA-1::eGFP knock-in line, in which the endogenous *suca-1* is tagged with e*gfp,* reveals its predominant expression in the intestine, pharynx, muscle, hypodermis, and neurons, while only weak expression in the germline (Scale bar: 100μm; White arrow: intestine; Gray arrow: neuron; Dashed white line: germline). (B, C) Germline-specific RNAi knockdown of either *sucg-1* or *sucl-2* results in a slight decrease in progeny number. (D) RNAi knockdown of *sucg-1* or *sucl-2,* but not *suca-1* slightly extends lifespan. (E) Germline-specific RNAi knockdown of *sucg-1* or *sucl-2,* but not *suca-1* leads to a slight lifespan extension. (F) Supplementation of either succinic acid or sodium succinate does not affect RLS of WT worms. (G) Germline-specific RNAi knockdown of either *mev-1* or *sdhb-1* shortens RLS. (B) n= 27 (EV), n = 27 (*sucg-1*); RNAi vs EV, *\* p < 0.05* by Student’s t-test. (C) n= 26 (EV), n = 25 (*sucl-2*); RNAi vs EV, *\*\*\* p < 0.001* by Student’s t-test. (D, E) n.s. *p > 0.05*, *\* p < 0.05, ** p < 0.01, *** p < 0.001* by log-rank test; n = 3 biological independent replicates, 70∼120 worms per replicate, see Supplementary Table 2 for full lifespan data. (F, G) n.s. *p > 0.05*, *\*\*\* p < 0.001* by log-rank test; n = 3 biological independent replicates, ∼20 worms per replicate, see Supplementary Table 1 for full RLS Data.

**Figure S2.**
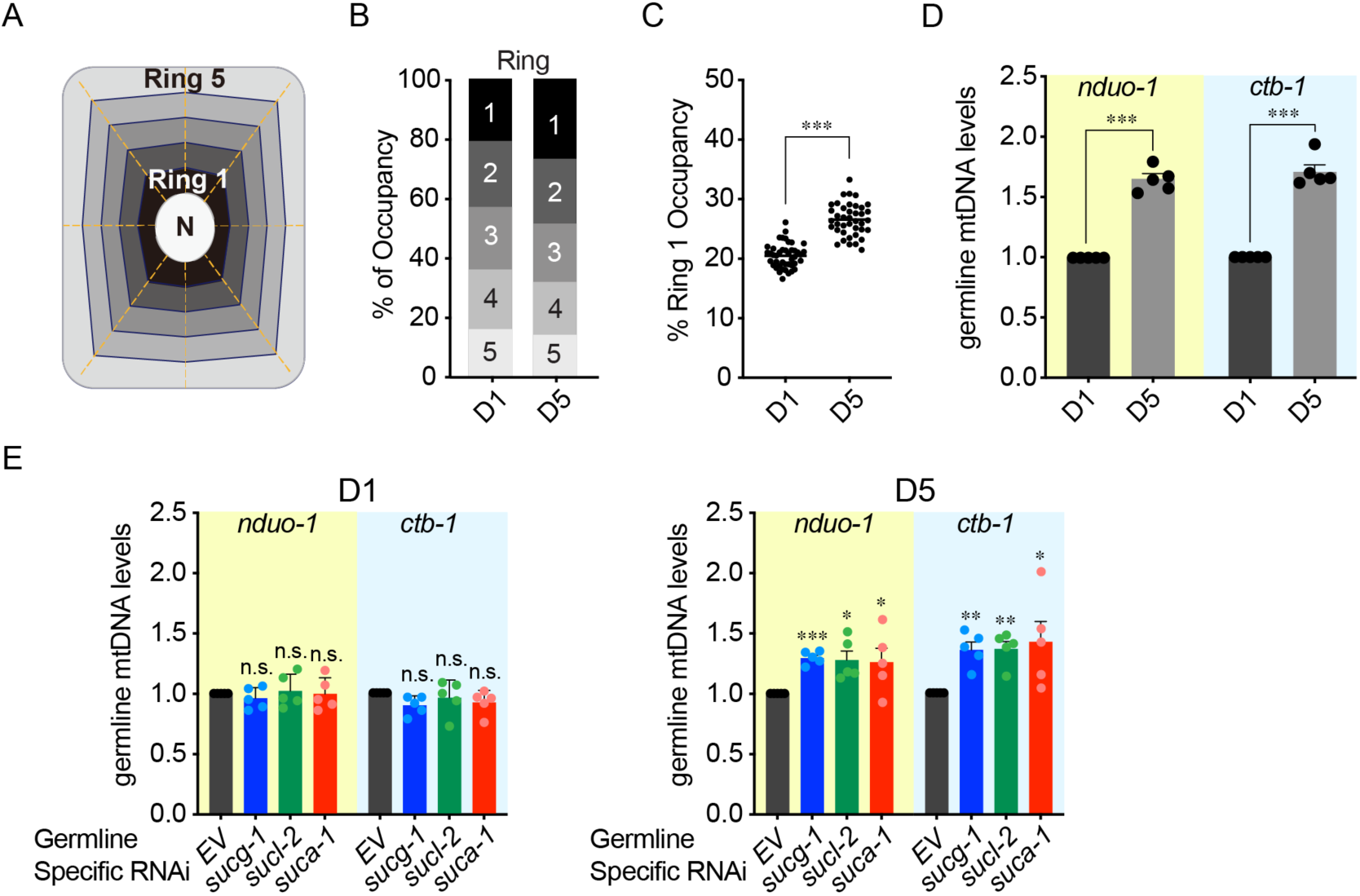
Quantification of mitochondrial positioning and mtDNA levels. (A) A computer algorithm is used to automatically divide oocytes into five rings, with ring 1 adjacent to the nucleus and ring 5 adjacent to the plasma membrane. (B) Mitochondrial GFP signal intensity in each ring is measured to calculate the percentage of mitochondrial occupancy at both day 1 and day 5. (C) The percentage of mitochondrial occupancy in ring 1 increased from day 1 to day 5. (D) Germline mtDNA level is increased in WT worms from day 1 to day 5. (E) Germline-specific RNAi knockdown of *sucg-1*, *sucl-2*, or *suca-1* does not affect the germline mtDNA level at day 1 but causes an increase at day 5. (C) n = 43 (day 1) and n = 40 (day 5); *** *p < 0.001* by Student’s t-test. (D, E) n.s. *p > 0.05*, *\* p < 0.05, ** p < 0.01, *** p < 0.001* by Student’s t-test adjusted with the Holm–Bonferroni method for multiple comparisons; n = 5 biological independent replicates.

**Figure S3.**
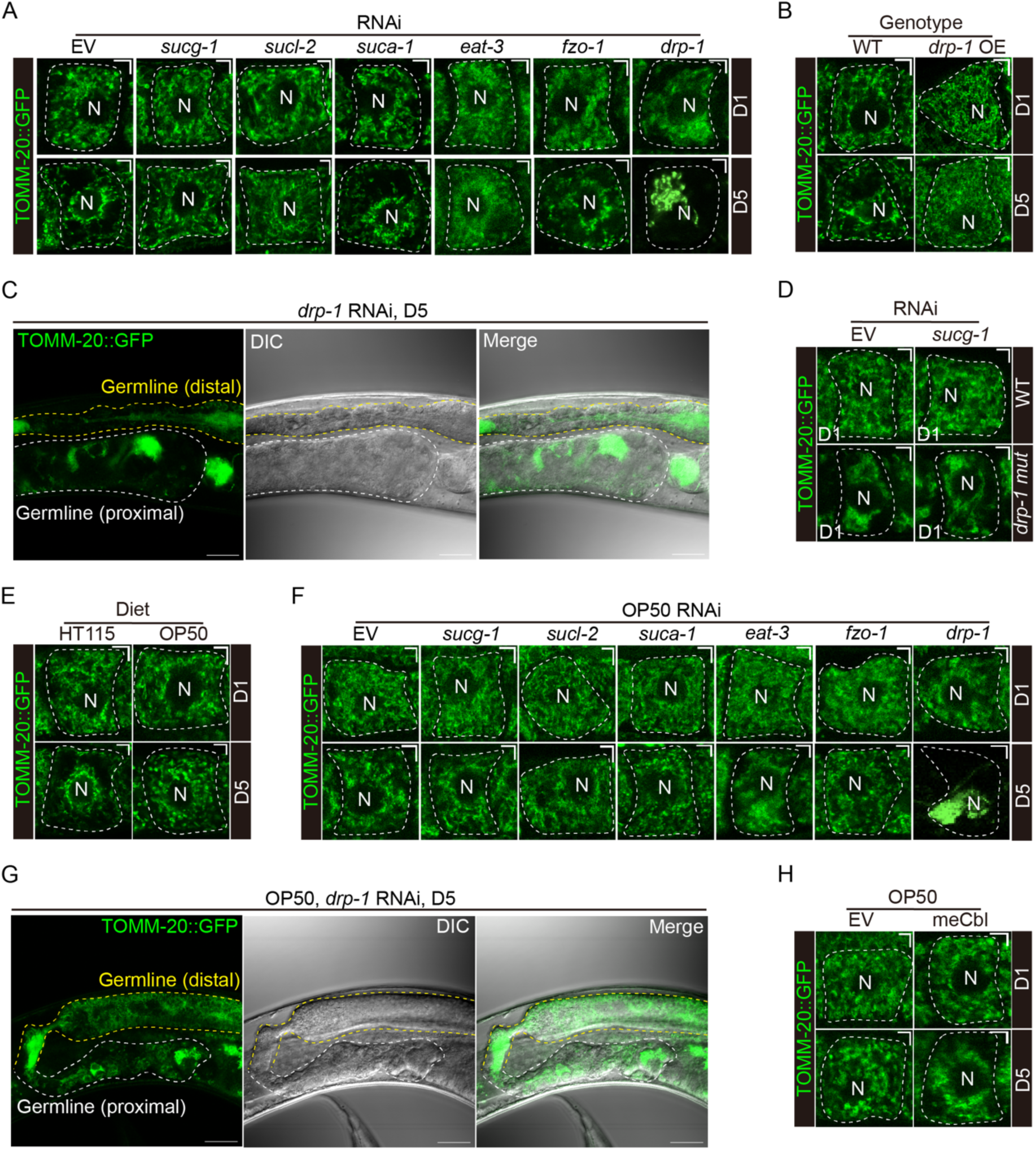
Representative images showing oocyte mitochondrial distribution under different conditions. (A) Representative images reveal that RNAi inactivation of *sucg-1, sucl-2,* or *eat-3* but not *suca-1 or fzo-1* suppresses the perinuclear clustering of oocyte mitochondria in worms at day 5. Upon *drp-1* RNAi knockdown, few oocytes with a recognizable cell boundary at day 5 show the one-sided perinuclear clustering of mitochondria (Scale bar: 5μm; Dashed white line: oocyte outline; N: nucleus). (B) Representative images of oocyte mitochondria show that the perinuclear clustering distribution at day 5 is suppressed by the germline-specific overexpression of *drp-1* (Scale bar: 5μm; Dashed white line: oocyte outline; N: nucleus). (C) Germline morphology and mitochondrial distribution of day 5 worms subjected to *drp-1* RNAi knockdown become largely disturbed and unorganized. (Scale bar: 30μm; Dashed white line: proximal germline; Dashed yellow line: distal germline). (D) Representative images reveal that RNAi inactivation of *sucg-1* does not suppress the perinuclear distribution of oocyte mitochondria in the *drp-1* mutant worms at day 1 (F, Scale bar: 5μm; Dashed white line: oocyte outline; N: nucleus). (E) Representative images of oocyte mitochondria show that the perinuclear clustering distribution at day 5 is suppressed in worms on OP50 bacteria compared to those on HT115 bacteria (Scale bar: 5μm; Dashed white line: oocyte outline; N: nucleus). (F) Representative images reveal that RNAi inactivation of *sucg-1, sucl-2, suca-1, eat-3* or *fzo-1* has no effect on the distribution of oocyte mitochondria in worms on OP50 bacteria at both day 1 and day 5. Upon *drp-1* RNAi knockdown, few oocytes with a recognizable cell boundary at day 5 show the one-sided perinuclear clustering of mitochondria (F, Scale bar: 5μm; Dashed white line: oocyte outline; N: nucleus). (G) With OP50 bacteria, germline morphology and mitochondrial distribution of day 5 worms subjected to *drp-1* RNAi knockdown become largely disturbed and unorganized. (Scale bar: 30μm; Dashed white line: proximal germline; Dashed yellow line: distal germline). (H) Representative images reveal that meCbl supplementation induces the perinuclear clustering of oocyte mitochondria in worms on OP50 bacteria at day 5 (F, Scale bar: 5μm; Dashed white line: oocyte outline; N: nucleus).

**Figure S4.**
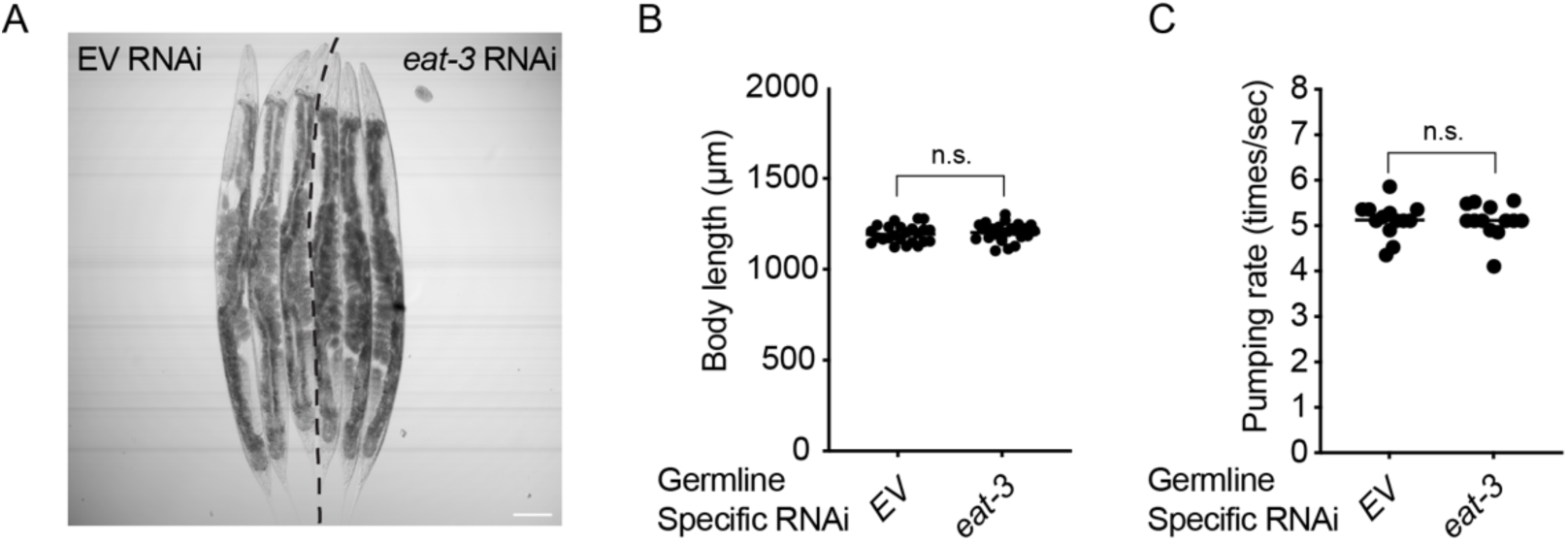
Effects of germline-specific *eat-3* knockdown on body size and food intake. (A, B) Germline-specific RNAi knockdown of *eat-3* does not affect the body size of day 1 worms, representative images are shown in A (Scale bar: 100μm), and the quantification results are shown in B. (C) The pharyngeal pumping rate of day 1 worms subjected to germline-specific RNAi knockdown of *eat-3* is not significantly different from the control. (A, B) n = 24 (EV), n = 24 (*eat-3*); RNAi vs EV, n.s. *p > 0.05* by Student’s t-test. (C) n = 13 (EV), n = 13 (*eat-3*); RNAi vs EV, n.s. *p > 0.05* by Student’s t-test.

**Figure S5.**
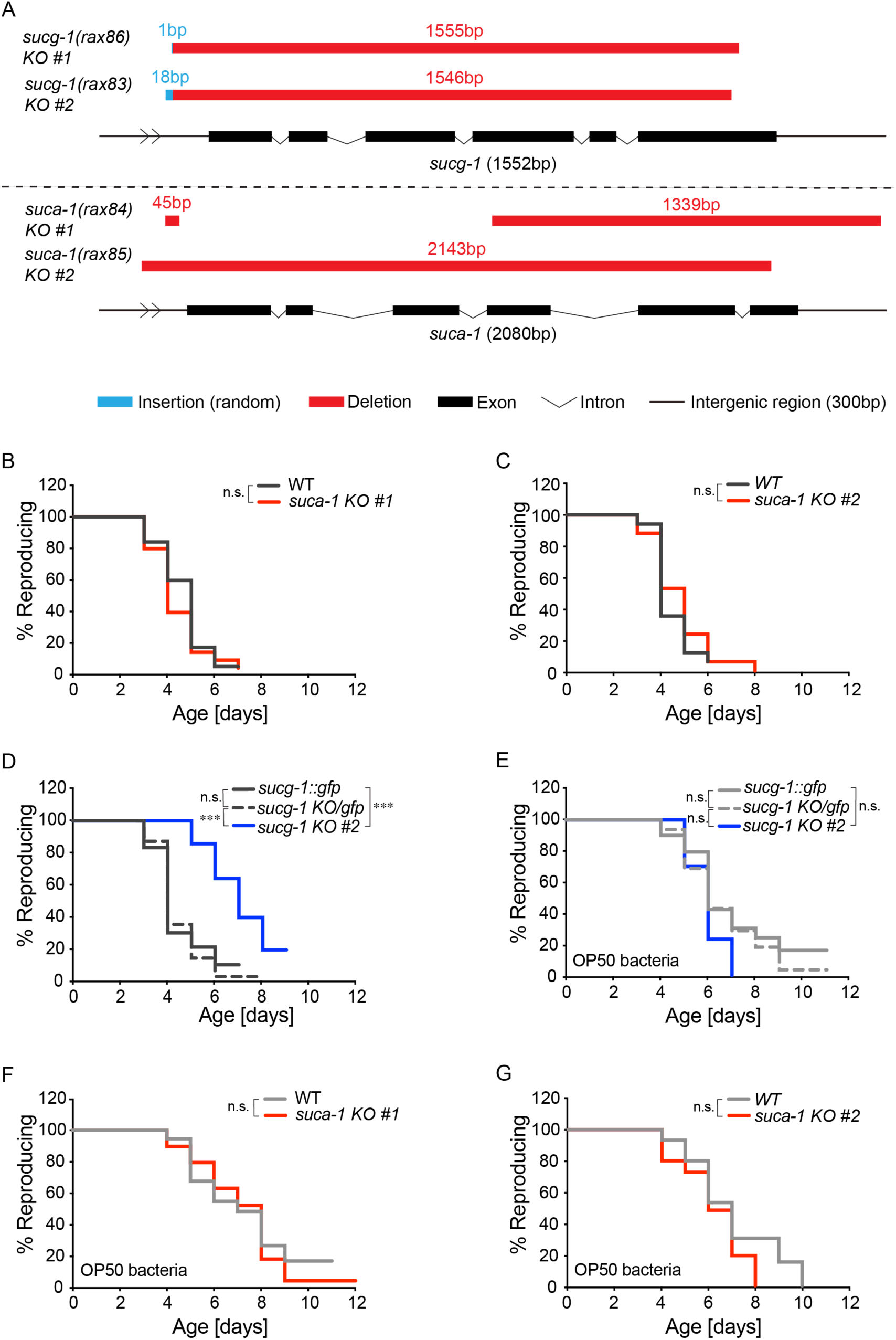
Effects of SCS knockout on RLS under different bacterial conditions. (A) A diagram showing the knockout (KO) lines of *sucg-1* and *suca-1* used in this study, *sucg-1(rax86)* is used in experiments shown in Main Figures. (B, C) *suca-1(rax84)* and *suca-1(rax85)* KO worms show no significant differences in RLS compared to WT worms. (D) *sucg-1(rax83)* KO mutants show a significant increase in RLS compared to *sucg-1::egfp* and *sucg-1* KO/*egfp* worms. (E) With OP50 bacteria, *sucg-1(rax83)* KO mutants show no significant differences in RLS compared to *sucg-1::egfp* or *sucg-1* KO/*egfp* worms. (F) With OP50 bacteria, *suca-1(rax84)* KO worms show no significant differences in RLS compared to WT animals. (G) With OP50 bacteria, *suca-1(rax85)* KO worms show no significant differences in RLS compared to WT animals. (B, C, F, G) n.s. *p > 0.05* by log-rank test; n = 3 biological independent replicates, ∼20 worms per replicate, see Supplementary Table 1 (B, C) and Supplementary Table 3 (F, G) for full RLS Data. (D, E) n.s. *p > 0.05,* *** *p < 0.001* by log-rank test; n = 3 biological independent replicates, ∼80 worms per replicate split into 3 genotypes, see Supplementary Table 1 (D) and Supplementary Table 3 (E) for full RLS Data.

**Figure S6.**
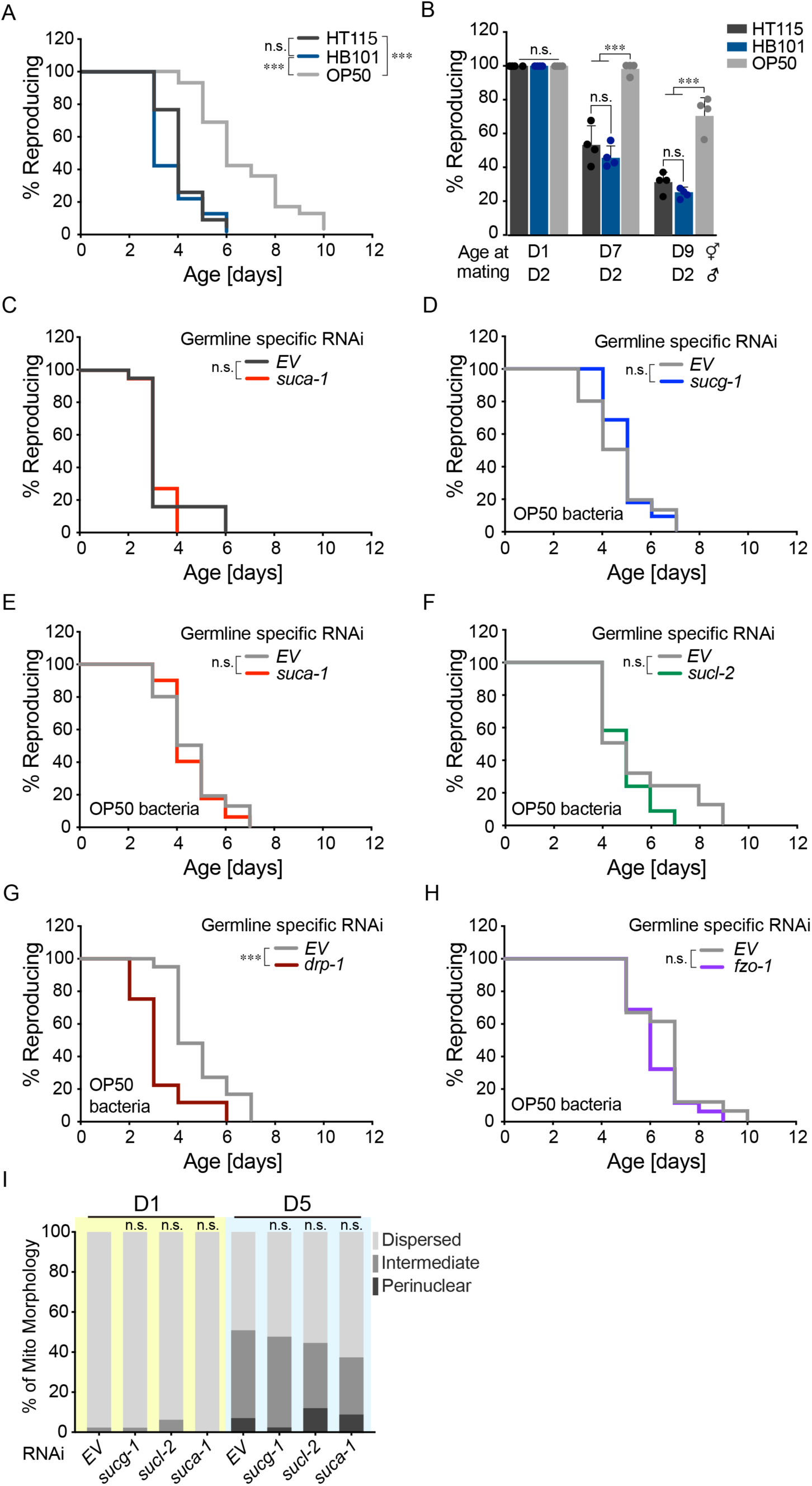
RLS regulation under different bacterial conditions. (A) WT worms on HT115 and HB101 show no significant differences in RLS, while WT worms with OP50 show significantly longer RLS compared to both HT115 and HB101. (B) Day 7 and 9 WT hermaphrodites with OP50 show higher rates of reproduction than those with HT115 or HB101, when mated with day-2-old young males. (C) Germline-specific RNAi inactivation of *suca-1* does not affect RLS in the background of HT115 bacteria. (D-F) With OP50 bacteria, germline-specific RNAi inactivation of *sucg-1, sucl-2,* or *suca-1* does not extend RLS. (G) Germline-specific RNAi inactivation of *drp-1* shortens RLS on OP50 bacteria. (H) Germline-specific RNAi inactivation of *fzo-1* does not show significant differences in RLS on OP50 bacteria. (I) With OP50, RNAi inactivation of *sucg-1, sucl-2, or suca-1* has no effect on oocyte mitochondrial distribution in worms at day 1 and day 5. (A, C, D, E, F, G, H) n.s. *p > 0.05,* *** *p < 0.001*, by log-rank test; n = 3 (A, C, E, F, G, H) or 4 (D) biological independent replicates, ∼20 worms per replicate, see Supplementary Table 1 (C), Supplementary Table 3 (D, E, F, G, H), and Supplementary Table 4 (A) for full RLS Data. (B) Error bars represent mean ± s.e.m., n = 4 biologically independent samples, n.s. *p > 0.05*, *** *p < 0.001* by Fisher’s exact test adjusted with the Holm–Bonferroni method for multiple comparisons, ∼15 worms per replicate. (I) n= 43 (EV, D1), n = 46 (*sucg-1*, D1), n = 45 (*sucl-2*, D1), n = 45 (*suca-1*, D1), n = 43 (EV, D5), n = 42 (*sucg-1*, D5), n = 43 (*sucl-2*, D5) and n = 45 (*suca-1*, D5); OP50 condition; RNAi vs. EV, n.s. *p > 0.05* by Chi-squared test adjusted with the Holm–Bonferroni method for multiple comparisons.

**Figure S7.**
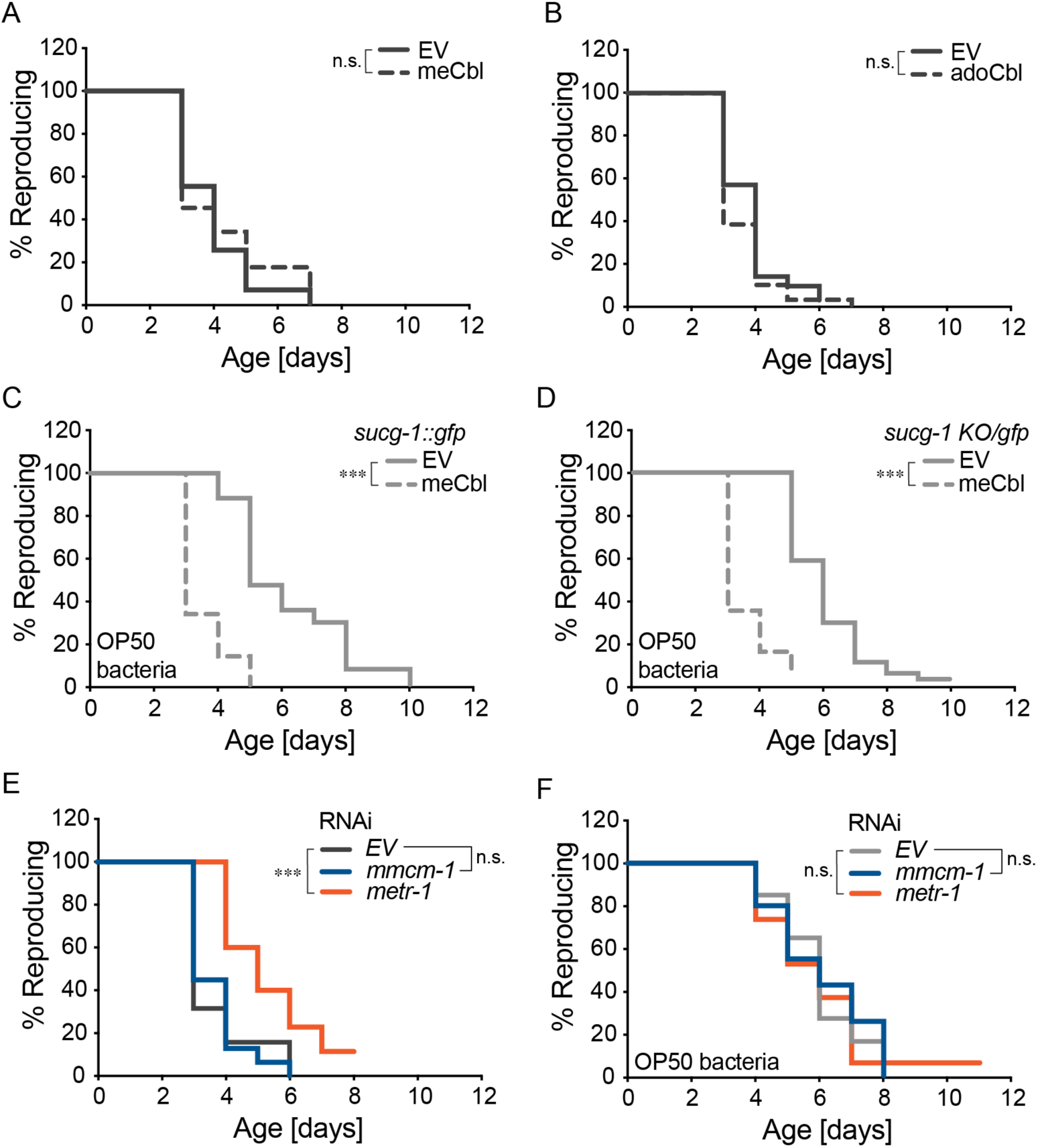
VB12 and its downstream effectors mediate bacterial regulation of RLS. (A, B) Supplementation of meCbl or adoCbl does not alter RLS of WT worms on HT115 bacteria. (C, D) Supplementation of meCbl significantly shortened RLS of *sucg-1::gfp* and *sucg-1 gfp/KO* control worms on OP50 bacteria. (E) On HT115 bacteria, WT worms subjected to *metr-1*, but not *mmcm-1* RNAi knockdown show a significant increase in RLS compared to those subjected to the EV control. (F) With OP50 bacteria, WT worms subjected to *mmcm-1* or *metr-1* RNAi show no significant differences in RLS compared to those subjected to the EV control. (A, B, E, F) n.s. *p > 0.05,* *** *p < 0.001* by log-rank test; n = 3 biological independent replicates, ∼20 worms per replicate, see Supplementary Table 1 (A, B, E) and Supplementary Table 3 (F) for full RLS Data. (C, D) *** *p < 0.001* by log-rank test; n = 3 biological independent replicates, ∼80 worms per replicate split into 3 genotypes, see Supplementary Table 3 for full RLS Data.

## Materials and Methods

### Strains and maintenance

*C. elegans* strains N2, DCL569, EGD629, EGD623, EU2917, CA1472, and CU6372 were obtained from the Caenorhabditis Genetics Center. PHX3617 and PHX4685 were acquired from Suny Biotech. MCW618, MCW1220, MCW1315, MCW1325, MCW1326, MCW1329, MCW1330, MCW1331, MW1357, MCW1373, MCW1375, MCW1385, MCW1408, MCW1473, MCW1550, MCW1581, MCW1584 were made in our lab. All *C. elegans* strains were kept at 20°C for both maintenance and experiment. All *C. elegans* were non-starved for at least 2 generations on NGM plates seeded with OP50 bacteria before any experiment. The detailed genotypes of each strain are listed in Supplementary Table 7.

The *E. coli* strain HT115 (DE3) was obtained from the Ahringer RNAi library. The *E. coli* strains OP50 and HB101 were obtained from the Caenorhabditis Genetics Center.

### Strain generation – Extrachromosomal array

MCW618 (raxEx190 [pie-1p::drp-1::tbb-2 3’UTR + myo-2p::GFP]) was generated by microinjecting the pie-1p::drp-1::tbb-2 3’UTR and myo-2p::GFP plasmids into the gonad of young adults. MCW1581 (raxEx618[pie-1p:: cox8(mitochondrial targeting sequence)::ndk-1::3xHA::pie-1 3’UTR + myo-2p::GFP]) was generated by microinjecting pie-1p::cox8(mitochondrial targeting sequence)::ndk-1::3xHA::pie-1 3’UTR PCR product and myo-2p::GFP plasmid into the gonad of young adults.

### Strain Generation – Integration of extrachromosomal array

MCW1220 (*raxIs141 [pie-1p::drp-1::tbb-2 3’UTR + myo-2p::GFP]*) was generated by the integration of extrachromosomal array in MCW618 which is induced by gamma irradiation exposures (4500rad, 5.9min) at the L4 stage. Later, the integrated progenies were backcrossed to N2 five times.

### Strain Generation – CRISPR-Cas9 mediated insertion and deletion

MCW1315 (*drp-1(rax82[GFP::Degron::drp-1]) IV*) was generated by inserting the Degron sequence into the *GFP::drp-1* locus of EU2917 between *GFP* and *drp-1* following the protocol from *Dokshin et al., 2018* with some modifications^76^. In short, a mixture of Cas9 protein (1.25μg/μl), tracrRNA (1μg/μl), target crRNA (0.4μg/μl), *dpy-10* crRNA (0.16μg/μl), and partially single-stranded DNA donor (300nM final concentration for each PCR product) was microinjected into the gonad of young adults. The partially single-stranded DNA donor was generated by mixing 2 PCR products – Degron sequence with 30 or 100 base pair homology arms on each side, and heat to 95°C then gradually cooling back to 20°C for melting and reannealing. After 3 days, the plates that have worms with *Dpy* phenotype were carefully chosen as jackpot plates for individualization of non-*Dpy* worms. These worms were subjected to pooled and then individual genotyping PCR after they reproduced to ensure passage of the genotype. The progenies (F2) of the specific F1 worm with the desired genotype were further individualized for identification of homozygosity using genotyping PCR and then sanger sequencing.

MCW1325*(sucg-1(rax83) IV)*, MCW1331(*sucg-1(rax86) IV)*, MCW1329(*suca-1(rax84) X)*, and MCW1330(*suca-1(rax85) X)* knockout or partial knockout strains were generated using methodologies described in *Chen et al., 2014* with modifications^77^. A mixture of Cas9 protein (1.25μg/μl), tracrRNA (1μg/μl), 2 target crRNAs (0.4μg/μl each) on 5’ and 3’ of a gene, and *dpy-10* crRNA (0.16μg/μl), were microinjected into the gonad of young adults. The screening process was the same as described for the knock-in strain MCW1315. MCW1329 and MCW1330 were backcrossed to N2 for three times.

MCW1408 *(raxIs89[sun-1p::eGFP::sun-1 3’UTR] III)* was generated by inserting *sun-1p::eGFP::sun-1 3’UTR* into ChrIII 7007.6. A mixture of Cas9 protein (1.25μg/μl), tracrRNA (1μg/μl), target crRNA (0.4μg/μl), and partially single-stranded DNA donor (10nM final concentration for each PCR product) was microinjected into the gonad of young adults. The partially single-stranded DNA donor was generated by mixing 2 PCR products - *sun-1p::eGFP::sun-1 3’UTR* sequence with 150bp of flanking homology arms on each side and the plain *sun-1p::eGFP::sun-1 3’UTR* sequence (both amplified using pYT17 plasmid as template), and heat to 95°C then gradually cool back to 20°C for melting and reannealing. Each injected worms were individualized post-injection. After 4 days, F1s were screened under fluorescence scope for green fluorescence in the germline. The progenies (F2) of the specific F1 worm with the desired genotype were further individualized for identification of homozygosity using fluorescence scope and then genotyping PCR followed by sanger sequencing.

MCW1473 *(raxIs98[sun-1p::eGFP::3xHA::sun-1 3’UTR] III)* was generated by inserting triple HA sequence between e*GFP* and *sun-1 3’UTR* at ChrIII 7007.6 position; *sun-1p::eGFP::sun-1 3’UTR* genetic locus in MCW1408. The experiment procedure was the same as generating MCW1315 except for the usage of single-strand oligodeoxynucleotides (with 30∼40nt homology arms on each side; 250ng/μl final concentration) instead of partially single-stranded DNA donor as the repair template, and melting and reannealing step by heating and cooling was not performed.

MCW1550 *(raxIs109[sun-1p::tomm-20(1-55aa)::eGFP::3xHA::sun-1 3’UTR] III)* as generated by inserting the first 165 nucleotides of *tomm-20* gene between *sun-1p* and *eGFP* at ChrIII 7007.6 position; *sun-1p::eGFP::3xHA::sun-1 3’UTR* genetic locus in MCW1473. The experiment procedure was the same as generating MCW1473. Later, MCW1550 was backcrossed to N2 for five times.

Genotyping PCR was performed using spanning primers for MCW1315, MCW1325, MCW1331, MCW1329, MCW1330, and MCW1408, and then followed by confirmation with sanger sequencing. For MCW1473 and MCW1550, genotyping PCR screen was performed using spanning primer on the 5’ and internal primer on the 3’, and the candidates were further verified using genotyping PCR by spanning primers followed by confirmation with sanger sequencing.

All primers used for genotyping are listed in Supplementary Table 5. Sequences of all crRNAs and the tracrRNA used for generating strains by CRISPR-Cas9 are listed in Supplementary Table 6.

### Strain Generation – Crossing

MCW1373 (*egxSi155 [mex-5p::tomm-20::mKate2::pie-1 3’UTR + unc-119(+)] II; unc-119(ed3) III; sucg-1(syb3617[sucg-1::eGFP]) IV*) was generated by crossing PHX3617 male to EGD629 hermaphrodite. eGFP^+^ F1s were selected to a population plate under the fluorescent scope, and the eGFP^+^ F2s on the population plate were then picked into individual plates. The F3s were later examined for green fluorescence, and individual plates with all eGFP^+^ (homozygous) F3 worms were then selected. Confocal imaging was then used to screen for the *tomm-20::mKate2* homozygous genotype, and genotyping PCR followed by sanger sequencing were used to examine the *unc-119* genotype.

MCW1326 (*ieSi68 [sun-1p::TIR1::mRuby::htp-1 3’UTR + Cbr-unc-119(+)] II; unc-119(ed3) III; drp-1(rax82[GFP::Degron::drp-1]) IV*) was generated by crossing MCW1315 male to CA1472 hermaphrodite. F1s were picked into individual plates, and then the *GFP::Degron::drp-1*; *TIR-1::mRuby* (heterozygous) genotype inspected by confocal imaging after egg laying. The F2s from F1 with the correct heterozygous genotype were then picked into individual plates. Later, F3s were later used to screen for the correct homozygous genotype of *GFP::Degron::drp-1*; *TIR-1::mRuby* by confocal imaging. Lastly, genotyping PCR followed by sanger sequencing were used to examine the *unc-119* genotype.

MCW1357 (raxIs141[pie-1p::drp-1.b::tbb-2 UTR + myo-2p::GFP]; egxSi152[mex5p::tomm-20::GFP::pie-1 3’UTR + unc-119(+)] II; unc-119(ed3) III) was generated by crossing EGD623 male to MCW1220 hermaphrodite. F1s were inspected for the mex5p::tomm-20::GFP::pie-1 3’UTR by the fluorescent microscope, and the worms with the correct (heterozygous) genotype were individualized. Later, the myo-2p::GFP^+^ F2s from F1 with the correct mex5p::tomm-20::GFP::pie-1 3’UTR heterozygous genotype were then picked into individual plates. Later, F3s were later used to screen for the correct homozygous genotype of myo-2p::GFP and mex5p::tomm-20::GFP::pie-1 3’UTR by fluorescence scope. Lastly, genotyping PCR followed by sanger sequencing were used to examine the unc-119 genotype.

MCW1584 (egxSi152[mex5p::tomm-20::GFP::pie-1 3’UTR + unc-119(+)] II; unc-119(ed3) III; drp-1(tm1108) IV) was generated by crossing EGD623 male to CU6372 hermaphrodite. F1s were then inspected for the mex5p::tomm-20::GFP::pie-1 3’UTR by fluorescent microscope, and the one with the correct (heterozygous) genotype were individualized. The F2s from F1 with the correct mex5p::tomm-20::GFP::pie-1 3’UTR heterozygous genotype were then picked into individual plates, and single worm lysed for drp-1(tm1108) PCR genotyping after egg laying. Later, F3s were used to screen for the correct homozygous genotype of mex5p::tomm-20::GFP::pie-1 3’UTR by fluorescent microscope. Lastly, genotyping PCR followed by sanger sequencing were used to examine the unc-119 genotype.

Genotyping PCR of for *drp-1(tm1108)* and *unc-119* was performed using spanning primers followed by confirmation with sanger sequencing. The primers used for *drp-1(tm1108)* and *unc-119* genotyping are listed in Supplementary Table 5.

MCW1375 (*sucg-1(syb3617[sucg-1::eGFP]); sucg-1(rax83) IV*) and MCW1385 (*sucg-1(syb3617[sucg-1::eGFP]); sucg-1(rax86) IV*) were obtained by crossing PHX3617 male to MCW1325 or MCW1331 hermaphrodites. eGFP^+^ F1s were picked under the fluorescent microscope and picked into individual plates. Later, F2s were used to confirm the *sucg-1::gfp/*KO heterozygous genotype of the F1 parental worms by fluorescent microscope (eGFP^+^/eGFP^−^ F2s should be around 3:1). Heterozygous genotypes were maintained by picked eGFP^+^ heterozygous worms (lower eGFP intensity than homozygous) for passage.

### RNA interference (RNAi) experiments

RNAi libraries created by the lab of Dr. Marc Vidal and Dr. Julie Ahringer were used in this study^78, 79^. *sucg-1, mev-1, sdhb-1, ogdh-1, drp-1, eat-3, and mmcm-1* RNAi clones were acquired from the Vidal library while *sucl-2, suca-1, and metr-1* RNAi clones were acquired from the Ahringer library. *fzo-1* RNAi clone was generated in the lab using L4440 as vector backbone and full-length *fzo-1* transcript as insert. All RNAi clones were verified by Sanger sequencing. For OP50 RNAi experiments, the genetically modified competent OP50 bacteria *[rnc14::DTn10 laczgA::T7pol camFRT]* generated by our lab (*Neve et al., 2019*) was used and transformed with 50 ng of the RNAi plasmid every time before the experiment^80^. All RNAi colonies were selected in both 50 µg ml^−1^ carbenicillin and 50 µg ml^−1^ tetracycline resistance. All RNAi bacteria were cultured for 14 hours in LB with 25 µg ml^−1^ carbenicillin, and then seeded onto RNAi agar plates that contain 1 mM IPTG and 50 µg ml^−1^ carbenicillin. The plates were then left at room temperature overnight for induction of dsRNA expression. For the RNAi experiments that require auxin treatment, fresh bacteria were concentrated 4 times before seeding onto the plates, and then left in 4°C overnight before usage.

### Construction of plasmid and fusion PCR product

The *pie-1p::drp-1::tbb2 3’UTR* plasmid was generated by PCR amplifying the complete coding sequence of *drp-1.b* transcript from N2 cDNA and utilized Gateway BP recombination to clone into pDONR221 which contains Gateway attLR recombination sequences. *drp-1.b* CDS entry clone was then recombined with the entry clones pCM1.36-*tbb-2* 3’UTR and pCM1.127-*pie-1p* into destination vector pCFJ150 using Gateway LR recombination.

The pie-1p::cox8(mitochondrial targeting sequence)::ndk-1::3xHA::pie-1 3’UTR oligonucleotide was generated by 3-fragment fusion PCR using cox8(mitochondrial targeting sequence)::ndk-1::3xHA, pie-1p, and pie-1 3’UTR PCR product. cox8(mitochondrial targeting sequence)::ndk-1::3xHA sequence was synthesized by IDT, and utilized as the template for amplification and homology arm tagging (tagged with pie-1p and pie-1 3’UTR homologies on 5’ and 3’ end respectively). Both pie-1p and pie-1 3’UTR PCR products were amplified using pPK605 plasmid (Addgene) as the template.

The pYT17-*sun-1p::eGFP::sun-1 3’UTR* plasmid was generated via 4-fragment Gibson cloning from vector backbone, *sun-1p*, modified *eGFP*, and *sun-1 3’UTR* PCR products. *sun-1p* and *sun-1 3’UTR* PCR products were amplified using N2 worm lysate as the template. Modified *eGFP* PCR product was amplified using PHX3617 worm lysate as the template.

Primers used for the amplification are listed in Supplementary Table 5.

### Reproductive lifespan assay

Synchronized L1 larvae from egg preparation were plated onto 6cm NGM plates seeded with the specific bacteria (default: HT115) and grew to L4 stage before being individualized into single 3cm NGM plates. The worms were transferred to a new plate every day except for the day right after individualization, which we collectively (L4 + day-1-old adult) count as day 1. The transferring stopped when we observed 2 days of non-reproducing events consecutively or until day 12. After each transfer, plates were stored at room temperature for 2 days before checking the reproductive status. The last day of progeny production was counted as the day of reproductive cessation, and worms that could not be tracked until the day of reproductive cessation due to missing, death, germline protrusion, or internal hatching were counted as censors on the last day which we could determine the reproductive status. The animals were removed from the analysis if they died before producing any progeny. Statistical analyses were performed in SPSS software using Kaplan-Meier survival method followed by a log-rank test.

For RLS experiments of MCW1581 *(raxEx618[pie-1p:: cox8(mitochondrial targeting sequence)::ndk-1::3xHA::pie-1 3’UTR + myo-2p::GFP])*, day 1 *myo-2p::GFP*^+^ F1s of injected parental worms were individually picked onto EV or *sucg-1* RNAi plates. 3 and 4 days later, the plates with *myo-2p::GFP*^+^ F2s were selected, and the same number of *myo-2p::GFP*^+^ and *myo-2p::GFP*^−^ F2 worms at L4 stage were picked from each population plate into individual EV or *sucg-1* RNAi plates. The later part of the RLS methodology follows the protocol above.

For RLS experiments of MCW1375 and MCW1385 strains, heterozygous parental worms were individualized onto the 6cm NGM plates at day 1 adulthood and the plates were kept for 4 days. The genotypes of the parental worms were then examined by the eGFP phenotypes in F1 under the fluorescent scope to ensure heterozygosity (of the parental line), and F1 progenies at L4 stage were randomly picked and individualized onto 3cm NGM plates. The later part of the RLS methodology follows protocol above, with an additional step of examining the genotype of each F1 worm by observing the eGFP phenotypes in F2s.

### Late fertility assay

Synchronized L1 larvae from egg preparation were plated on 6cm NGM plates seeded with the specific bacteria (default: HT115) and transferred every 2 days to new NGM plates from L4 until day 9. Individual hermaphrodites were transferred to a 3cm NGM plates seeded with OP50 bacteria together with 2 day-2-old young N2 males for mating. Hermaphrodites were mated for 2 days before the first round of examination, which will exclude the plates with dead hermaphrodites, germline protruded hermaphrodites, or 2 dead males. The plates were then kept for one more day until the second-round examination of progeny production. Unlike RLS, internal hatched worms were not censored but instead considered as a reproduction event in late fertility assay. 15-20 hermaphrodites were used for each experiment and was repeated at least 3 times independently to reach 60 worms per condition (before exclusion). The results from different trials were then pooled to conduct Fisher’s exact test to determine whether the number of worms that resumed reproduction after mating in each condition is significantly different from the controls.

### Confocal imaging

Sample preparations were done by anesthetizing the worms in 1% sodium azide (NaAz) in M9 buffer, mounted on 2% agarose pads on glass slides, and covered the pads with coverslips. The worms were then imaged on laser scanning confocal FV3000 (Olympus, US) with water immersion 60x objective (UPLSAPO 60XW, Olympus, US) for SUCG-1 mitochondrial localization in the germline, germline morphology and mitochondrial localization of day 5 worms subjected to *drp-1* RNAi knockdown, and oocyte mitochondrial distribution. 20x objective (UPLSAPO 20X, Olympus, US) was used for assessing the expression pattern of SUCG-1::eGFP and SUCA-1::eGFP, and intensity of SUCG-1::eGFP on day 1 and day 5. 10X objective (UPlanFL N 10X, Olympus, US) was used to measure the body length of worms subjected to EV or *eat-3* germline-specific RNAi knockdown.

### Germline fluorescent intensity profiling

The images of the germline SUCG-1::eGFP were generated by 20x z-stacked confocal imaging of PHX3617 strain. For a given 3D image stack of eGFP labeled germline, the max intensity at each (x,y) location was projected to a single image, i_max_. Multiple polygons p_1_, p_2_,…, p_m_ (m is the number of imaged germlines) were manually selected on i_max_ to outline germlines. A 2D mask m_i_ was generated for each p_i_, with i =1, 2,…, m. m_i_ was extended to 3D mask v_i_ by multiplying the depth of the stack and then use the v_i_ to selected 3D region for calculation total and average intensity of eGFP. The region selected spans from the proliferation zone to the mid-point of U-shaped loop due to technical difficulties of consistently getting quality image of the entire germline and the blurred border between oocyte and spermatheca in aged worms. All analyses above were done using MATLAB. Student’s test was used to determine whether the eGFP intensities of day-5-old worms are statistically distinct from the day-1-old worms. The code for the analyses is provided in Supplementary File 1.

### Analysis of oocyte mitochondrial network

The images of the oocyte mitochondrial network were generated by 60x confocal imaging of EGD623 strain or mutant and integrated strains crossed with EGD623, and the position −2 oocytes were used for downstream analysis. Stacked oocytes with little distance between the nuclear membrane and the lateral side of the plasma membrane were excluded from the analysis.

For code-based radial intensity profiling of oocyte mitochondrial network, mitochondrial distribution as their distance from cell nucleus was quantified by generating two masks using manual selection with polygon on the DIC images - polygon p_1_ outlining cell nucleus and p_2_ outlining cell body. A set of rays were calculated with their origins at the mass center of p_1_. The rays were customized to cover 360° with a step size of 1°. Each ray intersected with p_1_ and p_2_ and got a line segment. All line segments were divided into 5 equal segments, and labeled as ls_1_, ls_2_,…, ls_5_, starting from the segment closest to cell nucleus. All ends of ls_1_ were connected to get a ring shape r_1_, and then the same for ls_2_ to ls_5_ resulting in r_2_ to r_5_. These rings were used as mask to select regions in an oocyte for mitochondrial intensity calculation leading to generation of a radial mitochondrial distribution. All the above analyses were done using MATLAB. The code for the analyses is provided in Supplementary File 2.

Later, the ring 1 occupancy of each oocyte was converted into one of the three categories using the following cutoffs – dispersed when lower than 23.5%, intermediate when equal or higher than 23.5% but lower than 26.5%, and perinuclear when equal or higher than 26.5%. The cutoffs were defined through double-blind categorization. Chi-squared test was then used to determine whether the oocyte mitochondrial distribution of each condition is significantly different from the control.

### Germline mtDNA levels measurement by quantitative PCR (qPCR)

Around 30 germlines were dissected for each condition following the protocol from *Gervaise et al., 2016*^81^. After dissection, germlines in M9 solution were collected into a PCR tube with a glass Pasteur pipette, and centrifuged at 15000rpm for 2 minutes. Later, the excess M9 solution was removed from the PCR tube, and worm lysis buffer was added. The PCR tube was then placed at −80°C for at least 15 minutes before incubating at 60°C for 60 minutes followed by 95°C for 15 minutes for lysis and DNA release. qPCR was then performed using Power SYBR green master mix (Applied Biosystems *#4367659*) in a realplex 4 qPCR cycler (Eppendorf). To calculate the relative mtDNA levels, the cycle number of *nduo-1* and *ctb-1* (both encoded by mitochondrial DNA) were normalized to *ant-1.3* (encoded by genomic DNA).

### Body length measurement

The DIC channel on confocal microscopy was used to image the full body lengths of day 1 worms subjected to EV or *eat-3* germline-specific RNAi knockdown side by side. The images were then analyzed using ImageJ by drawing segmented lines spanning head to tail of the worms, which was then followed by distance measurement.

### Pharyngeal pumping measurement

A digital camera (ORCA-Flash4.0 LT, Hamamatsu) attached to the stereoscope was used to record the pharyngeal pumping rate of worms subjected to EV or *eat-3* germline-specific RNAi knockdown. After recording, the movies were played at 0.25X speed, and the times of pharyngeal pumping in each second (pumping rate) were counted. For each worm, the average pumping rate in 5-10 seconds was used for analysis.

### Auxin treatment

Auxin (Alfa Aesar #A10556) was administered to the *C. elegans* using methodologies described in *Zhang et al., 2015* with slight modification^44^. A 400mM auxin stock solution in ethanol was prepared and filtered through a 0.22µm filter, which was stored at 4°C for up to 2 weeks. Auxin stock solution was added into the NGM liquid agar with a concentration of 1 to 100 (1%) after the autoclaved liquid agar drops below 50°C and then poured into the plates making a final auxin concentration of 4mM. For the control plates, filtered ethanol was added to the NGM liquid agar with a concentration of 1 to 100 (1%). The plates were stored at 4°C inside a box with low photopermeability after the agar solidified. Before usage, fresh bacteria were concentrated by 4X before seeding onto the plates, and the plates that weren’t used immediately were stored at 4°C for up to 5 days.

### Germline mitochondrial GTP and ATP measurement

Synchronized MCW1550 L1 larvae from egg preparation were plated onto 15cm NGM plates seeded with the 20X concentrated bacteria and grew to day 1. The worms were then harvested (day 1 sample) or filtered daily (filter out eggs and progenies) using a 40µm cell strainer and seeded onto new 15cm NGM plate until day 5 before getting harvested (day 5 sample). Approximately 50k worms were used for day 5 sample collection and 100k worms were used for day 1 sample collection.

Germline mitochondria isolation was performed using methodologies described in *Ahier et al., 2018*^82^ with modifications. In short, worms were harvested into a 15cm centrifuge tube, and washed 3 times with 10ml M9 buffer and then 2 more times with cold KPBS buffer (136mM KCl, 10mM KH_2_PO_4_, pH = 7.2). The worms were then transferred to a dauncer on ice and daunced until most worms were clearly broken. Later, the lysates were transferred into a centrifuge tube for low-speed centrifugation to precipitate large fragments, and the supernatant containing the organelles was then collected and centrifuged again at high speed to precipitate the organelles. The pellet was resuspended in KPBS buffer, anti-HA magnetic beads (Pierce #88837) were added, and the tube was incubated at 4°C for an hour to ensure binding efficiency. The anti-HA magnetic beads were then washed three times with KPBS, portioned out for protein concentration measurement by BCA assay and mitochondrial DNA content detection by qPCR, and the remaining beads were stored at −80°C for later steps of GTP and ATP detection.

For detection of nucleotides, immunoprecipitated mitochondria (with around 100 to 200μg mitochondria protein) were resuspended in pre-chilled water to the concentration of 1μg mitochondria protein per μl water. 500μl pre-chilled chloroform was then immediately added to the resuspended mitochondria samples, followed by vigorous vortexing to quench metabolism and to extract soluble metabolites. The mitochondria extracts were centrifuged at 20,000g for 10min at 4°C to remove the organic phase, followed by another centrifugation at 20,000g for 10min at 4°C to remove cell debris. The resulting supernatants were diluted 10 times (to 0.1μg mitochondria protein per μl water) and analyzed immediately using HPLC-MS as described previously^83, 84^.

Data analysis was performed using the Metabolomics Analysis and Visualization Engine (MAVEN) software^85^. For each sample, ion counts of nucleotides were normalized to mitochondrial protein mass concentration followed by mtDNA (*nduo-1*) level. All samples were then normalized to the (HT115 bacteria; D1) condition to indicate fold changes.

### Cobalamin treatment

Methylcobalamin (Sigma-Aldrich #M9756) and adenosylcobalamin (Sigma-Aldrich #C0884) were administered to the *C. elegans* using methodologies similar to auxin treatment. A 1.28mM aqueous stock solution was freshly prepared and filtered through a 0.22µm filter. The stock solution was added into the NGM liquid agar with a concentration of 1 to 10000 (0.01%) after the autoclaved liquid agar drops below 50°C and then poured into the plates making a final cobalamin concentration of 128nM. For the control plates, filtered double-distilled water was added to the NGM liquid agar instead. The plates were stored at 4°C inside a box with low photopermeability after the agar solidified. Bacteria were seeded before usage, and the plates that weren’t used immediately were stored at 4°C for up to 5 days.

### Succinate treatment

Sodium succinate (Sigma Aldrich #S2378) and succinic acid (Thermo Scientific Chemicals #AA3327236) were administered to the *C. elegans* via supplementation into the NGM plates. Precalculated amounts of sodium succinate and succinic acid were added into the liquid agar right after being taken out from the autoclave to make 10mM final concentration, and the agar was then poured into the plates after cooling down. The plates were stored at 4°C inside a box with low photopermeability after the agar solidified. Bacteria were seeded before usage.

## QUANTIFICATION AND STATISTICAL ANALYSIS

The reproductive lifespan analyses were performed using Kaplan-Meier survival analysis and a log-rank test in the SPSS. Chi-squared tests and Fisher’s exact tests were performed in Graphpad PRISM to compare categorical variables, and Holm-Bonferroni method was used for correction as indicated in the corresponding figure legends. Student’s t-test (unpaired) were performed in Excel to compare the mean of different samples, and Holm-Bonferroni method was used for correction as indicated in the corresponding figure legends. For all figure legends, asterisks indicate statistical significance as follows: n.s. = not significant *p>0.05*; *\* p<0.05*; *\*\* p<0.01*; *\*\*\* p<0.001*. Data were collected from at least three independent biological replicates. Figures and graphs were constructed using BioRender, PRISM, and Illustrator.

## ACKNOWLEDGMENT

This work was supported by NIH grants R01AG045183 (M.C.W.), R01AT009050 (M.C.W.), R01AG062257 (M.C.W.), DP1DK113644 (M.C.W.), March of Dimes Foundation (M.C.W.), Welch Foundation (M.C.W.), HHMI investigator (M.C.W.), American Federation for Aging Research (Y.L.). We thank P. Svay for maintenance support and C. Huang for technical support. We thank I. Neve and H. Oakley for conducting preliminary screening of this study. We thank Dr. Bruce Bowerman for providing the *drp-1* endogenous locus sequence information of the EU2917 strain. We thank BioRender for the support on creating Fig 1A, Fig 3A, and Fig 4A. We thank the Caenorhabditis Genetics Center (CGC) for *C. elegans* strains.

## AUTHOR CONTRIBUTIONS

Y.L., J.S., and M.C.W. conceived the project. Y.L., M.Savini., T.C., J.Y., Q.Z., L.D., M.Senturk, and J.S. performed experiments. T.C. and S.G. wrote the code for imaging analysis. Y.L and M.C.W. wrote the manuscript. Y.L., J.J.W. and M.C.W. edited the manuscript.

## Notes

### Competing Interest Statement

The authors have declared no competing interest.

## REFERENCES

1. Hamilton, B., Martin, J. & Osterman, M. Births: Provisional Data for 2020. https://stacks.cdc.gov/view/cdc/104993 (2021) doi:10.15620/cdc:104993.

2. Duncan, F. E. & Gerton, J. L. Mammalian oogenesis and female reproductive aging. Aging (Albany NY) 10, 162–163 (2018).

3. te Velde, E. R. The variability of female reproductive ageing. Human Reproduction Update 8, 141–154 (2002).

4. Babayev, E. & Seli, E. Oocyte mitochondrial function and reproduction. Curr Opin Obstet Gynecol 27, 175–181 (2015).

5. May-Panloup, P., Boguenet, M., El Hachem, H., Bouet, P.-E. & Reynier, P. Embryo and Its Mitochondria. Antioxidants 10, 139 (2021).

6. Wilding, M. et al. Mitochondrial aggregation patterns and activity in human oocytes and preimplantation embryos. Human Reproduction 16, 909–917 (2001).

7. Reynier, P. et al. Mitochondrial DNA content affects the fertilizability of human oocytes. Mol Hum Reprod 7, 425–429 (2001).

8. Van Blerkom, J., Davis, P. W. & Lee, J. ATP content of human oocytes and developmental potential and outcome after in-vitro fertilization and embryo transfer. Hum Reprod 10, 415– 424 (1995).

9. Detmer, S. A. & Chan, D. C. Functions and dysfunctions of mitochondrial dynamics. Nat Rev Mol Cell Biol 8, 870–879 (2007).

10. Udagawa, O. et al. Mitochondrial Fission Factor Drp1 Maintains Oocyte Quality via Dynamic Rearrangement of Multiple Organelles. Current Biology 24, 2451–2458 (2014).

11. Wakai, T., Harada, Y., Miyado, K. & Kono, T. Mitochondrial dynamics controlled by mitofusins define organelle positioning and movement during mouse oocyte maturation. MHR: Basic science of reproductive medicine 20, 1090–1100 (2014).

12. Hou, X. et al. Mitofusin1 in oocyte is essential for female fertility. Redox Biology 21, 101110 (2019).

13. Zhang, M. et al. Mitofusin 1 is required for female fertility and to maintain ovarian follicular reserve. Cell Death Dis 10, 560 (2019).

14. Han, B. et al. Microbial Genetic Composition Tunes Host Longevity. Cell 169, 1249–1262.e13 (2017).

15. Weir, H. J. et al. Dietary Restriction and AMPK Increase Lifespan via Mitochondrial Network and Peroxisome Remodeling. Cell Metab 26, 884–896.e5 (2017).

16. Rana, A. et al. Promoting Drp1-mediated mitochondrial fission in midlife prolongs healthy lifespan of Drosophila melanogaster. Nat Commun 8, 448 (2017).

17. Luo, S., Kleemann, G. A., Ashraf, J. M., Shaw, W. M. & Murphy, C. T. TGF-β and Insulin Signaling Regulate Reproductive Aging via Oocyte and Germline Quality Maintenance. Cell 143, 299–312 (2010).

18. Hughes, S. E., Evason, K., Xiong, C. & Kornfeld, K. Genetic and Pharmacological Factors That Influence Reproductive Aging in Nematodes. PLOS Genetics 3, e25 (2007).

19. Sowa, J. N., Mutlu, A. S., Xia, F. & Wang, M. C. Olfaction Modulates Reproductive Plasticity through Neuroendocrine Signaling in Caenorhabditis elegans. Curr. Biol. 25, 2284–2289 (2015).

20. Wang, M. C., Oakley, H. D., Carr, C. E., Sowa, J. N. & Ruvkun, G. Gene Pathways That Delay Caenorhabditis elegans Reproductive Senescence. PLOS Genetics 10, e1004752 (2014).

21. Martínez-Reyes, I. & Chandel, N. S. Mitochondrial TCA cycle metabolites control physiology and disease. Nat Commun 11, 102 (2020).

22. Johnson, J. D., Muhonen, W. W. & Lambeth, D. O. Characterization of the ATP- and GTP-specific Succinyl-CoA Synthetases in Pigeon: THE ENZYMES INCORPORATE THE SAME α-SUBUNIT *. Journal of Biological Chemistry 273, 27573–27579 (1998).

23. Fraser, M. E., Hayakawa, K., Hume, M. S., Ryan, D. G. & Brownie, E. R. Interactions of GTP with the ATP-grasp Domain of GTP-specific Succinyl-CoA Synthetase *. Journal of Biological Chemistry 281, 11058–11065 (2006).

24. Lambeth, D. O., Tews, K. N., Adkins, S., Frohlich, D. & Milavetz, B. I. Expression of Two Succinyl-CoA Synthetases with Different Nucleotide Specificities in Mammalian Tissues *. Journal of Biological Chemistry 279, 36621–36624 (2004).

25. Kibbey, R. G. et al. Mitochondrial GTP Regulates Glucose-Stimulated Insulin Secretion. Cell Metabolism 5, 253–264 (2007).

26. Jesinkey, S. R. et al. Mitochondrial GTP Links Nutrient Sensing to β Cell Health, Mitochondrial Morphology, and Insulin Secretion Independent of OxPhos. Cell Reports 28, 759–772.e10 (2019).

27. Zhao, Y. et al. Loss of succinyl-CoA synthase ADP-forming β subunit disrupts mtDNA stability and mitochondrial dynamics in neurons. Sci Rep 7, 7169 (2017).

28. Wu, B. et al. Succinyl-CoA Ligase Deficiency in Pro-inflammatory and Tissue-Invasive T Cells. Cell Metabolism 32, 967–980.e5 (2020).

29. Zou, L. et al. Construction of a germline-specific RNAi tool in C. elegans. Sci Rep 9, 2354 (2019).

30. Gems, D. et al. Two pleiotropic classes of daf-2 mutation affect larval arrest, adult behavior, reproduction and longevity in Caenorhabditis elegans. Genetics 150, 129–155 (1998).

31. Ng, L. T. et al. Lifespan and healthspan benefits of exogenous H2S in C. elegans are independent from effects downstream of eat-2 mutation. NPJ Aging Mech Dis 6, 6 (2020).

32. Luo, S., Shaw, W. M., Ashraf, J. & Murphy, C. T. TGF-beta Sma/Mab signaling mutations uncouple reproductive aging from somatic aging. PLoS Genet 5, e1000789 (2009).

33. Lakowski, B. & Hekimi, S. The genetics of caloric restriction in Caenorhabditis elegans. Proc Natl Acad Sci U S A 95, 13091–13096 (1998).

34. Kenyon, C., Chang, J., Gensch, E., Rudner, A. & Tabtiang, R. A C. elegans mutant that lives twice as long as wild type. Nature 366, 461–464 (1993).

35. Fan, X. et al. SapTrap Assembly of Caenorhabditis elegans MosSCI Transgene Vectors. G3 (Bethesda) 10, 635–644 (2019).

36. Boissan, M., et al. Membrane trafficking. Nucleoside diphosphate kinases fuel dynamin superfamily proteins with GTP for membrane remodeling. Science 344, 1510–1515 (2014).

37. Westermann, B. Mitochondrial dynamics in model organisms: What yeasts, worms and flies have taught us about fusion and fission of mitochondria. Seminars in Cell & Developmental Biology 21, 542–549 (2010).

38. Kanazawa, T. et al. The C. elegans Opa1 Homologue EAT-3 Is Essential for Resistance to Free Radicals. PLOS Genetics 4, e1000022 (2008).

39. Ichishita, R. et al. An RNAi screen for mitochondrial proteins required to maintain the morphology of the organelle in Caenorhabditis elegans. J Biochem 143, 449–454 (2008).

40. Avery, L. The genetics of feeding in Caenorhabditis elegans. Genetics 133, 897–917 (1993).

41. Breckenridge, D. G. et al. Caenorhabditis elegans drp-1 and fis-2 regulate distinct cell-death execution pathways downstream of ced-3 and independent of ced-9. Mol Cell 31, 586–597 (2008).

42. Tan, F. J. et al. CED-9 and mitochondrial homeostasis in C. elegans muscle. J Cell Sci 121, 3373–3382 (2008).

43. Lowry, J. et al. High-Throughput Cloning of Temperature-Sensitive Caenorhabditis elegans Mutants with Adult Syncytial Germline Membrane Architecture Defects. G3 (Bethesda) 5, 2241–2255 (2015).

44. Zhang, L., Ward, J. D., Cheng, Z. & Dernburg, A. F. The auxin-inducible degradation (AID) system enables versatile conditional protein depletion in C. elegans. Development 142, 4374–4384 (2015).

45. Watson, E. et al. Interspecies systems biology uncovers metabolites affecting C. elegans gene expression and life history traits. Cell 156, 759–770 (2014).

46. Wei, W. & Ruvkun, G. Lysosomal activity regulates Caenorhabditis elegans mitochondrial dynamics through vitamin B12 metabolism. Proc Natl Acad Sci U S A 117, 19970–19981 (2020).

47. Revtovich, A. V., Lee, R. & Kirienko, N. V. Interplay between mitochondria and diet mediates pathogen and stress resistance in Caenorhabditis elegans. PLoS Genet 15, e1008011 (2019).

48. Ghergurovich, J. M. et al. Methionine synthase supports tumour tetrahydrofolate pools. Nat Metab 3, 1512–1520 (2021).

49. Sullivan, M. R. et al. Methionine synthase is essential for cancer cell proliferation in physiological folate environments. Nat Metab 3, 1500–1511 (2021).

50. Cota, V., Sohrabi, S., Kaletsky, R. & Murphy, C. T. Oocyte mitophagy is critical for extended reproductive longevity. PLoS Genet 18, e1010400 (2022).

51. Baixauli, F. et al. The mitochondrial fission factor dynamin-related protein 1 modulates T-cell receptor signalling at the immune synapse. EMBO J 30, 1238–1250 (2011).

52. Hennings, T. G. et al. In Vivo Deletion of β-Cell Drp1 Impairs Insulin Secretion Without Affecting Islet Oxygen Consumption. Endocrinology 159, 3245–3256 (2018).

53. Griparic, L., van der Wel, N. N., Orozco, I. J., Peters, P. J. & van der Bliek, A. M. Loss of the intermembrane space protein Mgm1/OPA1 induces swelling and localized constrictions along the lengths of mitochondria. J Biol Chem 279, 18792–18798 (2004).

54. Pich, S. et al. The Charcot-Marie-Tooth type 2A gene product, Mfn2, up-regulates fuel oxidation through expression of OXPHOS system. Hum Mol Genet 14, 1405–1415 (2005).

55. Shen, T. et al. Mitofusin-2 is a major determinant of oxidative stress-mediated heart muscle cell apoptosis. J Biol Chem 282, 23354–23361 (2007).

56. Rojo, M., Legros, F., Chateau, D. & Lombès, A. Membrane topology and mitochondrial targeting of mitofusins, ubiquitous mammalian homologs of the transmembrane GTPase Fzo. J Cell Sci 115, 1663–1674 (2002).

57. Santel, A. & Fuller, M. T. Control of mitochondrial morphology by a human mitofusin. J Cell Sci 114, 867–874 (2001).

58. Al-Mehdi, A.-B. et al. Perinuclear mitochondrial clustering creates an oxidant-rich nuclear domain required for hypoxia-induced transcription. Sci Signal 5, ra47 (2012).

59. Agarwal, S. & Ganesh, S. Perinuclear mitochondrial clustering, increased ROS levels, and HIF1 are required for the activation of HSF1 by heat stress. J Cell Sci 133, jcs245589 (2020).

60. Kim, S. et al. Hepatitis B virus x protein induces perinuclear mitochondrial clustering in microtubule- and Dynein-dependent manners. J Virol 81, 1714–1726 (2007).

61. Hu, M. et al. Respiratory syncytial virus co-opts host mitochondrial function to favour infectious virus production. Elife 8, e42448 (2019).

62. Pucci, B. et al. Detailing the role of Bax translocation, cytochrome c release, and perinuclear clustering of the mitochondria in the killing of HeLa cells by TNF. J Cell Physiol 217, 442– 449 (2008).

63. Dewitt, D. A. et al. Peri-nuclear clustering of mitochondria is triggered during aluminum maltolate induced apoptosis. J Alzheimers Dis 9, 195–205 (2006).

64. Thomas, W. D., Zhang, X. D., Franco, A. V., Nguyen, T. & Hersey, P. TNF-related apoptosis-inducing ligand-induced apoptosis of melanoma is associated with changes in mitochondrial membrane potential and perinuclear clustering of mitochondria. J Immunol 165, 5612–5620 (2000).

65. Geisler, S. et al. PINK1/Parkin-mediated mitophagy is dependent on VDAC1 and p62/SQSTM1. Nat Cell Biol 12, 119–131 (2010).

66. Mishra, P. & Chan, D. C. Metabolic regulation of mitochondrial dynamics. J Cell Biol 212, 379–387 (2016).

67. Bulutoglu, B., Garcia, K. E., Wu, F., Minteer, S. D. & Banta, S. Direct Evidence for Metabolon Formation and Substrate Channeling in Recombinant TCA Cycle Enzymes. ACS Chem. Biol. 11, 2847–2853 (2016).

68. Hatefi, Y. [6] Resolution of complex II and isolation of succinate dehydrogenase (EC 1.3.99.1). in Methods in Enzymology (eds. Fleischer, S. & Packer, L.) vol. 53 27–35 (Academic Press, 1978).

69. Meeusen, S., McCaffery, J. M. & Nunnari, J. Mitochondrial Fusion Intermediates Revealed in Vitro. Science 305, 1747–1752 (2004).

70. Schlattner, U. et al. Dual function of mitochondrial Nm23-H4 protein in phosphotransfer and intermembrane lipid transfer: a cardiolipin-dependent switch. J Biol Chem 288, 111–121 (2013).

71. MacNeil, L. T., Watson, E., Arda, H. E., Zhu, L. J. & Walhout, A. J. M. Diet-induced developmental acceleration independent of TOR and insulin in C. elegans. Cell 153, 240– 252 (2013).

72. Qin, S. et al. Early-life vitamin B12 orchestrates lipid peroxidation to ensure reproductive success via SBP-1/SREBP1 in Caenorhabditis elegans. Cell Rep 40, 111381 (2022).

73. Raghavan, R., et al. Maternal Multivitamin Intake, Plasma Folate and Vitamin B12 Levels and Autism Spectrum Disorder Risk in Offspring. Paediatr Perinat Epidemiol 32, 100–111 (2018).

74. Molloy, A. M., Kirke, P. N., Brody, L. C., Scott, J. M. & Mills, J. L. Effects of folate and vitamin B12 deficiencies during pregnancy on fetal, infant, and child development. Food Nutr Bull 29, S101–111; discussion S112-115 (2008).

75. Green, R. et al. Vitamin B12 deficiency. Nat Rev Dis Primers 3, 17040 (2017).

76. Dokshin, G. A., Ghanta, K. S., Piscopo, K. M. & Mello, C. C. Robust Genome Editing with Short Single-Stranded and Long, Partially Single-Stranded DNA Donors in Caenorhabditis elegans. Genetics 210, 781–787 (2018).

77. Chen, X. et al. Dual sgRNA-directed gene knockout using CRISPR/Cas9 technology in Caenorhabditis elegans. Sci Rep 4, 7581 (2014).

78. Rual, J.-F. et al. Toward Improving Caenorhabditis elegans Phenome Mapping With an ORFeome-Based RNAi Library. Genome Res 14, 2162–2168 (2004).

79. Kamath, R. Genome-wide RNAi screening in Caenorhabditis elegans. Methods 30, 313–321 (2003).

80. Neve, I. A. A. et al. Escherichia coli Metabolite Profiling Leads to the Development of an RNA Interference Strain for Caenorhabditis elegans. G3 (Bethesda) 10, 189–198 (2020).

81. Gervaise, A. L. & Arur, S. Spatial and Temporal Analysis of Active ERK in the C. elegans Germline. J Vis Exp 54901 (2016) doi:10.3791/54901.

82. Ahier, A. et al. Affinity purification of cell-specific mitochondria from whole animals resolves patterns of genetic mosaicism. Nat Cell Biol 20, 352–360 (2018).

83. Fung, D. K., Yang, J., Stevenson, D. M., Amador-Noguez, D. & Wang, J. D. Small Alarmone Synthetase SasA Expression Leads to Concomitant Accumulation of pGpp, ppApp, and AppppA in Bacillus subtilis. Front Microbiol 11, 2083 (2020).

84. Yang, J. et al. The nucleotide pGpp acts as a third alarmone in Bacillus, with functions distinct from those of (p) ppGpp. Nat Commun 11, 5388 (2020).

85. Clasquin, M. F., Melamud, E. & Rabinowitz, J. D. LC-MS data processing with MAVEN: a metabolomic analysis and visualization engine. Curr Protoc Bioinformatics Chapter 14, Unit14.11 (2012).

