## Supplementary File 1 for "Mitochondrial GTP Metabolism Regulates Reproductive Aging"

This script is made to automatically quantify the fluorescence of GFP labeled mitochondria in live C. elegans imaging. The three-dimensional quantification of GFP is performed based on masks to be applied to each germline. fluorescence is quantified in following steps. 1, Manually or automatically make mask that will select areas of germlime only 2, Apply the mask to each frame of the three-dimensional image stack 3, Quantify fluorescence intensity within each three-dimensional selected volumes The bioformat package is used to read the image file of factory format. Melissa Linkert, Curtis T. Rueden, Chris Allan, Jean-Marie Burel, Will Moore, Andrew Patterson, Brian Loranger, Josh Moore, Carlos Neves, Donald MacDonald, Aleksandra Tarkowska, Caitlin Sticco, Emma Hill, Mike Rossner, Kevin W. Eliceiri, and Jason R. Swedlow (2010) Metadata matters: access to image data in the real world. The Journal of Cell Biology 189(5), 777-782. doi: 10.1083/jcb.201004104

### Contents


- getting file
- selection of image
- selection of nucleus
- saving to files


### getting file

```
clear;
[file,path] = uigetfile('*.oir','select a oir file');
filename = fullfile(path,file);
meta = stackread(filename);
GFP = meta(:,:,1:2:end);
TD = meta(:,:,2:2:end);
dim = size(GFP);
alphab = 'ABCDEFGHIJKLMNOPQRSTUVWXYZ';
```

### selection of image

```
premask = max(GFP,[],3);
factor = 255/double(max(GFP(:)))*2;
i = 1;
choice = 'Yes';
```

### selection of nucleus

```
while(strcmp(choice,'Yes'))
    if strcmp(choice,'Yes')
        [mask{i},x{i},y{i}]=roipoly(adapthisteq(premask/factor));
    end
    i = i+1;
    dlgTitle = 'Germline selection';
    dlgQuestion = 'Continue to select?';
    choice = questdlg(dlgQuestion,dlgTitle,'Yes','No', 'Yes');
end
```

```
N = length(mask);
total_intensity = zeros(1,N);
volume = zeros(1,N);
for i = 1:1:N
    germ = maskapp(GFP,mask{i});
```

```
%     z_prof = squeeze(mean(germ,[1 2])); more precise z determination

    total_intensity(i) = sum(germ(:));
    volume(i) = sum(mask{i}(:)*dim(3));
end
avg = total_intensity./volume;
```

### saving to files

add title to each column and create the file if not existing

```
for i = 1:1:N
    header{i} = ['mean_g',num2str(i)];
    header{i+N} = ['total_g',num2str(i)];
    header{i+2*N} = ['size_g',num2str(i)];
end

if ~isfile('problem.xls')
    writecell({filename},'problem.xls');
    temp = readcell('problem.xls');temp = size(temp); temp = temp(1);
    range = ['A',num2str(temp+1),':',alphab(3*N),num2str(temp+1)];
    writecell(header,'problem.xls','Range',range);
else
    temp = readcell('problem.xls');temp = size(temp); temp = temp(1);
    range = ['A',num2str(temp+1)];
    writecell({filename},'problem.xls','Range',range);
    temp = readcell('problem.xls');temp = size(temp); temp = temp(1);
    range = ['A',num2str(temp+1),':',alphab(3*N),num2str(temp+1)];
    writecell(header,'problem.xls','Range',range);
end
temp = readcell('problem.xls');temp = size(temp); temp = temp(1);
range = ['A',num2str(temp+1),':',alphab(3*N),num2str(temp+1)];
writecell({avg,total_intensity,volume},'problem.xls','Range',range);

% saving polygon coordinates
% add title to each column and create the file if not existing
for i = 1:1:N
    header{i} = ['worm',num2str(i),'-x'];
    header{i+N} = ['worm',num2str(i),'-y'];
end

if ~isfile('wormshape.xls')
    writecell({filename},'wormshape.xls');
    temp = readcell('wormshape.xls');temp = size(temp); temp = temp(1);
    range = ['A',num2str(temp+1),':',alphab(2*N),num2str(temp+1)];
    writecell(header,'wormshape.xls','Range',range);
else
    temp = readcell('wormshape.xls');temp = size(temp); temp = temp(1);
    range = ['A',num2str(temp+1)];
    writecell({filename},'wormshape.xls','Range',range);
    temp = readcell('wormshape.xls');temp = size(temp); temp = temp(1);
    range = ['A',num2str(temp+1),':',alphab(2*N),num2str(temp+1)];
    writecell(header,'wormshape.xls','Range',range);
end
temp = readcell('wormshape.xls');temp = size(temp); temp = temp(1);
for i = 1:1:length(x)
    temp2 = length(x{i});
    range = [alphab(i),num2str(temp+1),':',alphab(i),num2str(temp+temp2)];
    writematrix(x{i},'wormshape.xls','Range',range);
```

```
    range = [alphab(i+N),num2str(temp+1),':',alphab(i+N),num2str(temp+temp2)];
    writematrix(y{i},'wormshape.xls','Range',range);
end
```

*Published with MATLAB® R2020a*
