## Supplementary File 2 for "Mitochondrial GTP Metabolism Regulates Reproductive Aging"

This script is made to automatically quantify the fluorescence of GFP labeled mitochondria in live cell imaging. The radial change of fluorescence is quantified in following steps. 1, Manually draw the outlines of cell boundary and cell nucleus to select cytoplasm region 2, Automatically divide the cytoplasm region into concentric polygon 3, Quantify then fluorescence intensity within each polygonal ring. The total, average, and distribution of intensities were calculated. The bioformat package is used to read the image file of factory format. Melissa Linkert, Curtis T. Rueden, Chris Allan, Jean-Marie Burel, Will Moore, Andrew Patterson, Brian Loranger, Josh Moore, Carlos Neves, Donald MacDonald, Aleksandra Tarkowska, Caitlin Sticco, Emma Hill, Mike Rossner, Kevin W. Eliceiri, and Jason R. Swedlow (2010) Metadata matters: access to image data in the real world. The Journal of Cell Biology 189(5), 777-782. doi: 10.1083/jcb.201004104

### Contents


- read file
- image selection
- generation of ring edges
- for display of polygons over image
- ring generation and statistics


### read file

```
addpath 'C:\Users\yitan\Desktop\bfmatlab'
clear;
[file,path]=uigetfile('*.oir','select a oir file');
filename=fullfile(path,file);
datain = bfopen(filename);
datafigure1=datain{1,1};
GFP=datafigure1{1,1};
RFP=datafigure1{2,1};
N = 6; % number of rings + 1
```

### image selection

```
factor = 255/double(max(GFP(:)))*1.5;
[gfp,rectout] = imcrop(adapthisteq(GFP/factor)); % histogram adjustment for display.
rfp = gfp;
figure;
imshow(gfp,[]);
i = 1;
nn = 1;
```

manual nucleus selection

```
while(i<=nn)
    t = title('select nucleus boundary');t.Color = 'red';
    keydoprown = waitforbuttonpress;
    if (keydoprown == 1) % press any key on you keyboard
        [bw1,xi1,yi1]=roipoly;
        i = i+1;
    end
end

% ell boundary selection
i = 1;
```

```
while(i<=nn)
    t = title('select cell boundary');t.Color = 'red';
    keydoprown = waitforbuttonpress;
    if (keydoprown == 1) % press any key on you keyboard
        [bw2,xi2,yi2]=roipoly;
        close all;
        i = i+1;
    end
end
```

### generation of ring edges

The 4 for loops generate radial lines to be intersected with the two polygonal rings.

```
daxiao = size(gfp);
polyin=polyshape(xi1,yi1);
[x,y]=centroid(polyin);
xf=zeros(360,N);
yf = zeros(360,N);
for n = 1:89
    xline=[x x-tand(90-n)*y];
    yline=[y 0];
    [xt1,yt1] = polyxpoly(xline,yline,xi1,yi1);
    [xt2,yt2] = polyxpoly(xline,yline,xi2,yi2);
    xf(n,:)=linspace(xt1,xt2,N);
    yf(n,:)=linspace(yt1,yt2,N);
end
xline6=[x x];
yline6=[y 0];
[x61,y61] = polyxpoly(xline6,yline6,xi1,yi1);
[x62,y62] = polyxpoly(xline6,yline6,xi2,yi2);
xf(90,:)=linspace(x61,x62,N);
yf(90,:) = linspace(y61,y62,N);
for n = 1:89
    xline=[x x+tand(n)*y];
    yline=[y 0];
    [xt1,yt1] = polyxpoly(xline,yline,xi1,yi1);
    [xt2,yt2] = polyxpoly(xline,yline,xi2,yi2);
    xf(n+90,:)=linspace(xt1,xt2,N);
    yf(n+90,:)=linspace(yt1,yt2,N);
end
xline12=[x daxiao(2)];
yline12=[y y];
[x121,y121] = polyxpoly(xline12,yline12,xi1,yi1);
[x122,y122] = polyxpoly(xline12,yline12,xi2,yi2);
xf(180,:)=linspace(x121,x122,N);
yf(180,:)=linspace(y121,y122,N);
for n = 1:89
    xline=[x x+tand(90-n)*(daxiao(1)-y)];
    yline=[y daxiao(1)];
    [xt1,yt1] = polyxpoly(xline,yline,xi1,yi1);
    [xt2,yt2] = polyxpoly(xline,yline,xi2,yi2);
    xf(n+180,:)=linspace(xt1,xt2,N);
    yf(n+180,:)=linspace(yt1,yt2,N);
end
xline18=[x x];
yline18=[y daxiao(1)];
[x181,y181] = polyxpoly(xline18,yline18,xi1,yi1);
[x182,y182] = polyxpoly(xline18,yline18,xi2,yi2);
```

```
xf(270,:)=linspace(x181,x182,N);
yf(270,:)=linspace(y181,y182,N);
for n = 1:89
    xline=[x x-tand(n)*(daxiao(1)-y)];
    yline=[y daxiao(1)];
    [xt1,yt1] = polyxpoly(xline,yline,xi1,yi1);
    [xt2,yt2] = polyxpoly(xline,yline,xi2,yi2);
    xf(n+270,:)=linspace(xt1,xt2,N);
    yf(n+270,:)=linspace(yt1,yt2,N);
end
xline24=[x 0];
yline24=[y y];
[x241,y241] = polyxpoly(xline24,yline24,xi1,yi1);
[x242,y242] = polyxpoly(xline24,yline24,xi2,yi2);
xf(360,:)=linspace(x241,x242,N);
yf(360,:)=linspace(y241,y242,N);
```

### for display of polygons over image

imshow(bw2,[]); hold on; hold off; imshow(bw2,[]); hold on; imshow(rfp,[]); hold on; plot(xi1,yi1,'g'); plot(xf(:,1),yf(:,1),'b'); plot(xf(:,2),yf(:,2),'g'); plot(xf(:,3),yf(:,3),'r'); plot(xf(:,4),yf(:,4),'b'); plot(xf(:,5),yf(:,5),'g'); plot(xf(:,6),yf(:,6),'r'); plot(xf(:,7),yf(:,7),'b'); plot(xf(:,8),yf(:,8),'g'); plot(xf(:,9),yf(:,9),'r'); hold off;

### ring generation and statistics

```
bwcal1 = poly2mask(xf(:,1),yf(:,1),daxiao(1),daxiao(2));
bwcal2 = poly2mask(xf(:,2),yf(:,2),daxiao(1),daxiao(2));
bwcal3 = poly2mask(xf(:,3),yf(:,3),daxiao(1),daxiao(2));
bwcal4 = poly2mask(xf(:,4),yf(:,4),daxiao(1),daxiao(2));
bwcal5 = poly2mask(xf(:,5),yf(:,5),daxiao(1),daxiao(2));
bwcal6 = poly2mask(xf(:,6),yf(:,6),daxiao(1),daxiao(2));
% bwcal7 = poly2mask(xf(:,7),yf(:,7),daxiao(1),daxiao(2));
% bwcal8 = poly2mask(xf(:,8),yf(:,8),daxiao(1),daxiao(2));
% bwcal9 = poly2mask(xf(:,9),yf(:,9),daxiao(1),daxiao(2));
region1=bwcal2&(~bwcal1);
region2=bwcal3&(~bwcal2);
region3=bwcal4&(~bwcal3);
region4=bwcal5&(~bwcal4);
region5=bwcal6&(~bwcal5);
% region6=bwcal7&(~bwcal6);
% region7=bwcal8&(~bwcal7);
% region8=bwcal9&(~bwcal8);
pin1=rfp(region1);
pin2=rfp(region2);
pin3=rfp(region3);
pin4=rfp(region4);
pin5=rfp(region5);
% pin6=rfp(region6);
% pin7=rfp(region7);
% pin8=rfp(region8);
sum5=[sum(double(pin1)),sum(double(pin2)),sum(double(pin3)),sum(double(pin4)),...
    sum(double(pin5))]; % ring standard deviation intensity
sum4 = [mean(pin1),mean(pin2),mean(pin3),mean(pin4),mean(pin5)]; % ring mean intensity
for n= 1: 5
    eval(['max',num2str(n),'=size(pin',num2str(n),');']);
end
sumsize=[max1(1),max2(1),max3(1),max4(1),max5(1)]; % ring size

% add title to each column and create the file if not existing
```

```
head = {'filename',...
    'mean_r1','mean_r2','mean_r3','mean_r4','mean_r5',...
    'size_r1','size_r2','size_r3','size_r4','size_r5',...
    'sum_r1','sum_r2','sum_r3','sum_r4','sum_r5'};
if ~isfile('problem.xls')
    writecell(head,'problem.xls');
end

writecell({file,sum4,sumsize,sum5},'problem.xls','WriteMode','append');
bb=[xi1,yi1];
bb2=[xi2,yi2];
savename=[file 'in.xls'];
writematrix(bb,savename);
savename=[file 'out.xls'];
writematrix(bb2,savename);
```

*Published with MATLAB® R2020a*
