## Supplementary Table 1 for "Mitochondrial GTP Metabolism Regulates Reproductive Aging"

### Supplementary Table 1. RLS summary on HT115 *E. coli*

| Group | Genotype | Condition | Replicates | Mean RLS ± SEM | Number Total (Censors) | p-Value | RLS Extension/ Reduction | Lab Code Strain Number | Figure Number (#Trial No.) |
|---|---|---|---|---|---|---|---|---|---|
| 1 | *WT* | *Ctrl RNAi* | #1 | 3.350 ± 0.150 | 20(0) | | | | Fig 1B (#2) |
| | | | #2 | 3.480 ± 0.201 | 20(1) | | | | |
| | | | #3 | 3.647 ± 0.226 | 20(3) | | | | |
| | | *sucg-1 RNAi* | #1 | 6.300 ± 0.469 | 20(4) | <0.001 | 88.1% | | |
| | | | #2 | 5.707 ± 0.531 | 20(5) | <0.001 | 64% | | |
| | | | #3 | 4.938 ± 0.299 | 20(2) | 0.001 | 35.4% | | |
| 2 | *WT* | *Ctrl RNAi* | #1 | 3.800 ± 0.332 | 20(3) | | | | Fig 1C (#3) |
| | | | #2 | 3.480 ± 0.201 | 20(1) | | | | |
| | | | #3 | 3.647 ± 0.226 | 20(3) | | | | |
| | | *sucl-2 RNAi* | #1 | 6.680 ± 0.241 | 20(4) | <0.001 | 75.8% | | |
| | | | #2 | 6.179 ± 0.456 | 20(8) | <0.001 | 77.6% | | |
| | | | #3 | 6.056 ± 0.239 | 20(2) | <0.001 | 66.1% | | |
| 3 | *WT* | *Ctrl RNAi* | #1 | 3.480 ± 0.201 | 20(1) | | | | Fig 1E (#2) |
| | | | #2 | 3.637 ± 0.140 | 20(2) | | | | |
| | | | #3 | 3.647 ± 0.226 | 20(3) | | | | |
| | | *suca-1 RNAi* | #1 | 3.857 ± 0.287 | 20(3) | 0.277 | N.A. | | |
| | | | #2 | 3.650 ± 0.182 | 20(1) | 0.84 | N.A. | | |
| | | | #3 | 3.925 ± 0.355 | 20(3) | 0.573 | N.A. | | |
| 4 | *rde-1; sun-1p::rde-1* | *Ctrl RNAi* | #1 | 3.100 ± 0.069 | 20(0) | | | DCL569 | Fig 2D (#1) |
| | | | #2 | 3.300 ± 0.179 | 20(0) | | | | |
| | | | #3 | 3.100 ± 0.069 | 20(1) | | | | |
| | | | #4 | 3.400 ± 0.256 | 20(2) | | | | |
| | | *sucg-1 RNAi* | #1 | 5.294 ± 0.445 | 20(4) | <0.001 | 70.8% | | |
| | | | #2 | 4.775 ± 0.316 | 20(2) | <0.001 | 44.7% | | |
| | | | #3 | 4.764 ± 0.336 | 20(1) | <0.001 | 53.7% | | |
| | | | #4 | 4.425 ± 0.248 | 20(1) | <0.001 | 30.1% | | |
| 5 | *rde-1; sun-1p::rde-1* | *Ctrl RNAi* | #1 | 3.100 ± 0.069 | 20(0) | | | DCL569 | Fig 2E (#2) |
| | | | #2 | 3.300 ± 0.179 | 20(0) | | | | |
| | | | #3 | 4.211 ± 0.096 | 20(1) | | | | |
| | | *sucl-2 RNAi* | #1 | 5.700 ± 0.355 | 20(4) | <0.001 | 83.9% | | |
| | | | #2 | 5.373 ± 0.435 | 20(4) | <0.001 | 62.8% | | |
| | | | #3 | 6.300 ± 0.385 | 20(0) | <0.001 | 49.6% | | |
| 6 | *WT* | *EV* | #1 | 4.313 ± 0.277 | 20(4) | | | | Fig S1F (#3) |
| | | | #2 | 4.184 ± 0.125 | 20(5) | | | | |
| | | | #3 | 4.350 ± 0.227 | 20(5) | | | | |
| | | *10mM sodium succinate* | #1 | 3.700 ± 0.219 | 20(1) | 0.133 | N.A. | | |
| | | | #2 | 4.158 ± 0.207 | 20(3) | 0.67 | N.A. | | |
| | | | #3 | 4.182 ± 0.260 | 20(6) | 0.501 | N.A. | | |
| | | *10mM succinic acid* | #1 | 4.500 ± 0.217 | 20(4) | 0.39 | N.A. | | |
| | | | #2 | 4.263 ± 0.104 | 20(4) | 0.683 | N.A. | | |
| | | | #3 | 4.632 ± 0.207 | 20(6) | 0.315 | N.A. | | |
| 7 | *rde-1; sun-1p::rde-1* | *Ctrl RNAi* | #1 | 4.400 ± 0.443 | 20(6) | | | DCL569 | Fig S1G (#2) |
| | | | #2 | 3.158 ± 0.086 | 20(2) | | | | |
| | | | #3 | 3.300 ± 0.105 | 20(1) | | | | |
| | | *mev-1 RNAi* | #1 | 2.105 ± 0.072 | 19(1) | <0.001 | -52.2% | | |
| | | | #2 | 2.278 ± 0.109 | 18(2) | <0.001 | -27.9% | | |
| | | | #3 | 2.150 ± 0.082 | 20(0) | <0.001 | -34.8% | | |
| | | *sdhb-1 RNAi* | #1 | 2.540 ± 0.226 | 20(3) | <0.001 | -42.3% | | |
| | | | #2 | 2.300 ± 0.105 | 20(2) | <0.001 | -27.2% | | |
| | | | #3 | 2.050 ± 0.050 | 20(0) | <0.001 | -37.9% | | |

| | | | | | | | | | |
|---|---|---|---|---|---|---|---|---|---|
| 8 | *Ex[pie-1p:: mito::ndk-1] non-trans siblings* | *Ctrl RNAi* | #1 | 4.563 ± 0.263 | 20(1) | | | MCW1581 | Fig 2G (#1) |
| | | | #2 | 4.535 ± 0.244 | 20(1) | | | | |
| | | | #3 | 4.667 ± 0.360 | 20(2) | | | | |
| | | *sucg-1 RNAi* | #1 | 6.148 ± 0.459 | 20(3) | 0.004 | 34.7% | | |
| | | | #2 | 6.528 ± 0.264 | 20(2) | <0.001 | 43.9% | | |
| | | | #3 | 10.871 ± 0.358 | 19(13) | <0.001 | 132.9% | | |
| | *Ex[pie-1p:: mito::ndk-1] transgenic* | *Ctrl RNAi* | #1 | 4.700 ± 0.291 | 20(0) | | | | |
| | | | #2 | 5.602 ± 0.334 | 20(5) | | | | |
| | | | #3 | 4.800 ± 0.378 | 20(2) | | | | |
| | | *sucg-1 RNAi* | #1 | 4.632 ± 0.370 | 20(3) | 0.957 | N.A. | | |
| | | | #2 | 6.092 ± 0.448 | 19(3) | 0.503 | N.A. | | |
| | | | #3 | 5.600 ± 0.303 | 20(0) | 0.083 | N.A. | | |
| 9 | *rde-1; sun-1p::rde-1* | *Ctrl RNAi* | #1 | 3.300 ± 0.197 | 20(1) | | | DCL569 | Fig 3B (#2) |
| | | | #2 | 3.100 ± 0.069 | 20(1) | | | | |
| | | | #3 | 3.400 ± 0.256 | 20(2) | | | | |
| | | *eat-3 RNAi* | #1 | 4.227 ± 0.222 | 20(1) | 0.001 | 28.1% | | |
| | | | #2 | 4.700 ± 0.272 | 20(0) | <0.001 | 51.6% | | |
| | | | #3 | 4.947 ± 0.408 | 19(2) | <0.001 | 45.5% | | |
| 10 | *rde-1; sun-1p::rde-1* | *Ctrl RNAi* | #1 | 3.100 ± 0.069 | 20(1) | | | DCL569 | Fig 3D (#3) |
| | | | #2 | 3.400 ± 0.256 | 20(2) | | | | |
| | | | #3 | 4.211 ± 0.096 | 20(1) | | | | |
| | | *fzo-1 RNAi* | #1 | 3.457 ± 0.140 | 20(1) | 0.028 | 11.5% | | |
| | | | #2 | 2.947 ± 0.120 | 19(1) | 0.222 | N.A. | | |
| | | | #3 | 4.500 ± 0.170 | 20(0) | 0.127 | N.A. | | |
| 11 | *rde-1; sun-1p::rde-1* | *Ctrl RNAi* | #1 | 3.300 ± 0.197 | 20(1) | | | DCL569 | Fig 3E (#1) |
| | | | #2 | 3.100 ± 0.069 | 20(1) | | | | |
| | | | #3 | 3.400 ± 0.256 | 20(2) | | | | |
| | | *drp-1 RNAi* | #1 | 3.150 ± 0.218 | 20(3) | 0.567 | N.A. | | |
| | | | #2 | 2.963 ± 0.339 | 20(2) | 0.1 | N.A. | | |
| | | | #3 | 2.619 ± 0.176 | 20(2) | 0.016 | -23.0% | | |
| 12 | *WT* | | #1 | 4.345 ± 0.196 | 20(3) | | | | Fig 3F (#3) |
| | | | #2 | 3.285 ± 0.133 | 20(4) | | | | |
| | | | #3 | 3.550 ± 0.211 | 20(0) | | | | |
| | *pie-1p:: drp-1 TG* | | #1 | 9.875 ± 0.165 | 20(17) | <0.001 | 127.3% | MCW1220 | |
| | | | #2 | 7.635 ± 0.417 | 20(7) | <0.001 | 132.4% | | |
| | | | #3 | 9.833 ± 0.345 | 20(8) | <0.001 | 177.0% | | |
| 13 | *gfp::degron::drp-1; sun-1p:: TIR1::mRuby* | *- Auxin* | #1 | 4.474 ± 0.234 | 19(0) | | | MCW1326 | Fig 4C (#2) |
| | | | #2 | 4.579 ± 0.268 | 20(1) | | | | |
| | | | #3 | 4.878 ± 0.511 | 20(3) | | | | |
| | | *+ Auxin* | #1 | 4.610 ± 0.279 | 20(1) | 0.666 | N.A. | | |
| | | | #2 | 4.245 ± 0.163 | 20(3) | 0.322 | N.A. | | |
| | | | #3 | 4.088 ± 0.214 | 19(2) | 0.331 | N.A. | | |
| 14 | *gfp::degron::drp-1; sun-1p:: TIR1::mRuby* | *- Auxin & Ctrl RNAi* | #1 | 3.944 ± 0.298 | 20(1) | | | MCW1326 | Fig 4D (#3) |
| | | | #2 | 5.267 ± 0.475 | 20(3) | | | | |
| | | | #3 | 4.170 ± 0.217 | 20(1) | | | | |
| | | *- Auxin & sucg-1 RNAi* | #1 | 5.100 ± 0.278 | 20(5) | 0.011 | 29.3% | | |
| | | | #2 | 6.467 ± 0.366 | 20(4) | 0.127 | N.A. | | |
| | | | #3 | 6.102 ± 0.466 | 20(4) | <0.001 | 46.3% | | |
| | | *+ Auxin & Ctrl RNAi* | #1 | 3.917 ± 0.222 | 18(1) | | | | |
| | | | #2 | 4.077 ± 0.185 | 19(2) | | | | |
| | | | #3 | 4.250 ± 0.310 | 19(4) | | | | |
| | | *+ Auxin & sucg-1 RNAi* | #1 | 4.235 ± 0.369 | 18(3) | 0.468 | N.A. | | |
| | | | #2 | 3.684 ± 0.134 | 19(0) | 0.069 | N.A. | | |
| | | | #3 | 3.815 ± 0.187 | 18(2) | 0.248 | N.A. | | |
| 15 | *WT* | | #1 | 4.412 ± 0.193 | 18(2) | | | | Fig S5B |
| | | | #2 | 4.684 ± 0.253 | 20(4) | | | | |

| | | | | | | | | | |
|---|---|---|---|---|---|---|---|---|---|
| | | | #3 | 3.285 ± 0.133 | 20(4) | | | | (#1), S5C (#2) |
| | *suca-1 (rax84)* | | #1 | 4.278 ± 0.253 | 18(0) | 0.608 | N.A. | MCW1329 | |
| | | | #2 | 4.450 ± 0.267 | 20(1) | 0.637 | N.A. | | |
| | | | #3 | 3.474 ± 0.140 | 20(1) | 0.376 | N.A. | | |
| | *suca-1 (rax85)* | | #1 | 4.765 ± 0.304 | 17(0) | 0.512 | N.A. | MCW1330 | |
| | | | #2 | 4.625 ± 0.305 | 20(2) | 0.915 | N.A. | | |
| | | | #3 | 3.420 ± 0.147 | 20(2) | 0.573 | N.A. | | |
| 16 | *Paternal line: sucg-1::gfp/ sucg-1 (rax86)* | *F1: gfp/gfp* | #1 | 3.813 ± 0.179 | 20(1) | | | PHX3617 | Fig 5B (#2) |
| | | | #2 | 3.889 ± 0.219 | 24(2) | | | | |
| | | | #3 | 3.719 ± 0.167 | 17(3) | | | | |
| | | *F1: gfp/KO* | #1 | 4.396 ± 0.258 | 39(6) | 0.132 | N.A. | MCW1385 | |
| | | | #2 | 4.150 ± 0.177 | 36(5) | 0.357 | N.A. | | |
| | | | #3 | 4.114 ± 0.107 | 38(3) | 0.15 | N.A. | | |
| | | *F1: KO/KO* | #1 | 6.182 ± 0.197 | 21(11) | <0.001 *(gfp/gfp)* <0.001 *(gfp/KO)* | 62.1% *(gfp/gfp)* 40.6% *(gfp/KO)* | MCW1331 | |
| | | | #2 | 7.015 ± 0.256 | 20(9) | <0.001 *(gfp/gfp)* <0.001 *(gfp/KO)* | 80.4% *(gfp/gfp)* 69.0% *(gfp/KO)* | | |
| | | | #3 | 6.832 ± 0.324 | 22(7) | <0.001 *(gfp/gfp)* <0.001 *(gfp/KO)* | 83.7% *(gfp/gfp)* 66.1% *(gfp/KO)* | | |
| 17 | *Paternal line: sucg-1::gfp/ sucg-1 (rax83)* | *F1: gfp/gfp* | #1 | 4.469 ± 0.252 | 24(4) | | | PHX3617 | Fig S5D (#1) |
| | | | #2 | 4.125 ± 0.301 | 16(0) | | | | |
| | | | #3 | 3.618 ± 0.205 | 17(2) | | | | |
| | | *F1: gfp/KO* | #1 | 4.455 ± 0.180 | 39(4) | 0.779 | N.A. | MCW1375 | |
| | | | #2 | 4.116 ± 0.150 | 43(0) | 0.939 | N.A. | | |
| | | | #3 | 3.200 ± 0.057 | 50(5) | 0.088 | N.A. | | |
| | | *F1: KO/KO* | #1 | 7.103 ± 0.365 | 15(5) | <0.001 *(gfp/gfp)* <0.001 *(gfp/KO)* | 59.0% *(gfp/gfp)* 59.4% *(gfp/KO)* | MCW1325 | |
| | | | #2 | 6.667 ± 0.457 | 21(3) | <0.001 *(gfp/gfp)* <0.001 *(gfp/KO)* | 61.6% *(gfp/gfp)* 62.0% *(gfp/KO)* | | |
| | | | #3 | 8.077 ± 0.431 | 13(0) | <0.001 *(gfp/gfp)* <0.001 *(gfp/KO)* | 123.2% *(gfp/gfp)* 152.4% *(gfp/KO)* | | |
| 18 | *rde-1; sun-1p::rde-1* | *Ctrl RNAi* | #1 | 3.100 ± 0.069 | 20(0) | | | DCL569 | Fig S6C (#3) |
| | | | #2 | 3.300 ± 0.179 | 20(0) | | | | |
| | | | #3 | 3.400 ± 0.256 | 20(2) | | | | |
| | | *suca-1 RNAi* | #1 | 3.050 ± 0.050 | 20(0) | 0.553 | N.A. | | |
| | | | #2 | 3.479 ± 0.197 | 20(3) | 0.408 | N.A. | | |
| | | | #3 | 3.211 ± 0.123 | 19(3) | 0.785 | N.A. | | |
| 19 | *WT* | *EV* | #1 | 3.925 ± 0.256 | 20(1) | | | | Fig S7A (#1) |
| | | | #2 | 3.889 ± 0.285 | 20(3) | | | | |
| | | | #3 | 3.733 ± 0.246 | 20(2) | | | | |
| | | *128nM meCbl* | #1 | 4.125 ± 0.365 | 20(4) | 0.675 | N.A. | | |
| | | | #2 | 3.350 ± 0.131 | 20(0) | 0.096 | N.A. | | |
| | | | #3 | 3.481 ± 0.145 | 20(2) | 0.388 | N.A. | | |
| 20 | *WT* | *EV* | #1 | 3.650 ± 0.150 | 20(1) | | | | Fig S7B (#2) |
| | | | #2 | 3.600 ± 0.169 | 20(0) | | | | |

| | | | | | | | | | |
|---|---|---|---|---|---|---|---|---|---|
| | | | #3 | 4.088 ± 0.249 | 20(2) | | | | |
| | | *128nM adoCbl* | #1 | 3.611 ± 0.158 | 20(2) | 0.871 | N.A. | | |
| | | | #2 | 3.379 ± 0.144 | 20(2) | 0.264 | N.A. | | |
| | | | #3 | 3.526 ± 0.234 | 20(1) | 0.104 | N.A. | | |
| 21 | *WT* | *Ctrl RNAi* | #1 | 3.875 ± 0.235 | 20(2) | | | | Fig S7E (#2) |
| | | | #2 | 3.632 ± 0.256 | 20(2) | | | | |
| | | | #3 | 3.867 ± 0.318 | 20(2) | | | | |
| | | *metr-1 RNAi* | #1 | 4.638 ± 0.197 | 20(2) | 0.029 | 19.7% | | |
| | | | #2 | 5.343 ± 0.320 | 20(4) | 0.001 | 47.1% | | |
| | | | #3 | 5.368 ± 0.297 | 20(4) | 0.001 | 38.8% | | |
| | | *mmcm-1 RNAi* | #1 | 3.770 ± 0.199 | 20(1) | 0.732 | N.A. | | |
| | | | #2 | 3.643 ± 0.207 | 20(2) | 0.902 | N.A. | | |
| | | | #3 | 3.211 ± 0.123 | 20(1) | 0.057 | N.A. | | |
