## Supplementary Table 2 for "Mitochondrial GTP Metabolism Regulates Reproductive Aging"

### Supplementary Table 2. Lifespan summary

| Group | Genotype | Condition | Replicates | Mean LS ± SEM | Number Total (Censors) | p-Value | LS Extension/ Reduction | Lab Code Strain Number | Figure Number (#Trial No.) |
|---|---|---|---|---|---|---|---|---|---|
| 1 | *WT* | *Ctrl RNAi* | #1 | 20.276 ± 0.478 | 89 (23) | | | | Fig S1D (#2) |
| | | | #2 | 19.978 ± 0.396 | 100 (19) | | | | |
| | | | #3 | 20.664 ± 0.567 | 100 (12) | | | | |
| | | *sucg-1 RNAi* | #1 | 22.363 ± 0.792 | 95 (22) | 0.002 | 10.3% | | |
| | | | #2 | 23.294 ± 0.776 | 96 (28) | <0.001 | 16.6% | | |
| | | | #3 | 24.327 ± 0.867 | 100 (36) | <0.001 | 17.7% | | |
| | | *sucl-2 RNAi* | #1 | 24.150 ± 0.683 | 103 (41) | <0.001 | 19.1% | | |
| | | | #2 | 23.014 ± 0.543 | 100 (24) | 0.037 | 15.2% | | |
| | | | #3 | 22.337 ± 0.867 | 90 (31) | <0.001 | 8.1% | | |
| | | *suca-1 RNAi* | #1 | 20.316 ± 0.457 | 92 (25) | 0.801 | N.A. | | |
| | | | #2 | 19.745 ± 0.395 | 90 (22) | 0.472 | N.A. | | |
| | | | #3 | 21.560 ± 0.814 | 100 (24) | 0.093 | N.A. | | |
| 2 | *rde-1; sun-1p::rde-1* | *Ctrl RNAi* | #1 | 17.942 ± 0.408 | 95 (6) | | | DCL569 | Fig S1E (#2) |
| | | | #2 | 17.779 ± 0.510 | 95 (20) | | | | |
| | | | #3 | 18.760 ± 0.353 | 130 (21) | | | | |
| | | *sucg-1 RNAi* | #1 | 20.328 ± 0.670 | 90 (15) | <0.001 | 13.3% | | |
| | | | #2 | 20.728 ± 0.667 | 90 (19) | <0.001 | 16.6% | | |
| | | | #3 | 18.060 ± 0.416 | 126 (15) | 0.784 | N.A. | | |
| | | *sucl-2 RNAi* | #1 | 20.014 ± 0.592 | 100 (37) | 0.002 | 11.5% | | |
| | | | #2 | 20.600 ± 0.583 | 95 (31) | 0.001 | 15.9% | | |
| | | | #3 | 20.194 ± 0.357 | 119 (21) | 0.006 | 7.6% | | |
| | | *suca-1 RNAi* | #1 | 17.631 ± 0.338 | 95 (9) | 0.318 | N.A. | | |
| | | | #2 | 17.543 ± 0.418 | 92 (23) | 0.439 | N.A. | | |
| | | | #3 | 17.010 ± 0.375 | 107 (8) | 0.011 | -9.3% | | |
