## Supplementary Table 3 for "Mitochondrial GTP Metabolism Regulates Reproductive Aging"

### Supplementary Table 3. RLS summary on OP50 *E. coli*

| Group | Genotype | Condition | Replicates | Mean RLS ± SEM | Number Total (Censors) | p-Value | RLS Extension/ Reduction | Lab Code Strain Number | Figure Number (#Trial No.) |
|---|---|---|---|---|---|---|---|---|---|
| 1 | *Paternal line: sucg-1:: gfp/ sucg-1 (rax86)* | *F1: gfp/gfp* | #1 | 6.458 ± 0.638 | 17(5) | | | PHX3617 | Fig 5C (#2) |
| | | | #2 | 6.471 ± 0.365 | 17(0) | | | | |
| | | | #3 | 7.935 ± 0.638 | 19(7) | | | | |
| | | *F1: gfp/KO* | #1 | 6.296 ± 0.298 | 43(13) | 0.839 | N.A. | MCW1385 | |
| | | | #2 | 6.327 ± 0.308 | 39(5) | 0.989 | N.A. | | |
| | | | #3 | 7.118 ± 0.489 | 34(13) | 0.414 | N.A. | | |
| | | *F1: KO/KO* | #1 | 6.953 ± 0.569 | 16(5) | 0.705 *(gfp/gfp)*<br>0.363 *(gfp/KO)* | N.A. *(gfp/gfp)*<br>N.A. *(gfp/KO)* | MCW1331 | |
| | | | #2 | 6.354 ± 0.250 | 21(4) | 0.853 *(gfp/gfp)*<br>0.742 *(gfp/KO)* | N.A. *(gfp/gfp)*<br>N.A. *(gfp/KO)* | | |
| | | | #3 | 6.765 ± 0.345 | 24(5) | 0.574 *(gfp/gfp)*<br>0.074 *(gfp/KO)* | N.A. *(gfp/gfp)*<br>N.A. *(gfp/KO)* | | |
| 2 | *Paternal line: sucg-1:: gfp/ sucg-1 (rax83)* | *F1: gfp/gfp* | #1 | 6.985 ± 0.507 | 21(6) | | | PHX3617 | Fig S5E (#1) |
| | | | #2 | 5.526 ± 0.309 | 19(0) | | | | |
| | | | #3 | 6.617 ± 0.569 | 17(4) | | | | |
| | | *F1: gfp/KO* | #1 | 6.596 ± 0.255 | 48(7) | 0.472 | N.A. | MCW1375 | |
| | | | #2 | 5.976 ± 0.294 | 42(0) | 0.243 | N.A. | | |
| | | | #3 | 7.667 ± 0.231 | 54(5) | 0.09 | N.A. | | |
| | | *F1: KO/KO* | #1 | 5.933 ± 0.250 | 10(1) | 0.155 *(gfp/gfp)*<br>0.212 *(gfp/KO)* | N.A. *(gfp/gfp)*<br>N.A. *(gfp/KO)* | MCW1325 | |
| | | | #2 | 6.842 ± 0.479 | 19(0) | 0.028 *(gfp/gfp)*<br>0.144 *(gfp/KO)* | N.A. *(gfp/gfp)*<br>N.A. *(gfp/KO)* | | |
| | | | #3 | 7.741 ± 0.899 | 9(2) | 0.215 *(gfp/gfp)*<br>0.424 *(gfp/KO)* | N.A. *(gfp/gfp)*<br>N.A. *(gfp/KO)* | | |
| 3 | *WT* | | #1 | 7.029 ± 0.511 | 17(5) | | | | Fig S5F (#1), S5G (#2) |
| | | | #2 | 7.352 ± 0.553 | 20(7) | | | | |
| | | | #3 | 6.200 ± 0.392 | 20(6) | | | | |
| | *suca-1 (rax84)* | | #1 | 6.063 ± 0.471 | 18(3) | 0.209 | N.A. | MCW1329 | |
| | | | #2 | 7.273 ± 0.463 | 20(4) | 0.796 | N.A. | | |
| | | | #3 | 6.019 ± 0.262 | 20(4) | 0.746 | N.A. | | |
| | *suca-1 (rax85)* | | #1 | 6.206 ± 0.387 | 16(5) | 0.343 | N.A. | MCW1330 | |
| | | | #2 | 7.052 ± 0.571 | 20(4) | 0.501 | N.A. | | |
| | | | #3 | 5.315 ± 0.255 | 20(4) | 0.091 | N.A. | | |
| 4 | *rde-1; sun-1p::rde-1* | *Ctrl RNAi* | #1 | 4.768 ± 0.196 | 20(1) | | | DCL569 | Fig S6D (#4) |
| | | | #2 | 5.398 ± 0.455 | 20(4) | | | | |
| | | | #3 | 4.900 ± 0.299 | 20(1) | | | | |
| | | | #4 | 4.613 ± 0.291 | 20(2) | | | | |
| | | | #1 | 5.613 ± 0.404 | 20(6) | 0.07 | N.A. | | |

| | | | | | | | | | |
|---|---|---|---|---|---|---|---|---|---|
| | | *sucg-1 RNAi* | #2 | 5.600 ± 0.462 | 20(2) | 0.64 | N.A. | | |
| | | | #3 | 5.280 ± 0.386 | 20(3) | 0.602 | N.A. | | |
| | | | #4 | 4.941 ± 0.210 | 20(3) | 0.426 | N.A. | | |
| 5 | *rde-1; sun-1p::rde-1* | *Ctrl RNAi* | #1 | 4.768 ± 0.196 | 20(1) | | | DCL569 | Fig S6E (#3) |
| | | | #2 | 5.398 ± 0.455 | 20(4) | | | | |
| | | | #3 | 4.900 ± 0.299 | 20(1) | | | | |
| | | | #4 | 4.613 ± 0.291 | 20(2) | | | | |
| | | *suca-1 RNAi* | #1 | 4.425 ± 0.175 | 20(1) | 0.19 | N.A. | | |
| | | | #2 | 5.181 ± 0.457 | 20(3) | 0.879 | N.A. | | |
| | | | #3 | 4.250 ± 0.216 | 20(1) | 0.122 | N.A. | | |
| | | | #4 | 4.529 ± 0.236 | 20(1) | 0.708 | N.A. | | |
| 6 | *rde-1; sun-1p::rde-1* | *Ctrl RNAi* | #1 | 4.768 ± 0.196 | 20(1) | | | DCL569 | Fig S6F (#2) |
| | | | #2 | 5.398 ± 0.455 | 20(4) | | | | |
| | | | #3 | 6.556 ± 0.336 | 20(2) | | | | |
| | | *sucl-2 RNAi* | #1 | 5.553 ± 0.260 | 20(2) | 0.024 | 16.5% | | |
| | | | #2 | 4.888 ± 0.227 | 19(2) | 0.441 | N.A. | | |
| | | | #3 | 6.167 ± 0.336 | 20(2) | 0.254 | N.A. | | |
| 5 | *rde-1; sun-1p::rde-1* | *Ctrl RNAi* | #1 | 4.847 ± 0.277 | 20(1) | | | DCL569 | Fig S6G (#1) |
| | | | #2 | 4.900 ± 0.299 | 20(1) | | | | |
| | | | #3 | 4.613 ± 0.291 | 20(2) | | | | |
| | | *drp-1 RNAi* | #1 | 3.179 ± 0.267 | 20(2) | <0.001 | -34.3% | | |
| | | | #2 | 2.825 ± 0.201 | 20(1) | <0.001 | -42.3% | | |
| | | | #3 | 2.684 ± 0.172 | 19(1) | <0.001 | -41.8% | | |
| 6 | *gfp::degron::drp-1; sun-1p::TIR1::mRuby* | *- Auxin* | #1 | 7.121 ± 0.486 | 20(4) | | | MCW1326 | Fig 6B (#2), 6C (#1) |
| | | | #2 | 7.063 ± 0.426 | 19(4) | | | | |
| | | | #3 | 7.506 ± 0.556 | 19(5) | | | | |
| | | *+ Auxin* | #1 | 4.944 ± 0.276 | 20(1) | <0.001 | -30.6% | | |
| | | | #2 | 4.933 ± 0.341 | 20(3) | 0.002 | -30.2% | | |
| | | | #3 | 4.519 ± 0.293 | 20(4) | <0.001 | -39.8% | | |
| | | *+ Auxin Post-L4* | #1 | 4.874 ± 0.307 | 19(1) | <0.001 | -31.6% | | |
| | | | #2 | 5.033 ± 0.330 | 20(5) | 0.003 | -28.8% | | |
| | | | #3 | 4.019 ± 0.209 | 18(1) | <0.001 | -46.5% | | |
| 7 | *WT* | | #1 | 6.066 ± 0.326 | 20(3) | | | | Fig 6D (#3) |
| | | | #2 | 6.200 ± 0.392 | 20(6) | | | | |
| | | | #3 | 7.335 ± 0.467 | 20(1) | | | | |
| | *pie-1p::drp-1 TG* | | #1 | 10.800 ± 0.179 | 20(18) | <0.001 | 78.0% | MCW1220 | |
| | | | #2 | 10.282 ± 0.281 | 20(15) | <0.001 | 65.8% | | |
| | | | #3 | 9.412 ± 0.412 | 20(3) | 0.003 | 28.3% | | |
| 8 | *rde-1; sun-1p::rde-1* | *Ctrl RNAi* | #1 | 4.847 ± 0.277 | 20(1) | | | DCL569 | Fig 6E (#3) |
| | | | #2 | 4.900 ± 0.299 | 20(1) | | | | |
| | | | #3 | 4.613 ± 0.291 | 20(2) | | | | |
| | | | #4 | 6.556 ± 0.336 | 20(2) | | | | |
| | | *eat-3 RNAi* | #1 | 5.319 ± 0.286 | 20(2) | 0.282 | N.A. | | |
| | | | #2 | 5.000 ± 0.171 | 19(0) | 0.774 | N.A. | | |
| | | | #3 | 5.368 ± 0.311 | 20(6) | 0.099 | N.A. | | |
| | | | #4 | 5.474 ± 0.234 | 20(1) | 0.005 | -16.5% | | |
| 9 | *rde-1; sun-1p::rde-1* | *Ctrl RNAi* | #1 | 4.900 ± 0.299 | 20(1) | | | DCL569 | Fig S6H (#3) |
| | | | #2 | 4.613 ± 0.291 | 20(2) | | | | |
| | | | #3 | 6.556 ± 0.336 | 20(2) | | | | |
| | | *fzo-1 RNAi* | #1 | 5.125 ± 0.292 | 20(1) | 0.62 | N.A. | | |
| | | | #2 | 5.088 ± 0.370 | 20(3) | 0.296 | N.A. | | |
| | | | #3 | 6.158 ± 0.257 | 20(1) | 0.178 | N.A. | | |
| 10 | *WT* | *EV* | #1 | 7.250 ± 0.380 | 20(6) | | | | Fig 7A (#1) |
| | | | #2 | 8.018 ± 0.507 | 19(7) | | | | |
| | | | #3 | 7.158 ± 0.507 | 20(4) | | | | |
| | | | #1 | 3.825 ± 0.224 | 20(2) | <0.001 | -47.2% | | |

| | | | | | | | | | |
|---|---|---|---|---|---|---|---|---|---|
| | | *128nM meCbl* | #2 | 3.526 ± 0.160 | 20(1) | <0.001 | -56% | | |
| | | | #3 | 3.368 ± 0.137 | 20(1) | <0.001 | -52.9% | | |
| 11 | *WT* | *EV* | #1 | 6.444 ± 0.349 | 20(5) | | | | Fig 7B (#2) |
| | | | #2 | 6.904 ± 0.523 | 20(5) | | | | |
| | | | #3 | 6.667 ± 0.444 | 20(2) | | | | |
| | | *128nM adoCbl* | #1 | 3.964 ± 0.329 | 20(3) | <0.001 | -38.5% | | |
| | | | #2 | 3.719 ± 0.278 | 20(2) | <0.001 | -46.1% | | |
| | | | #3 | 4.053 ± 0.450 | 19(5) | <0.001 | -39.2% | | |
| 12 | *Paternal line: sucg-1:: gfp/ sucg-1 (rax83)* | *F1: gfp/gfp EV* | #1 | 7.482 ± 0.553 | 16(7) | | | PHX3617 | Fig 7D (#3), S7C (#3), S7D (#3) |
| | | | #2 | 6.928 ± 0.358 | 23(8) | | | | |
| | | | #3 | 6.147 ± 0.436 | 20(4) | | | | |
| | | *F1: gfp/KO EV* | #1 | 6.947 ± 0.332 | 38(13) | 0.293 | N.A. | MCW1375 | |
| | | | #2 | 6.644 ± 0.326 | 38(11) | 0.651 | N.A. | | |
| | | | #3 | 6.064 ± 0.194 | 44(6) | 0.805 | N.A. | | |
| | | *F1: KO/KO EV* | #1 | 7.061 ± 0.293 | 25(7) | 0.453 *(gfp/gfp)*<br>0.881 *(gfp/KO)* | N.A. (gfp/gfp)<br>N.A. (gfp/KO) | MCW1325 | |
| | | | #2 | 6.451 ± 0.260 | 14(4) | 0.286 *(gfp/gfp)*<br>0.579 *(gfp/KO)* | N.A. (gfp/gfp)<br>N.A. (gfp/KO) | | |
| | | | #3 | 6.408 ± 0.445 | 16(4) | 0.745 *(gfp/gfp)*<br>0.531 *(gfp/KO)* | N.A. (gfp/gfp)<br>N.A. (gfp/KO) | | |
| | | *F1: gfp/gfp 128nM meCbl* | #1 | 4.889 ± 0.414 | 21(8) | | | PHX3617 | |
| | | | #2 | 3.734 ± 0.219 | 23(3) | | | | |
| | | | #3 | 3.467 ± 0.157 | 24(3) | | | | |
| | | *F1: gfp/KO 128nM meCbl* | #1 | 4.661 ± 0.326 | 34(9) | 0.62 | N.A. | MCW1375 | |
| | | | #2 | 3.686 ± 0.209 | 42(4) | 0.654 | N.A. | | |
| | | | #3 | 3.504 ± 0.122 | 44(10) | 0.682 | N.A. | | |
| | | *F1: KO/KO 128nM meCbl* | #1 | 6.772 ± 0.384 | 25(11) | 0.004 *(gfp/gfp)*<br>0.001 *(gfp/KO)* | 38.5% (gfp/gfp)<br>45.3% (gfp/KO) | MCW1325 | |
| | | | #2 | 6.528 ± 0.339 | 15(7) | <0.001 *(gfp/gfp)*<br><0.001 *(gfp/KO)* | 74.8% (gfp/gfp)<br>77.1% (gfp/KO) | | |
| | | | #3 | 6.331 ± 0.520 | 12(4) | <0.001 *(gfp/gfp)*<br><0.001 *(gfp/KO)* | 82.6% (gfp/gfp)<br>80.7% (gfp/KO) | | |
| 13 | *WT* | *Ctrl RNAi* | #1 | 6.534 ± 0.330 | 20(2) | | | | Fig S7F (#2) |
| | | | #2 | 5.933 ± 0.291 | 20(3) | | | | |
| | | | #3 | 6.899 ± 0.618 | 20(6) | | | | |
| | | *metr-1 RNAi* | #1 | 7.043 ± 0.386 | 20(7) | 0.184 | N.A. | | |
| | | | #2 | 5.877 ± 0.412 | 20(3) | 0.79 | N.A. | | |
| | | | #3 | 5.952 ± 0.614 | 20(5) | 0.394 | N.A. | | |
| | | *mmcm-1 RNAi* | #1 | 6.463 ± 0.372 | 20(6) | 0.778 | N.A. | | |
| | | | #2 | 6.034 ± 0.357 | 20(5) | 0.751 | N.A. | | |
| | | | #3 | 6.453 ± 0.382 | 20(4) | 0.525 | N.A. | | |
