## Supplementary Table 4 for "Mitochondrial GTP Metabolism Regulates Reproductive Aging"

### Supplementary Table 4. RLS summary on various dietary conditions

| Group | Genotype | Condition | Replicates | Mean RLS ± SEM | Number Total (Censors) | p-Value | RLS Extension/ Reduction | Lab Code Strain Number | Figure Number (#Trial No.) |
|---|---|---|---|---|---|---|---|---|---|
| 1 | *WT* | *HT115 bacteria* | #1 | 4.110 ± 0.196 | 20(1) | | | | Fig S6A (#1) |
| | | | #2 | 3.875 ± 0.191 | 20(1) | | | | |
| | | | #3 | 4.345 ± 0.196 | 20(3) | | | | |
| | | *HB101 bacteria* | #1 | 3.775 ± 0.251 | 20(1) | 0.43 | N.A. | | |
| | | | #2 | 3.526 ± 0.193 | 20(2) | 0.133 | N.A. | | |
| | | | #3 | 4.000 ± 0.272 | 20(1) | 0.353 | N.A. | | |
| | | *OP50 bacteria* | #1 | 7.069 ± 0.364 | 20(3) | <0.001 (HT115) <0.001 (HB101) | 72.0% (HT115) 87.3% (HB101) | | |
| | | | #2 | 7.044 ± 0.554 | 20(6) | <0.001 (HT115) <0.001 (HB101) | 81.8% (HT115) 99.8% (HB101) | | |
| | | | #3 | 6.066 ± 0.326 | 20(3) | 0.001 (HT115) 0.001 (HB101) | 39.6% (HT115) 51.7% (HB101) | | |
