## Supplementary Table 5 for "Mitochondrial GTP Metabolism Regulates Reproductive Aging"

### Supplementary Table 5. Primers

| Primer name | Primer sequence | Source | Experimental assay |
|---|---|---|---|
| *sucg-1_gfp_fwd* | GTGGAGCAGTCACAGAGGAT | This study | Genotyping |
| *sucg-1_gfp_rev* | CAGTGTACGAAGGATCAGTG | This study | Genotyping |
| *suca-1_gfp_fwd* | AGATGGGATTCTCGGTACGT | This study | Genotyping |
| *suca-1_gfp_rev* | CCCAGAGAAGAAAACAACCA | This study | Genotyping |
| *unc-119_fwd* | AAATTGCATGCCAGCACCGGTC | *Maduro and Pilgrim, 1995* | Genotyping |
| *unc-119_rev* | AGTCGGCCTTATTGTGCATTAC | *Maduro and Pilgrim, 1995* | Genotyping |
| *pcrYT1_pie-1p_fwd* | ATAAAATTGAGCTTTGTTTGTAAATAATC | This study | Cloning |
| *pcrYT1_pie-1p_rev* | TGATCATTTTTGCTAGCCAAGG | This study | Cloning |
| *pcrYT1_mito_ndk-1_fwd* | TAGAGGATCCCCGGGGACCCTTGGCTAGCA AAAATGATCAATGTCCGTCCTTACCCCACT | This study | Cloning |
| *pcrYT1_mito_ndk-1_rev* | GATAAATATACAAAACAAAATATGGAAAATAC GGCAAAATTACGCATAATCTGGAACATC | This study | Cloning |
| *pcrYT1_pie-1_3'UTR_fwd* | TTTTGCCGTATTTTCCATATTTTGTTTTG | This study | Cloning |
| *pcrYT1_pie-1_3'UTR_rev* | ATCATCGTTCACTTTTCACGG | This study | Cloning |
| *attB1_drp-1_fwd* | GGGGACAAGTTTGTACAAAAAAGCAGGCTC AAAAATGGAAAATCTCATTCCTGTCG | This study | Cloning |
| *attB2_drp-1_rev* | GGGGACCACTTTGTACAAGAAAGCTGGGTG GAATATCACCAAACTTGTGTTTCTCTC | This study | Cloning |
| *GFP_Degron_drp-1_30bp_HA_fwd* | TGGCATGGATGAACTATACAAACACTCCACC TCAGGTTCTCCTAAAGATCCAGCCAAACC | This study | PCR Amplification (Repair Template for CRISPR-Cas9 Mediated Insertion) |
| *GFP_Degron_drp-1_30bp_HA_rev* | TGACGACAGGAATGAGATTTTCTCTAGAGGT TCCAGAGCCCTTCACGAACGCCGCCGCCT | This study | PCR Amplification (Repair Template for CRISPR-Cas9 Mediated Insertion) |
| *GFP_Degron_drp-1_100bp_HA_fwd* | CGAAAGATCCCAACGAAAAGAGAGACCACA TGGTCCTTCTTGAGTTTGTAACAGCTGCTGG GATTACACATGGCATGGATGAACTATACAAA CACTCCACCTCAGGTTCTCCTAAAGATCCAG CCAAACCTCCGG | This study | PCR Amplification (Repair Template for CRISPR-Cas9 Mediated Insertion) |
| *GFP_Degron_drp-1_100bp_HA_rev* | AATCTGTGGAAGTTGAATTTGATCTTCTTTCC TGCCTAACGTTGCGAAAACATCCTGTAGTTT ATTGACGACAGGAATGAGATTTTCTCTAGAG GTTCCAGAGCCCTTCACGAACGCCGCCGCC TCCGGGCCACC | This study | PCR Amplification (Repair Template for CRISPR-Cas9 Mediated Insertion) |
| *GFP_drp-1_fwd* | AGACTTCGATGCCGTGCGAA | This study | Genotyping |
| *GFP_drp-1_rev* | ACCTGTTCTCCATGAGCACT | This study | Genotyping |
| *drp-1_tm1108_fwd* | TTCCCGGCATCACAAAGATC | This study | Genotyping |
| *drp-1_tm1108_rev* | TCCTGTCGCGTTTCTAATCG | This study | Genotyping |
| *sucg-1_KO_fwd* | TAGGCGTTTAGAACATGGTTC | This study | Genotyping |
| *sucg-1_KO_rev* | CAGGAAAACACAACGGTAAAG | This study | Genotyping |
| *suca-1_KO_fwd* | TCCAACTGATCATGCATTTGAAA | This study | Genotyping |
| *suca-1_KO_rev* | AACCGGATTCCTCTCTTCAC | This study | Genotyping |
| *pYT17_backbone_fwd* | TCGTCTCGCGCGTTTCGG | This study | Cloning |
| *pYT17_backbone_rev* | CATGGTCATAGCTGTTTCCT | This study | Cloning |
| *pYT17_sun-1p_fwd* | AGGAAACAGCTATGACCATGTCACAGCAGA GAAGGCAAAC | This study | Cloning |
| *pYT17_sun-1p_rev* | AGCTCCTCTCCCTTGGACATACCGAGTAGAT CTGGAAGTTTAG | This study | Cloning |
| *pYT17_modified_eGFP_fwd* | ATGTCCAAGGGAGAGGAGCTCTTCA | This study | Cloning |

| | | | |
|---|---|---|---|
| *pYT17_modified_eGFP_rev* | CTTGTAGAGCTCGTCCATTCCGTG | This study | Cloning |
| *pYT17_sun-1_3'UTR_fwd* | CACGGAATGGACGAGCTCTACAAGGGATAAAAACGCCGTATTATTGTTCCTGC | This study | Cloning |
| *pYT17_sun-1_3'UTR_rev* | CCGAAACGCGCGAGACGAAAAAACTCTAGAGAAAACAACAGTG | This study | Cloning |
| *sun-1p_eGFP_sun-1 3'UTR_150bp_HA_fwd* | TCCGTATTTTTCCCGAATAAATATTCAGTGATTGCGAGAAGAAATTATTTTAATGATCAAACTCTAATGATTCTTGATAAAGAATGTATTGTTTTGTAAAATAAAGAAAACAAGTATTCTAGGTAACATATTAACCTGGGAACAATAAGTTCACAGCAGAGAAGGCAAAC | This study | PCR Amplification (Repair Template for CRISPR-Cas9 Mediated Insertion) |
| *sun-1p_eGFP_sun-1 3'UTR_150bp_HA_rev* | AAAGTTAAAAAATACAATTTCTTCAATCAACCAAACTTAACTAGAACATCAATAATTAATTACAATTTATAATTAAAAATCACAGTCTTTAACTATAATAACATGAAAATGACATCGCCATTATGTATTTTCAATGACATCACTTCACCGAAAAACTCTAGAGAAAACAACAGTGG | This study | PCR Amplification (Repair Template for CRISPR-Cas9 Mediated Insertion) |
| *sun-1p_eGFP_sun-1 3'UTR_fwd* | TCACAGCAGAGAAGGCAAAC | This study | PCR Amplification (Repair Template for CRISPR-Cas9 Mediated Insertion) |
| *sun-1p_eGFP_sun-1 3'UTR_rev* | AAAAACTCTAGAGAAAACAACAGTGG | This study | PCR Amplification (Repair Template for CRISPR-Cas9 Mediated Insertion) |
| *ChrIII_7007.6_fwd* | TTCAGTGATTGCGAGAAGAAATTA | This study | Genotyping |
| *ChrIII_7007.6_rev* | CTTCAATCAACCAAACTTAACTAG | This study | Genotyping |
| *3xHA_rev* | GGACATCATATGGGTAGGC | This study | Genotyping |
| *tomm-20(1-55aa)_rev* | TGCTCCAGCCTGGGCACG | This study | Genotyping |
