## Supplementary Table 6 for "Mitochondrial GTP Metabolism Regulates Reproductive Aging"

### Supplementary Table 6. tracrRNA & crRNA

| RNA name | Sequence | Source | Strains generated |
|---|---|---|---|
| *tracrRNA* | AACAGCAUAGCAAGUUAAAAUAAGGCUAGUCCGUUAUCAACUUGAAAAAGUGGCACCGAGUCGGUGCUUUUUUU | *Deltcheva et al., 2011* | N.A. |
| *dpy-10 crRNA* | GCUACCAUAGGCACCACGAG | *Arribere et al., 2014* | N.A. |
| *gfp_drp-1 crRNA* | GAACUAUACAAACACUCCAC | This study | MCW1315 |
| *sucg-1_5' crRNA* | GAGUAUCGUUGGGAGCUCGA | This study | MCW1325, MCW1331 |
| *sucg-1_3' crRNA* | CCUUCCAAACGAACAACCAU | This study | MCW1325, MCW1331 |
| *suca-1_5' crRNA* | CGGGACGCUACGACACAUUG | This study | MCW1329, MCW1330 |
| *suca-1_3' crRNA* | GCUUCGUCGAGGUUAUCGCA | This study | MCW1329, MCW1330 |
| *ChrIII_7007.6_crRNA* | UUAACCUGGGAACAAUAAGU | This study | MCW1408 |
| *gfp_3'_crRNA* | GGAAUGGACGAGCUCUACAA | This study | MCW1473 |
| *gfp_5'_crRNA* | AGAUCUACUCGGUAUGUCCA | This study | MCW1550 |
