## Supplementary Table 7 for "Mitochondrial GTP Metabolism Regulates Reproductive Aging"

### Supplementary Table 7 – Strain List

| REAGENT or RESOURCE | SOURCE | IDENTIFIER |
|---|---|---|
| Bacterial and Virus Strains | | |
| *Escherichia coli* HT115(DE3) | Caenorhabditis Genetics Center | HT115 |
| *Escherichia coli* OP50 | Caenorhabditis Genetics Center | OP50 |
| *Escherichia coli* HB101 | Caenorhabditis Genetics Center | HB101 |
| Vidal RNAi library | Open Biosystems | ORF RNAi Collection V1.1 |
| Ahringer RNAi library | Source BioScience | *C. elegans* RNAi Collection (Ahringer) |
| RNA interference competent OP50 strain *(rnc:14::DTn10; laczgA::T7pol camFRT)* | *(Neve et al., 2020)* | |
| Experimental Models: Organisms/Strains | | |
| *C. elegans* N2 Wild-type | Caenorhabditis Genetics Center | N2 (RRID:WB-STRAIN:WBStrain00000001) |
| *sucg-1(syb3617[sucg-1::eGFP]) IV* | Suny Biotech | PHX3617 |
| *suca-1(syb4685[suca-1::eGFP]) X* | Suny Biotech | PHX4685 |
| *mkcSi13[sun-1p::rde-1::sun-1 3'UTR + unc-119(+)] II; rde-1(mkc36) V* | Caenorhabditis Genetics Center | DCL569 (RRID:WB-STRAIN:WBStrain00005607) |
| *egxSi155[mex-5p::tomm-20::mKate2::pie-1 3'UTR + unc-119(+)] II; unc-119(ed3) III* | Caenorhabditis Genetics Center | EGD629 |
| *egxSi155[mex-5p::tomm-20::mKate2::pie-1 3'UTR + unc-119(+)] II; unc-119(ed3) III; sucg-1(syb3617[sucg-1::eGFP]) IV* | This paper | MCW1373 |
| *raxEx618[pie-1p::cox8(MTS)::ndk-1::3xHA::pie-1 3'UTR + myo-2p::GFP]* | This paper | MCW1581 |
| *egxSi152[mex-5p: tomm-20::gfp::pie-1 3'UTR + unc-119(+)] II; unc-119(ed3) III* | Caenorhabditis Genetics Center | EGD623 |
| *raxEx190[pie-1p::drp-1::tbb-2 3'UTR + myo-2p::GFP]* | This paper | MCW618 |
| *raxIs141[pie-1p::drp-1::tbb-2 3'UTR + myo-2p::GFP]* | This paper | MCW1220 |
| *raxIs141[pie-1p::drp-1.b::tbb-2 3'UTR + myo-2p::GFP]; egxSi152[mex5p::tomm-20::gfp::pie-1 3'UTR + unc-119(+)] II; unc-119(ed3) III* | This paper | MCW1357 |
| *drp-1(or1941[GFP::drp-1]) IV* | Caenorhabditis Genetics Center | EU2917 (RRID:WB-STRAIN:WBStrain00007414) |
| *drp-1(rax82[GFP::Degron::drp-1]) IV* | This paper | MCW1315 |
| *ieSi68[sun-1p::TIR1::mRuby::htp-1 3'UTR + Cbr-unc-119(+)] II; unc-119(ed3) III* | Caenorhabditis Genetics Center | CA1472 (RRID:WB-STRAIN:WBStrain00004073) |
| *ieSi68[sun-1p::TIR1::mRuby::htp-1 3'UTR + Cbr-unc-119(+)] II; unc-119(ed3) III; drp-1(rax82[GFP::Degron::drp-1]) IV* | This paper | MCW1326 |
| *drp-1(tm1108) IV* | Caenorhabditis Genetics Center | CU6372 (RRID:WB-STRAIN:WBStrain00005196) |
| *egxSi152[mex5p::tomm-20::GFP::pie-1 3'UTR + unc-119(+)] II; unc-119(ed3) III; drp-1(tm1108) IV* | This paper | MCW1584 |
| *sucg-1(rax83) IV* | This paper | MCW1325 |
| *sucg-1(rax86) IV* | This paper | MCW1331 |
| *suca-1(rax84) X* | This paper | MCW1329 |
| *suca-1(rax85) X* | This paper | MCW1330 |

| | | |
|---|---|---|
| *sucg-1(syb3617[sucg-1::eGFP]); sucg-1(rax83) IV* | This paper | MCW1375 |
| *sucg-1(syb3617[sucg-1::eGFP]); sucg-1(rax86) IV* | This paper | MCW1385 |
| *raxIs89[sun-1p::GFP::sun-1 3'UTR] III)* | This paper | MCW1408 |
| *raxIs98[sun-1p::GFP::3xHA::sun-1 3'UTR] III)* | This paper | MCW1473 |
| *raxIs109[sun-1p::tomm-20(1-55aa)::GFP::3xHA::sun-1 3'UTR] III* | This paper | MCW1550 |
